# Liver sinusoidal endothelial cells constitute a major route for hemoglobin clearance

**DOI:** 10.1101/2023.11.14.566925

**Authors:** Gabriela Zurawska, Zuzanna Sas, Aneta Jończy, Raghunandan Mahadeva, Patryk Slusarczyk, Marta Chwałek, Daniel Seehofer, Georg Damm, Maria Kulecka, Izabela Rumieńczyk, Morgane Moulin, Kamil Jastrzębski, Michal Mikula, Anders Etzerodt, Remigiusz Serwa, Marta Miączyńska, Tomasz P. Rygiel, Katarzyna Mleczko-Sanecka

## Abstract

Mild rupture of aged erythrocytes occurs physiologically in the spleen, leading to the release of hemoglobin (Hb), while pathological hemolysis characterizes several diseases. The detoxification of Hb has traditionally been attributed to the sequestration of Hb-haptoglobin complexes by macrophages. However, this process remains incompletely studied in animal models or primary cells, leaving the precise mechanisms of Hb clearance elusive. Using mice and primary liver cell cultures (murine and human), we uncovered that Hb uptake is chiefly performed by liver sinusoidal endothelial cells (LSECs) and involves macropinocytosis. Consistently, mouse LSECs displayed proteomic signatures indicative of active heme catabolism, ferritin iron storage, antioxidant defense, and macropinocytic capacity. LSECs also exhibited high iron content and the expression of hepcidin-regulated iron exporter ferroportin. Using erythrocyte/Hb transfusion assays in mice, we demonstrated that while splenic macrophages excel in phagocytosis of erythrocytes, LSECs primarily scavenge Hb and Kupffer cells clear erythrocyte membranes, the spleen-borne hemolysis products delivered to the liver via the portal circulation. High-dose Hb injections resulted in transient hepatic iron retention, early LSEC-specific induction of heme-catabolizing *Hmox1* and iron-sensing *Bmp6*, culminating in hepcidin-mediated temporary hypoferremia. Transcriptional induction of *Bmp6* in mice was phenocopied by erythrocyte lysis upon phenylhydrazine or iron citrate injection, although the latter elicited a distinct LSEC transcriptional signature compared to Hb. In conclusion, we identify LSECs as key Hb scavengers, a function that establishes the spleen-to-liver axis for iron recycling and contributes to heme detoxification during hemolysis, coupled with the induction of the BMP6-hepcidin axis to restore iron homeostasis.

## Introduction

Internal iron recycling from senescent red blood cells (RBCs) satisfies most of the body’s iron needs (1, 2) and relies mainly on the phagocytic clearance of aged RBCs by splenic red pulp macrophages (RPMs) (3, 4). However, recent evidence suggests that some aged RBCs escape erythrophagocytosis and lyse locally in the spleen, thus releasing hemoglobin (Hb) (5). Several inherited and acquired disorders, including hereditary anemias, autoimmune hemolytic diseases, or infections, are characterized by compromised erythrocyte stability and an increased risk of hemolysis (6, 7). Free Hb is captured by the acute-phase plasma protein haptoglobin (8). The Hb-haptoglobin complexes are sequestered via CD163 (9), a receptor that is highly expressed by both splenic RPMs and liver macrophages, Kupffer cells (KCs) (1, 5). However, the role of these macrophage populations in Hb uptake has not been well characterized. Interestingly, pharmacokinetic studies in non-rodent mammals have shown that the clearance rate of the Hb-haptoglobin complex is significantly slower than that of free Hb (10), and that Hb sequestration may occur independently of haptoglobin and/or CD163 (11, 12). These observations are clinically relevant as enhanced erythrophagocytosis and prolonged erythrolytic conditions lead to the partial loss of the CD163-expressing iron-recycling macrophages (3, 13) and are characterized by depletion of the plasma haptoglobin pool (7, 14, 15). Collectively, this evidence suggests that macrophages may be dispensable for Hb clearance. It is well established that free Hb undergoes renal glomerular filtration (16, 17). However, it remains unclear whether other specialized routes of extra-renal and macrophage-independent Hb clearance operate in the body.

The liver receives approximately 25% of the cardiac output and is exposed to blood from the portal circulation, which drains the gastrointestinal tract and the spleen (18). The hepatic capillary network, which is composed of venous sinusoids, is specialized for monitoring and filtering blood components. Liver sinusoidal endothelial cells (LSECs), along with KCs, constitute the most efficient dual scavenging system in the body (19). While KCs engulf large particles, LSECs remove macromolecules and nanoparticles, a function that protects the body from waste by-products and noxious blood factors (20, 21). The maintenance of appropriate iron homeostasis adds to the growing spectrum of homeostatic functions of LSECs. Importantly, LSECs are the sensors of body iron levels and the major producers of inducible angiokine bone morphogenetic protein 6 (BMP6), and homeostatic BMP2 (22, 23). BMPs function as upstream activators of hepcidin, a key iron-regulatory hormone produced by liver hepatocytes (2). Hepcidin suppresses iron release into the bloodstream via the sole known iron exporter ferroportin (FPN), thereby limiting iron availability under iron-rich conditions (24). However, it remains unclear how different types of iron signals are sensed and detoxified in the liver microenvironment, and whether the emerging scavenging functions of LSECs cross-talk with their role in maintaining iron homeostasis. Here, we show that LSECs constitute a major route for Hb clearance. They contribute to steady-state iron recycling from spleen-derived Hb that enters the liver via the portal vein and participate in heme detoxification during hemolysis, timely coupled with induction of the iron-sensing BMP6-hepcidin axis.

## Material and Methods

### Mice and *in vivo* procedures

Female BALB/c mice (8-10 weeks old) were obtained from the Experimental Medicine Centre of the Medical University of Bialystok or the Mossakowski Medical Research Institute of the Polish Academy of Sciences. Female C57BL/6-Tg(UBC-GFP)30Scha/J (UBI-GFP/BL6) mice were kindly provided by Aneta Suwińska (Faculty of Biology, University of Warsaw, Poland). Female and male (8-12 weeks old) WT C57BL/6J and *Cd163^-/-^* mice were kindly provided by Anders Etzerodt (Department of Biomedicine, Aarhus University, Denmark). For the dietary experiment, C57BL/6J females (4 weeks old) were obtained from the Experimental Medicine Centre of the Medical University of Bialystok. Mice were fed a standard iron diet of 200 mg/kg (control) or a low iron diet containing <6 mg/kg (iron deficient) for 5 weeks before analysis. All mice were maintained at the SPF facility under standard conditions (20°C, humidity 60%, 12-h light/dark cycle). All data reporting the uptake of Hb by LSECs were obtained using female mice or primary female LSECs. However, we expect the findings to be relevant for both sexes.

Proteins or their conjugates were dissolved in PBS and administered intravenously (*i.v.*) at doses indicated in the figure legends. For macrophage depletion, mice received an *i.v.* solution of liposomes containing clodronic acid (LIPOSOMA, #C-SUV-005) (5 ml/kg) or control empty liposomes for 24 h. A sterile aqueous iron citrate (FeCit, 150 µg/mouse) solution (Sigma-Aldrich, #F3388) or sterile citric acid buffer (0.05 M, Sigma-Aldrich, #251275) was normalized to pH 7.0 and administered *i.v.* for 5 h. Mini-hepcidin (PR73, 50 nmol/mouse) (kind gift from Elizabeta Nemeth, UCLA, USA) was injected intraperitoneally (*i.p.*) for 4 h.

To induce hemolysis, a sterile solution of phenylhydrazine (PHZ) (Sigma-Aldrich, #P26252) in PBS was administered *i.p.* at a dose of 0.125 mg/g of body weight for 6 h. PKH26-stained (Sigma Aldrich, #PKH26GL-1KT) temperature-stressed UBI-GFP RBCs were resuspended to 50% hematocrit in HBSS (Capricorn, #HBSS-1A) and administered *i.v.* for 1.5 h. PKH26-stained RBCs ghosts were mixed with fluorescently labeled Hb (10 μg/mouse) and administered *i.v.*for 1.5 h.

### Isolation of human non-parenchymal liver cells (NPCs)

Liver tissue samples were obtained from the Department of Hepatobiliary Surgery and Visceral Transplantation at Leipzig University Medical Center. Specimens were collected from macroscopically healthy tissue adjacent to resected segments in patients with primary or secondary liver tumors or benign local liver diseases. During resections, both diseased and surrounding unaffected liver tissue were removed; a portion of this unaffected tissue was used for liver cell isolation. All patients provided informed consent for tissue use in research, following Leipzig University Hospital ethical guidelines (approval numbers 322/17-ek and 057/23-ek). Detailed protocols for cell isolation and further examination are provided in Supplementary Methods.

### Cell-based and biochemical assays

Isolation of murine and human NPCs for primary cell cultures, cell treatments, and immunofluorescence are described in Supplementary Methods. Isolation of cells from the spleen, heart, bone marrow, and aorta, flow cytometry, FACS-sorting, liver tissue immunofluorescence, heme/iron measurements, preparation of conjugated mouse Hb, stressed RBCs, and RBC ghosts, and real-time quantitative PCR (RT-qPCR) were performed as previously reported (25), or/and described in detail in Supplementary Methods.

### Statistics

Data are represented as the mean ± SEM. The number of mice/samples per group or the number of independent cell-based experiments are shown in the figures. Statistical analyses were conducted using GraphPad Prism software (Version 9.4.1). When two groups were compared two-tailed unpaired Welch’s t-test was applied, whereas for more groups, the One-Way or Two-Way Analysis of Variance (ANOVA) with post-hoc Tukey’s test for multiple comparisons was performed, as indicated in the figure legends. For all the experiments, α=0.05. Statistically significant changes between means are indicated as follows: ns, not significant at p>0.05, * for p<0.05, ** for p<0.01, *** for p<0.001, and **** for p<0.0001.

### Ethical Approval

All animal experiments were conducted following the guidelines of European Directive 2010/63/EU and the Federation for Laboratory Animal Science Associations. The procedures were approved by the Local Ethics Committee in Warsaw (No. WAW2/150/2019, WAW2/138/2019, WAW2/137/2020, WAW2/137/2021, WAW2/179/2021, and WAW2/094/2021), or for the dietary experiment by the local ethical committee in Olsztyn No. 26/2018. For the use of human cells, this study adhered to the Declaration of Helsinki and received approval from the Ethics Committee of the Faculty of Medicine at Leipzig University (Biobank of Surgical Research, registration no. 322/17-ek, 2020/06/10, ratified on 2021/11/30; Role of human LSEC in hemoglobin clearance, registration no. 057/23-ek, 2023/08/21).

### Data availability

Transcriptome analysis of LSECs was conducted using the AmpliSeq method and proteomics was performed by label-free quantification with mass spectrometry as described in the Supplementary Methods. LSEC transcriptomic data for iron citrate injection have been previously deposited in the GEO repository under accession no. GSE235976, whereas data for Hb injection are accessible under no. GSE240270 (with a token: uhkrgywyhledjmj to allow for review). Proteomics data were deposited to the ProteomeXchange Consortium via the PRIDE partner repository with the dataset identifier PXD051274 (username; password: 0QJOAbzS for review).

## Results

### LSECs engage macropinocytosis to sequester Hb

Studies using radiolabeled Hb have shown that the clearance of injected hemoglobin is rapid (16) and mainly mediated by the liver, kidney, and spleen (16, 17). To address the involvement of macrophages in Hb uptake, we first performed experiments using clodronate liposomes, a well-established strategy for depleting tissue macrophages, including hepatic KCs. Using a whole-organ imaging system, we observed that regardless of the presence of KCs, the liver emerged as the major organ sequestering fluorescently-labeled mouse Hb (Fig. 1A and S1A and B). Next, using flow cytometry and confocal microscopy imaging of mouse liver sections, we identified CD146^+^STAB2^+^ LSECs as the major hepatic cell type that takes up Hb, surpassing KCs, hepatocytes, and monocytes (Fig. 1B and C, Fig. S1C; see Data Supplements for gating strategies). We found that murine primary LSEC cultures recapitulated the high capacity of this cell type for Hb uptake, as validated by confocal microscopy and flow cytometry (Fig. 1D and Fig. S1D). We further confirmed that freshly isolated human primary LSECs, distinguished by CD36 and CD32B markers, as previously reported (26, 27) and confirmed by the Human Liver Cell Atlas (28) (Fig. S1E), outperformed other NPCs in human Hb uptake (Fig. 1E and F). Notably, flow cytometric analysis revealed that cells with high CD36 and CD23B expression, albeit not highly abundant, showed the highest capacity for Hb scavenging (Fig. 1E).

**Figure 1.**
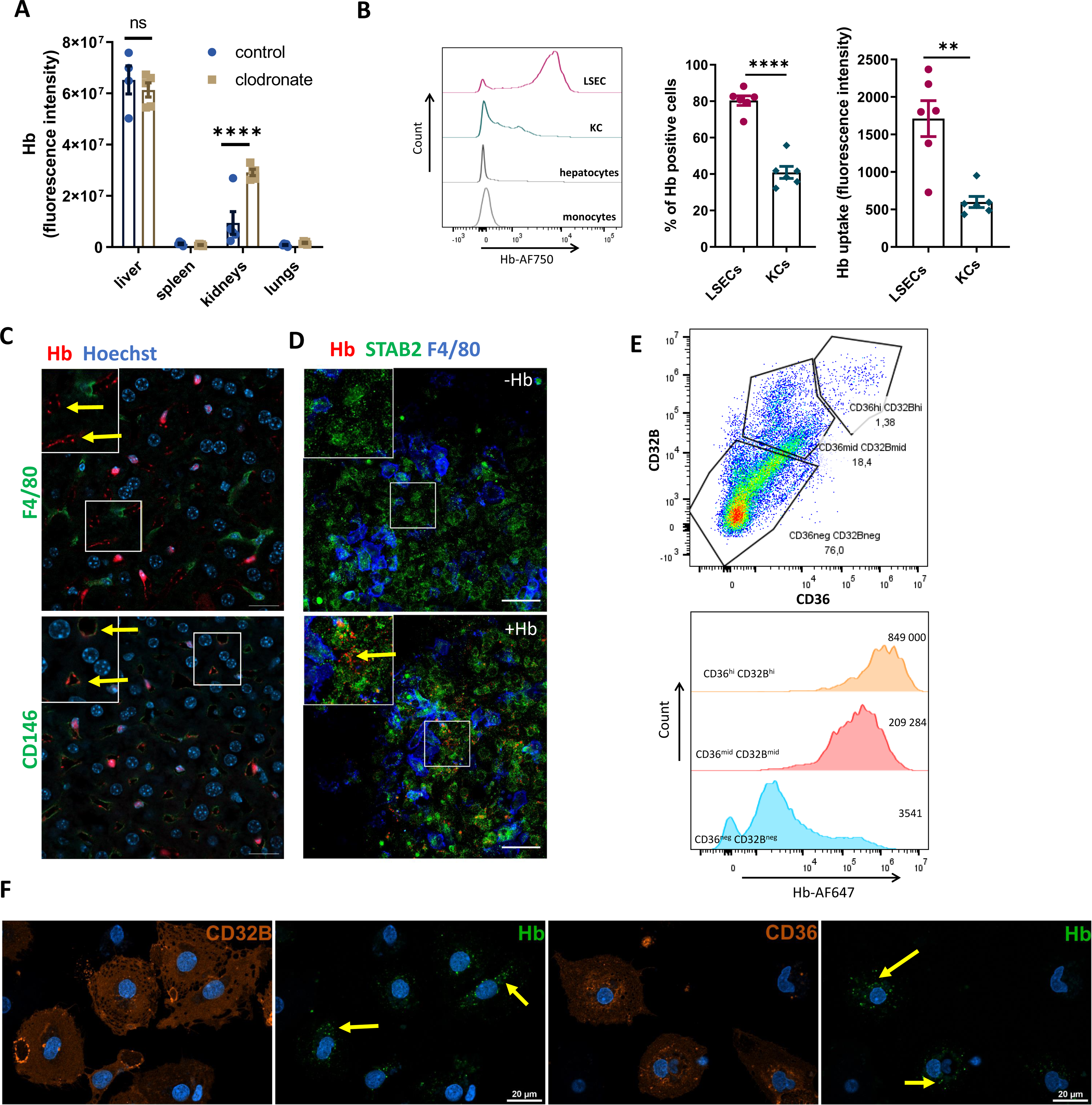
LSECs represent the major cell type that sequesters Hb. (A) Hemoglobin (Hb) distribution in control and macrophage-depleted mice (clodronate) injected with Alexa Fluor 750-labeled Hb (Hb-AF750, 10 µg/mouse), imaged with Bruker *in vivo* Imaging System. Results are presented as total fluorescent counts from the isolated organs. (B) AF750 fluorescence from indicated liver cell populations isolated from Hb-AF750-injected mice was analyzed with flow cytometry. (C) Frozen liver slices from mice injected with Hb-AF647 (red) were processed and stained for Kupffer cells (KCs) (F4/80, green) or LSECs (CD146, green) and nuclei (blue). (D) Murine liver primary non-parenchymal cells (NPCs) were treated *in vitro* with Hb-AF750 (red) (0.5 µg/ml) and stained for F4/80 (blue) and the LSEC marker STAB2 (green). (C, D) Areas in the highlighted rectangles are shown at 5x higher magnification. (E) Flow cytometric analysis of human freshly isolated primary NPCs, exposed in culture to human Hb-AF647 and stained with the human LSEC markers CD36 and CD32B. The intensity of Hb-A647 uptake was measured in two LSEC subpopulations, more pure CD36^hi^CD32B^hi^ and CD36^mid^CD32B^mid^, as compared to other NPCs CD36^neg^CD32B^neg^. Numbers indicate MFI. Data are representative of three independent donors of human NPCs. (F) Hb-AF647 (green) uptake was imaged in freshly isolated human liver NPCs stained for the LSEC markers CD36 and CD32B (orange). (C, D, F) Arrows indicate Hb presence in LSECs; Scale bars, 20 μm. Numerical data are expressed as mean ± SEM, each data point represents one biological replicate. Two-way ANOVA with Tukey’s Multiple Comparison tests was used in A, and Welch’s unpaired t-test was used to determine statistical significance in B; ns - not significant, **p<0.01 and ****p<0.0001.

To better understand the mechanism of Hb uptake we used mouse primary liver cell cultures. In contrast to murine KCs, LSECs were negative for the Hb-haptoglobin complex receptor, CD163 (Fig. 2A), and took up free Hb more robustly than haptoglobin-bound Hb (Hb:Hp; Fig. 2B). Therefore, we hypothesized that LSECs may sequester Hb in a receptor-independent manner, possibly via macropinocytosis, which is a non-specific internalization of extracellular fluid (29). Staining of primary murine LSECs with an early endosome marker, EEA1, revealed many constitutively formed intracellular vesicles of macropinosome size (1-2 μm) (Fig. 2C and S2A). Fluorescently-labeled Hb was entrapped, though not exclusively, within such vesicles (Fig. 2C and S2A), and co-localized with high-molecular-weight dextran, a known macropinocytic cargo (Fig. 2D and S2B) (30, 31). Human primary CD36-high LSECs likewise displayed numerous macropinosome-like vesicles and efficiently sequestered dextran (Fig. 2E and F). Consistently, ethyl-isopropyl amiloride (EIPA), a well-established blocker of macropinocytosis, in contrast to an inhibitor of clathrin-mediated endocytosis, chlorpromazine (CPZ), abolished Hb uptake in murine primary LSECs (Fig. 2G and S2C) and decreased Hb scavenging by their human counterparts (Fig. 2H). Likewise, Hb internalization by murine LSECs was partially dependent on actin remodeling (suppressed by latrunculin A, LAT-A), and Cdc42 activity (inhibited by ML141), both of which are important for macropinosome formation (30). Hb uptake also required canonical Wnt signaling via β-catenin (blocked by PRI-724), a pathway that is active in LSECs (32) and has previously been shown to activate macropinocytosis (Fig. 2G) (33, 34). Similarly, Hb uptake by human primary LSECs was suppressed by LAT-A (Fig. 2H). Noteworthy, Hb uptake was not altered by the micropinocytosis blocker, nystatin (Fig. 2G). Interestingly, close to 100% of LSECs actively sequestered Hb even at low doses and the capacity for the uptake did not show saturation with higher cargo presence, implying a receptor-independent entry route (Fig. S2D). However, the same doses of dextran, a cargo of similar molecular weight, were internalized by LSECs with far lower efficiency (Fig. S2D). This implied a certain specialization of LSECs towards Hb scavenging, even though the uptake of dextran by LSECs was likewise abolished by the macropinocytosis inhibitor EIPA or actin remodeling blocker (Fig. S2E). Confirming the high macropinocytic capacity of LSECs *in vivo*, we demonstrated their ability for efficient uptake of fluorescently-labeled albumin, another known macropinocytic cargo (Fig. 2I) (31). Of note, we observed that the uptake of Hb by primary murine KCs was not affected by the clathrin endocytosis inhibitor CPZ, and appeared to be dependent on macropinocytosis (Fig. S2F). Interestingly, the uptake of the Hb:Hp complex by LSECs and KCs was suppressed by both EIPA and CPZ, implying the involvement of both clathrin-dependent endocytosis and macropinocytosis routes in this process (Fig. S2G). Taken together, our data identify LSECs as efficient scavengers of both free and haptoglobin-bound Hb and implicate macropinocytosis as an important alternative pathway for Hb uptake, which also extends to KC-mediated Hb scavenging.

**Figure 2.**
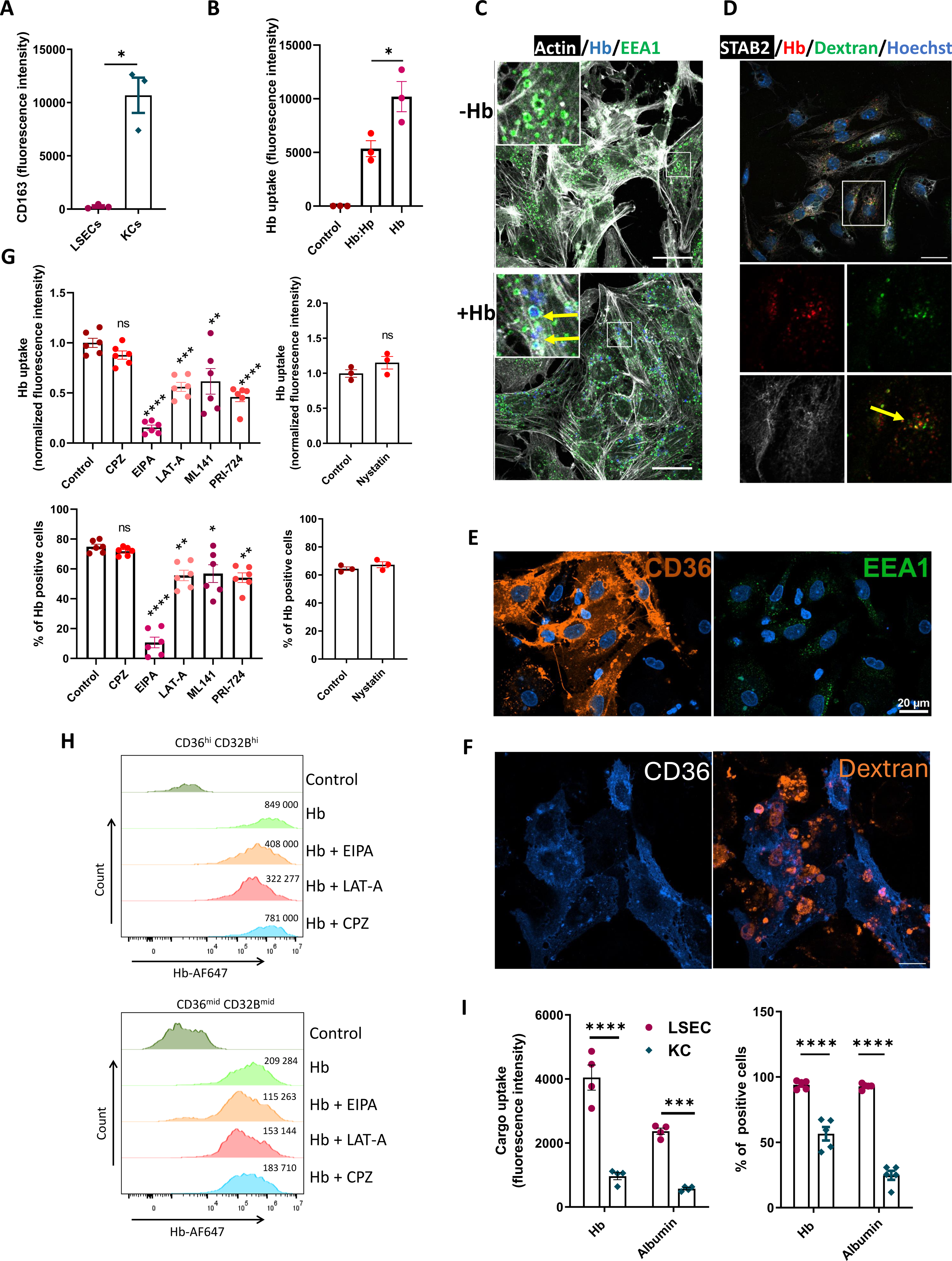
LSECs engage macropinocytosis to sequester Hb. (A) Cell-surface expression of the CD163 receptor on LSECs and KCs was measured by flow cytometry and presented as fluorescence intensity. (B) Uptake of free Hb and Hb in the presence of haptoglobin (Hp, Hb:Hp) in primary murine LSEC cultures was measured with flow cytometry. (C) Hb-AF647 vesicle localization was imaged with microscopy in NPCs *in vitro* cultures depleted from macrophages. Arrows indicate Hb-AF647 (blue) presence in the EEA1^+^ (green) and Actin^+^ (white) vesicles. Areas in the highlighted rectangles are shown at higher 5x magnification (left corner). Scale bars, 20 μm. (D) Co-localization of Hb-AF647 (red) with rhodamine-dextran (green) was imaged with microscopy in STAB2^+^ LSECs (white). Arrow indicates colocalization of Hb and rhodamine-dextran. The area in the highlighted rectangle is shown at higher 5x magnification in separate channels below. Scale bar, 20 μm. EEA1^+^ vesicles (E) and FITC-dextran uptake (blue) (F) were imaged with microscopy in human NPCs cultures stained with the LSEC marker CD36. (G) NPCs were pre-treated with the clathrin-mediated endocytosis inhibitor chlorpromazine (CPZ), the macropinocytosis blocker EIPA, actin polymerization blocker latrunculin A (LAT-A), inhibitor of Cdc42 GTPase ML141, β-catenin inhibitor PRI-724 and with micropinocytosis blocker nystatin before Hb-AF750 treatment for 10 min. Flow cytometry was used to determine Hb uptake by the LSECs. (H) Flow cytometric analysis of CD36^hi^CD32B^hi^ and CD36^mid^CD32B^mid^ human primary LSECs, treated with Hb-AF647 and exposed to EIPA, LAT-1 and CPZ. Numbers indicate MFI. Data are representative of three independent donors of human NPCs. (I) Mice were injected for 1 h with Hb-AF750 or albumin-AF750, both at 10 µg/mouse. The fluorescence intensity of AF750, the percentage of AF750^+^ LSECs and KCs populations were measured with flow cytometry. (C-F) Scale bars, 20 μm. Numerical data are expressed as mean ± SEM, each data point represents one biological replicate. Welch’s unpaired t-test was used to determine statistical significance in A and G for nystatin, two-way ANOVA with Tukey’s Multiple Comparison tests was used in I; while one-way ANOVA with Tukey’s Multiple Comparison tests was used in B and G. ns - not significant; *p<0.05, **p<0.01, ***p<0.001 and ****p<0.0001.

### LSECs outperform other endothelial and macrophage populations in Hb uptake

Similarly to the liver, sinusoidal endothelial cells are also present in the spleen and bone marrow (20), organs that perform critical functions in systemic iron metabolism owing to the presence of CD163^+^ iron-recycling RPMs and erythroblastic island macrophages, respectively. Therefore, we sought to accurately compare the Hb-scavenging capacity of endothelial and macrophage populations from the spleen and bone marrow with those of hepatic LSECs and KCs over a wide range of Hb doses. We used 1 μg of Hb, which is below the binding capacity of circulating haptoglobin (10-20 μg/ml plasma), an intermediate dose of 10 μg, and 100 μg (injected together with 10 mg of unlabeled Hb), which saturates the haptoglobin pool and mimics hemolytic conditions. We also injected mice with 1 μg of Hb bound to haptoglobin. Myeloid cells and ECs from the bone marrow were not capable of efficient Hb sequestration (Fig. S3A). Strikingly, we observed the extraordinary ability of LSECs to internalize Hb as compared to the other cell types analyzed as exemplified by the 70-99% of Hb-positive cells depending on the dose (Fig. 3A) and the highest mean intensity of the Hb signal per cell as compared to KCs, splenic ECs, or RPMs (Fig. 3B, D, F, H). The scavenging capacity of KCs and splenic ECs gradually increased with the Hb dose (Fig. 3C-F), whereas RPMs contributed, albeit slightly, to Hb clearance only at the highest dose, mimicking hemolysis (Fig. 3G and H). Finally, using flow cytometry and whole-organ imaging, we did not detect Hb accumulation in the aorta, lined by non-sinusoidal ECs (Fig. S3B and C). In conclusion, our data suggest that LSECs outperform other cell types for Hb clearance over a wide range of circulating Hb concentrations.

**Figure 3.**
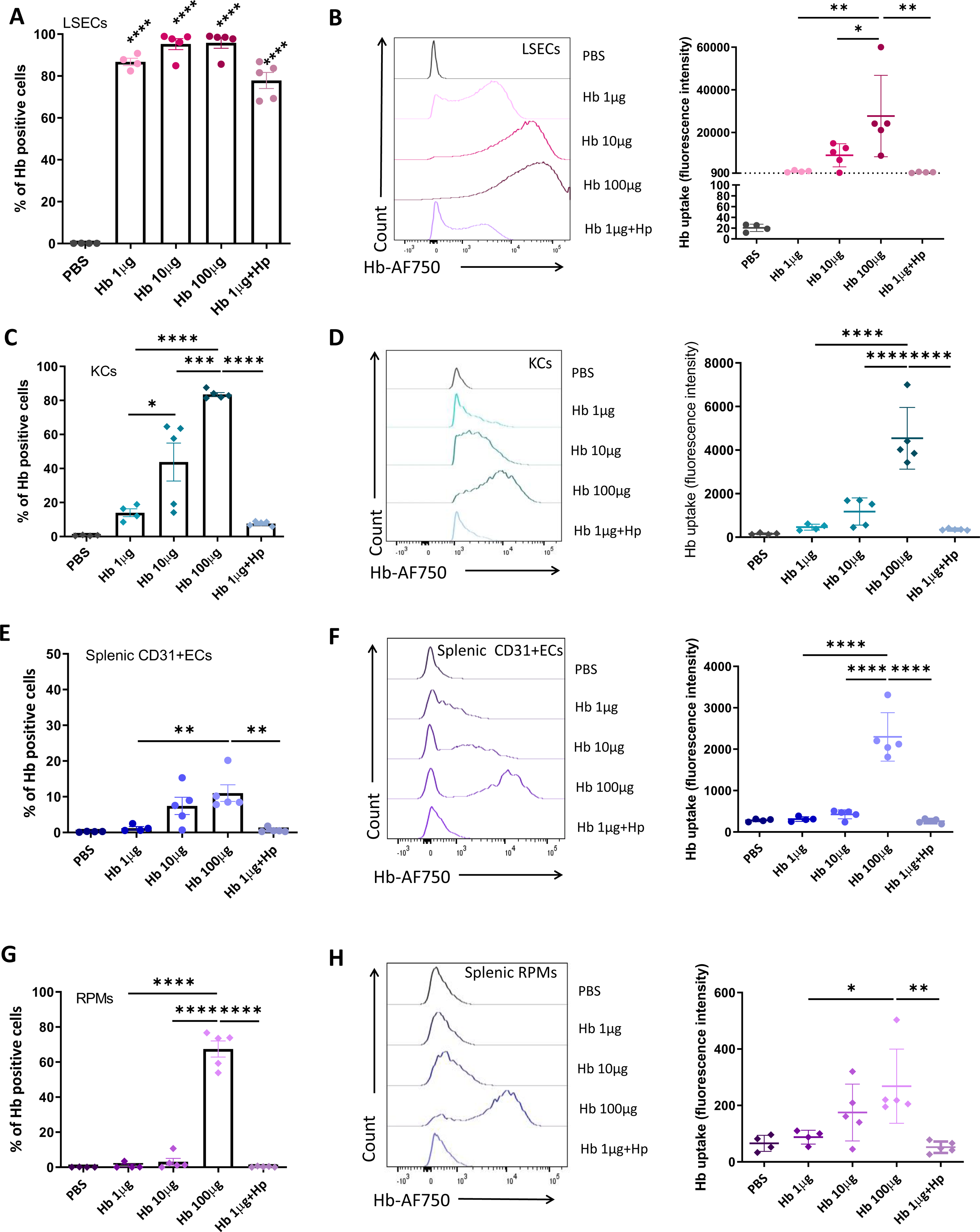
LSECs outperform other endothelial populations and macrophages in Hb uptake. Percentage of Hb-AF750 uptake by mouse (A) LSECs, (C) KCs, (E) splenic endothelial cells (ECs), and (G) RPMs. Fluorescence intensity of Hb-AF750 in (B) LSECs, (D) KCs, (F) splenic ECs, and (H) RPMs. Both parameters were analyzed with flow cytometry. Data are expressed as mean ± SEM and each data point represents one biological replicate. One-way ANOVA with Tukey’s Multiple Comparison tests was used to determine statistical significance; ns - not significant, *p<0.05, **p<0.01, ***p<0.001 and ****p<0.0001

### LSECs constitutively express proteins critical for iron recycling from heme

We were intrigued by the fact that LSECs efficiently scavenged low doses of Hb. To further explore their specialization, we examined data from single-cell RNA sequencing of ECs from different mouse organs (35). Interestingly, both *Hmox1* [encoding heme oxygenase 1 (HO-1)] and *Blrvb* (encoding biliverdin reductase b, BLRVB), which are important for heme catabolism, together with important antioxidant enzymes such as *Gclc*, *Gpx4,* and *Nqo1*, were identified by Kalucka et al. (35) as metabolic markers exclusively specific for liver EC. Using the EC Atlas, we visualized the expression levels of *Hmox1* and *Blvrb,* as well as *Slc40a1*, the gene encoding the iron exporter FPN, and found a clear signature for the high expression of these transcripts in liver ECs compared to other organs (Fig. S4A). Consistently, by employing dedicated single-cell RNA-seq databases we observed high expression levels of *Slc40a1*, *Hmox1* and *Hmox2* in pig liver ECs and *Slc40a1* in human LSECs (Fig. S4B and C) (28, 36). Following up on these transcriptomic signatures, we detected protein expression of FPN and HO-1 in mouse liver sections (Fig. 4A) and primary murine LSECs (Fig. 4B). Flow cytometry confirmed clear expression of cell-surface FPN on LSECs, reaching approximately 40% of cells, albeit at lower levels compared to KCs (Fig. 4C). Furthermore, close to 100% of LSECs were positive for HO-1, although the expression levels appeared lower than those in KCs (Fig. 4D). Next, to obtain comprehensive information on LSEC adaptation to Hb clearance, we performed label-free proteomic profiling of FACS-sorted LSECs in comparison with iron recycling macrophages (RPMs and KCs) and splenic and heart ECs. This analysis identified 418 protein groups that were significantly more abundant in LSECs than in splenic and cardiac ECs (Fig. 4E, Table S1). In addition to well-established LSEC markers (STAB1, STAB2, CD206), intriguingly, these included L and H ferritin, HO-1 and BLVRB, heme binding protein HEBP1, and enzymes involved in the antioxidant response (e.g., NQO1, GCLC or PRDX1). Interestingly, the hits also included several known macropinocytosis effectors, such as Rabankirin-5, ATP6V0A1, RAB5, or septin 2 (37–40), and several putative players identified by previous RNAi screens (38) (Table S1). This proteomic signature of LSEC, implying their active steady-state involvement in Hb uptake and iron recycling from heme, was consistent with their relatively high labile iron levels compared to other ECs (Fig. 4F). FPN was robustly induced in LSECs under conditions of systemic iron deficiency, mimicking the response observed in KCs (Fig. 4G), suggesting a tight control of LSECs’ FPN by circulating hepcidin. Confirming the iron export function of LSEC FPN, we detected a significant increase in labile iron in LSECs in response to mini-hepcidin PR73 injection, a response that was even more pronounced than in KCs (Fig. 4H) (41). Collectively, these data indicate that LSECs are equipped with protein machinery that drives iron recycling from heme, suggesting their role in steady-state iron turnover.

**Figure 4.**
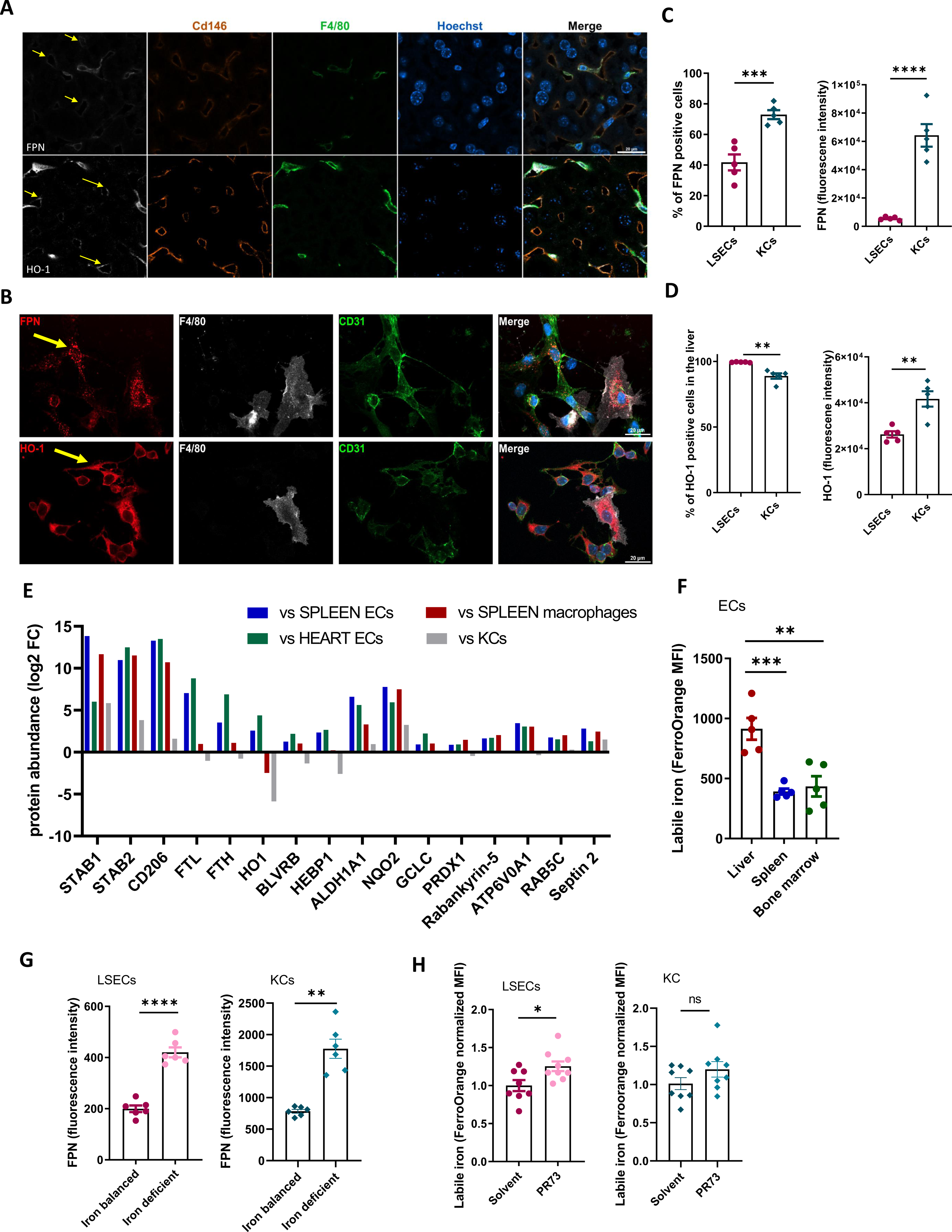
LSECs constitutively express proteins critical for iron recycling and macropinocytosis. (A) Frozen mouse liver slices were processed and stained for FPN and HO-1 (white) in KCs (F4/80, green) or LSECs (CD146, orange). (B) FPN and HO-1 presence in F4/80^+^ (white) KCs and CD31+ endothelial cells (green) imaged by microscopy in murine NPC *in vitro* cultures. (A, B) Nuclei (blue), scale bars, 20 μm. Arrows indicate ECs positive for FPN or HO-1. (C) Flow cytometry analysis of FPN expression in LSECs and KCs. (D) Flow cytometry analysis of HO-1 expression in LSECs and KCs. (E) Shown are selected proteins statistically significantly (p<0.01) more abundant in mouse FACS-sorted LSECs than in spleen and heart endothelial cells (ECs). Log2 fold change (from 3 independent cell isolates) in protein abundance in LSECs versus indicated cell type is presented. Individual label-free quantification values of protein abundance and calculated p-values are included as Data Supplements. (F) Cytosolic ferrous iron (Fe^2+^) content in endothelial cells was measured with a FerroOrange probe. (G) Flow cytometry analysis of FPN expression in LSEC and KCs isolated from mice maintained on balanced or iron-deficient diets. (H) Cytosolic ferrous iron (Fe^2+^) content was measured with a FerroOrange probe in LSECs and KCs derived from mice injected with mini-hepcidin (PR73). Numerical data are expressed as mean ±SEM and each data point represents one biological replicate. Welch’s unpaired t-test was used to determine statistical significance in C, D, G, and H; one-way ANOVA with Tukey’s Multiple Comparison tests was used in F; ns - not significant, *p<0.05, **p<0.01, ***p<0.001 and ****p<0.0001.

### LSECs and KCs support iron recycling by removing hemolysis products from the spleen

Next, we sought to understand how LSECs may be involved in physiological iron recycling. The hemolysis-driven iron recycling model proposed by Klei et al. implied but did not formally demonstrate that RPMs likely sequester spleen-derived Hb via highly expressed CD163 (5). However, CD163 knock-out mice did not show major differences in systemic and splenic iron parameters (Fig. S5). Inspired by the high capacity of LSECs for Hb scavenging, we hypothesized that hepatic cells may play a role in the clearance of splenic hemolytic products delivered via portal circulation. Therefore, we extended previous studies by quantifying the contributions of splenic and hepatic myeloid and endothelial cells to the uptake of intact RBCs, RBCs devoid of cytoplasm (RBC ghosts), and free Hb. To this end, we injected mice with temperature-stressed RBCs, derived from Ubi-GFP transgenic mice and stained with the membrane label PKH26 (Fig. 5A and B). This approach revealed that RPMs outperformed other splenic cell types, F4/80^high^ CD11b^high^ pre-RPMs, CD11c^+^ dendritic cells (DCs), monocytes, and splenic ECs in the sequestration of PKH26^+^GFP^+^ intact RBCs. An equal percentage of RPMs (approximately 30% of the population) was effective in removing PKH26^+^GFP^-^ RBC ghosts, a function that was efficiently supported by splenic pre-RPMs and, to a lesser extent, by DCs, monocytes, or ECs. Interestingly, we found that in the liver phagocytosis of intact RBCs by KCs was less efficient than the uptake of RBC ghosts (Fig. 5B). No erythrophagocytosis of intact RBCs was detected in hepatic DCs and LSECs, but they showed some capacity for the sequestration of RBC membranes, albeit lower than that of KCs. Independently, injection of fluorescently labeled RBC ghosts along with free Hb confirmed that KCs and LSECs, rather than RPMs or splenic ECs, respectively, are specialized in the clearance of these two major hemolysis products (Fig. 5C and D). Corroborating these data, we detected an increase in heme levels in the portal vein plasma and we identified hemoglobin α (HBA) and β (HBB) chains as top proteins upregulated in LSECs at the proteome-wide level after injection of stressed RBCs compared to control PBS-injected mice (Fig. 5E and F, Table S2). Interestingly, among the five proteins that were significantly increased along with HBA and HBB in LSECs following RBC transfusion, we detected two dehydrogenases ALDH1A1 and ALDH1L1 (Fig. 5F), the former with established roles in the detoxification of aldehydes formed by lipid peroxidation (42).

**Figure 5.**
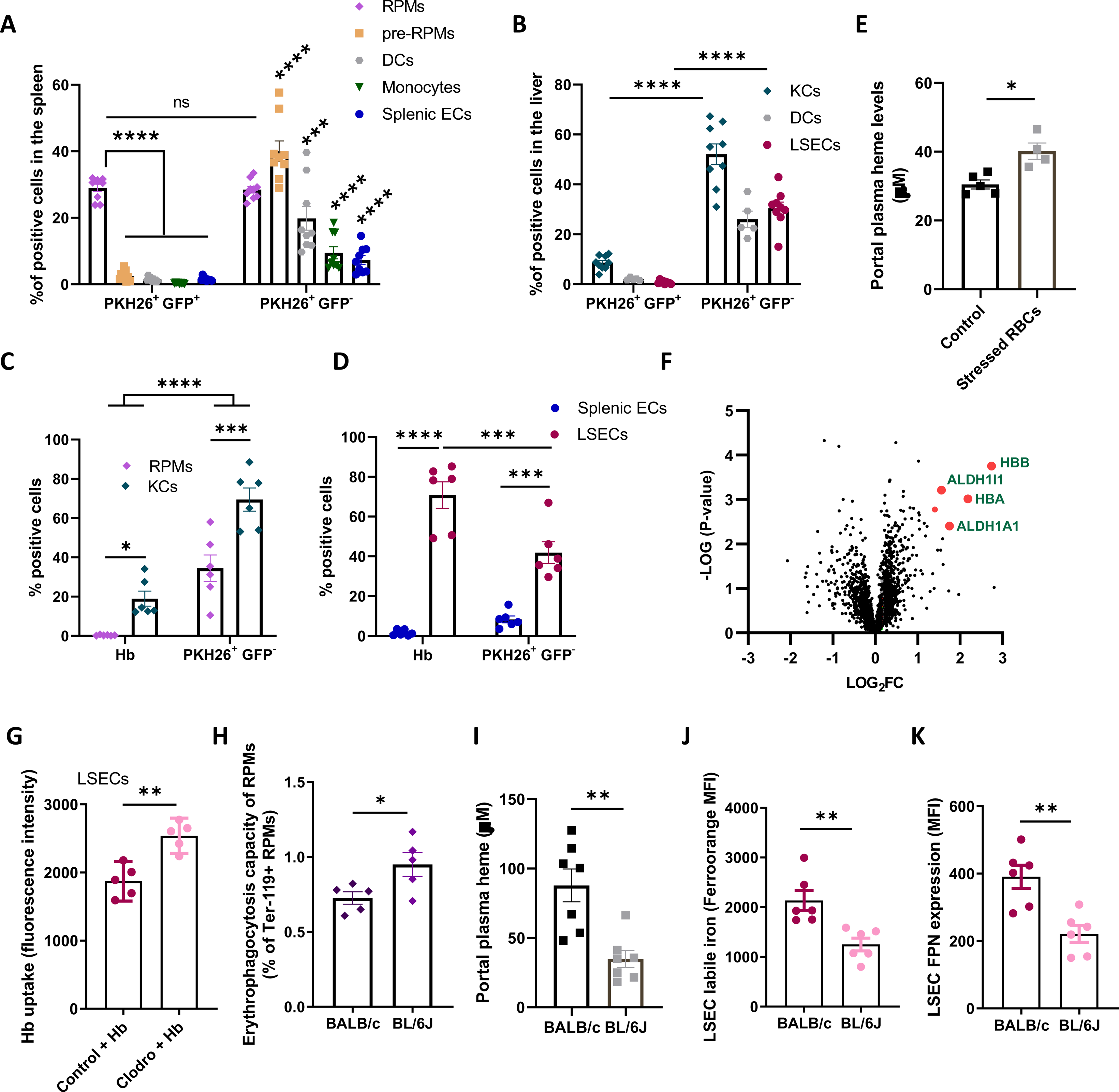
LSECs and KCs support iron recycling by removing hemolysis products from the spleen. (A-D) Mice were administered i.v. with (A-B) stressed GFP^+^ RBCs stained with a membrane marker PKH26 or (C-D) Hb-AF750 and PKH26^+^ RBCs devoid of cytoplasm (RBC ghosts). Splenic and liver cells were isolated, stained, and analyzed by flow cytometry. (A) The percentage of cells positive for markers of intact RBCs (PKH26^+^GFP^+^) and RBC ghosts (PKH26^+^GFP^-^) in splenic RPMs, pre-RPMs, dendritic cells (DCs), monocytes, and endothelial cells (ECs). (B) The percentage of PKH26^+^GFP^+^ and PKH26^+^GFP^-^ cells in liver KCs, dendritic cells (DCs), and LSECs. (C-D) The percentage of cells positive for Hb (AF750) and RBC ghosts. (E) Heme levels from the portal vein plasma of control mice and mice transfused with temperature-stressed RBCs, measured by Heme Assay Kit. (F) Volcano plot illustrating the changes in the proteome of FACS-sorted LSECs 90 min after i.v. transfusion of stressed RBCs. Proteins significantly upregulated are marked in red. (G) Hb-AF750 uptake by LSECs derived from control and macrophage-depleted mice (Clodro). (H-K) BALB/c and C57BL/6J (BL/6J) mice phenotype comparison. (H) The capacity of endogenous erythrophagocytosis was assessed by intracellular staining of the erythrocytic marker (Ter-119) in RPMs. (I) Heme levels in the portal vein were measured with Heme Assay Kit. (J) Cytosolic ferrous iron (Fe^2+^) levels in LSECs were measured with a FerroOrange probe. (K) FPN levels in LSECs measured by flow cytometry. Data are expressed as mean ± SEM and each data point represents one biological replicate. Welch’s unpaired t-test was used to determine statistical significance in E, G-K; while two-way ANOVA with Tukey’s Multiple Comparison tests was used in A, B, C, and D; ns - not significant, *p<0.05, **p<0.01, ***p<0.001 and ****p<0.0001.

Next, we sought to determine whether the ability of LSECs to sequester Hb could be modulated by the altered capacity of splenic RPMs to fully execute erythrophagocytosis. First, we observed that LSECs were more effective at Hb uptake in response to macrophage depletion after clodronate injection (Fig. 5G). Second, we took advantage of the physiological difference in iron parameters between the BALB/c and C57BL/6J (BL/6J) mice. We observed a lower erythrophagocytic capacity of RPMs from BALB/c mice compared to BL/6J mice (Fig. 5H), which was likely underlain by higher iron levels and reduced RPM FPN expression (Fig. S6A and B), as previously reported (25). This led to heme accumulation in the extracellular space of the spleen and portal plasma (Fig. 5I, S6C and S6D). This phenotype was coupled with elevated levels of labile iron and higher FPN expression in the LSECs of BALB/c mice (Fig. 5J and K), indicating their enhanced activity in iron recycling from Hb. Taken together, our findings support the physiological role of LSECs in the sequestration of endogenous spleen-derived Hb, thereby establishing a spleen-liver axis for effective iron recycling from senescent RBCs.

### LSECs detoxify hemoglobin upon hemolysis and trigger the iron-sensing BMP6 angiokine

We next investigated the response of LSECs to hemolytic conditions. To this end, we first injected mice with 10 mg of mouse Hb (equivalent to the RBC fraction in approximately 100 μl of blood) in a time-dependent manner. We detected a significant increase in heme iron content in the liver at the early time point which was normalized by the 24-hour endpoint (Fig. 6A). Consistently, we observed a transient increase in iron levels in the liver, but not in the spleen (Fig. 6B and S7A). Notably, the kidney also accumulated iron that could not be mobilized and remained elevated throughout the experiment (Fig. S7B). Hemoglobin clearance in the liver resulted in a strong transcriptional induction of the heme catabolizing enzyme *Hmox1*, a response that could be attributed specifically to LSECs (Fig. 6C and D).

**Figure 6.**
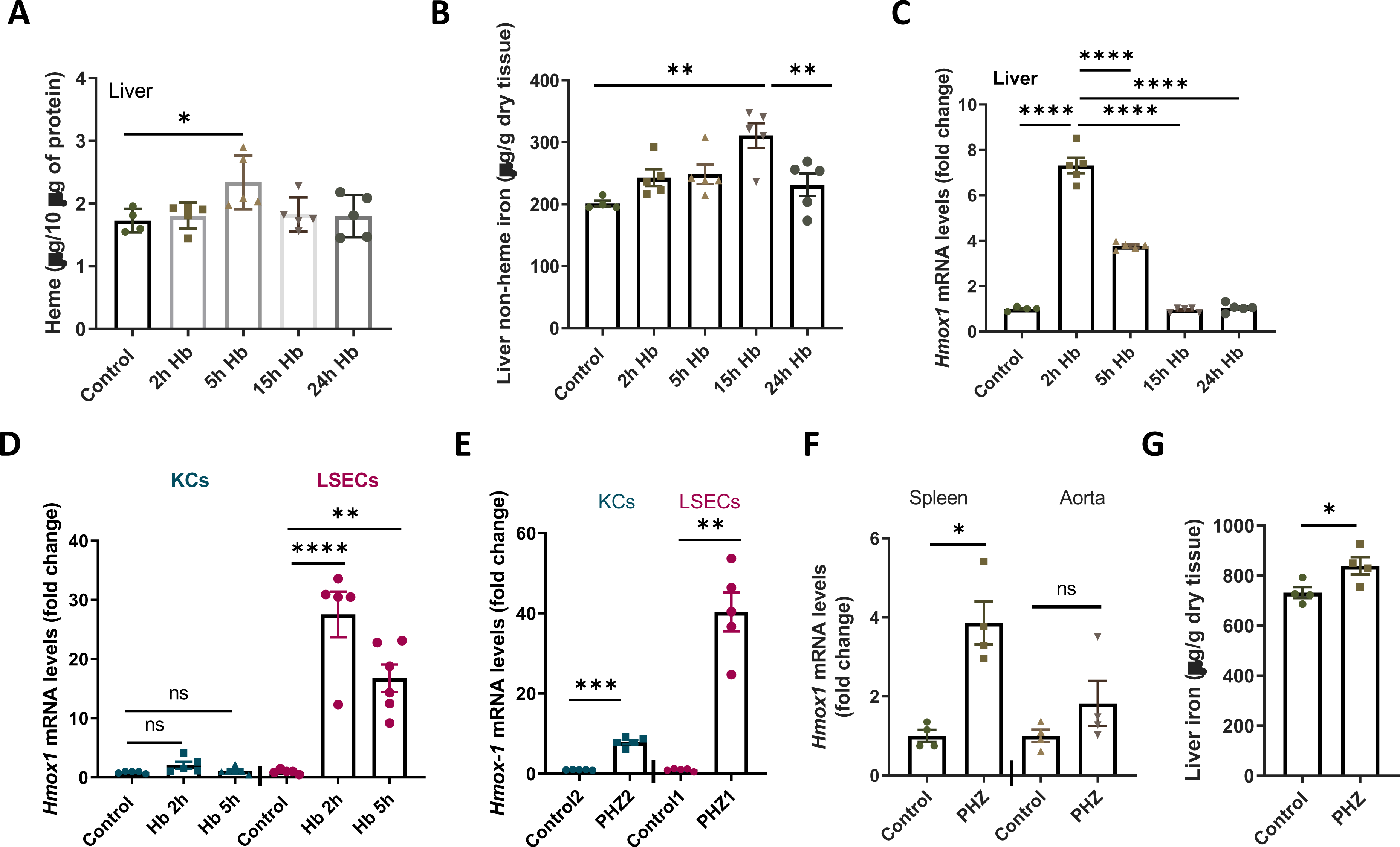
LSECs detoxify hemoglobin upon hemolysis. (A-D) Mice were injected with native hemoglobin (Hb, 10 mg/mouse) and analyzed at indicated time points. (A) Heme iron levels in the liver were determined with Heme Assy Kit and normalized to protein content. (B) Non-heme iron content in the liver was measured by bathophenanthroline colorimetric assay. (C and D) *Hmox1* gene expression measurement by RT-PCR in (C) the liver and (D) FACS-sorted KC and LSEC populations. (E-G) Hemolysis was induced in mice by i.p. injection of phenylhydrazine (PHZ, 0.125 mg/g) 6 h before analysis. *Hmox1* gene expression in (E) FACS-sorted cell populations of KCs and LSECs, and (F) spleen and aorta. (G) Non-heme iron content in the liver. Data are expressed as mean ± SEM and each data point represents one biological replicate. One-way ANOVA with Tukey’s Multiple Comparison tests was used in A-C, and separately for KCs and LSECs in D, while Welch’s unpaired t-test was used to determine statistical significance in E-G, separately for each cell type/organ; ns - not significant, *p<0.05, **p<0.01, ***p<0.001 and ****p<0.0001.

The rapid induction of *Hmox1* in LSECs upon intravenous Hb delivery was phenocopied by injecting the mice with the hemolytic agent phenylhydrazine (PHZ), which far exceeded the response of KCs, splenic cells, and the aorta (Fig. 6E and F). PHZ also caused significant, albeit mild, iron accumulation in the liver (Fig. 6G), but not in other organs (Fig. S7C and D). Hemoglobin challenge rapidly increased the hepatic expression of the iron-sensing gene *Bmp6*, attributable to FACS-sorted LSECs, accompanied by transient upregulation of the BMP target gene hepcidin (*Hamp*) in the liver (Fig. 7A-C). As expected, induction of the BMP6-hepcidin axis caused serum hypoferremia at the 15-h time-point which normalized after 24 h (Fig. 7D), reflecting the changes in liver iron content (Fig. 6B). PHZ-induced hemolysis likewise led to the activation of *Bmp6* transcription in sorted LSECs (Fig. 7E). Finally, we aimed to determine whether LSECs induce a specific response when exposed to Hb compared with non-heme iron. To this end, we performed RNA sequencing to assess global gene expression signatures upon injection of 10 mg of Hb and compared these to a previously reported response to a quantitatively matched dose of iron citrate (43). Both stimuli induced *Bmp6* at very comparable levels, and triggered an ETS1-controlled transcriptional program, as previously reported for iron citrate response (43) (Fig. S8). However, the two iron sources elicited different responses of LSECs at the transcriptome-wide level (Fig. 7F). While only 64 genes, mainly attributable to the response to oxidative stress and iron were co-induced by Hb and iron citrate, as many as 372 and 202 genes increased their expression specifically upon Hb and iron citrate injection, respectively (Fig. 7F). These differences were reflected by a distinct functional enrichment within the Hb- and iron-induced transcriptional signature (Fig. 7G). Interestingly, Hb-exposed LSECs specifically induced genes associated with immune and inflammatory responses, such as the phagocyte marker *Cd68*, the chemokines *Ccl24*, *Ccl6,* and *Ccl9*, and importantly, *Il18*, a cytokine implicated in the pathogenesis of hemolytic sickle cell disease (44). Furthermore, Hb ingestion activated gene expression signatures associated with phagocytosis, the PI3K-Akt pathway, growth factor activity, and intracellular signaling, enriched categories with several links to actin remodeling and, in some cases, directly to macropinocytosis. Indeed, several genes induced by Hb, such as the receptors *Axl, Csf1r,* and *Met*, the growth factors *Hgf* and *Pdgfa*, and the actin cytoskeleton-interacting factors *Evl* and *Coro1a* are known to play a role in the regulation and/or execution of macropinocytosis (30, 45–47).

**Figure 7.**
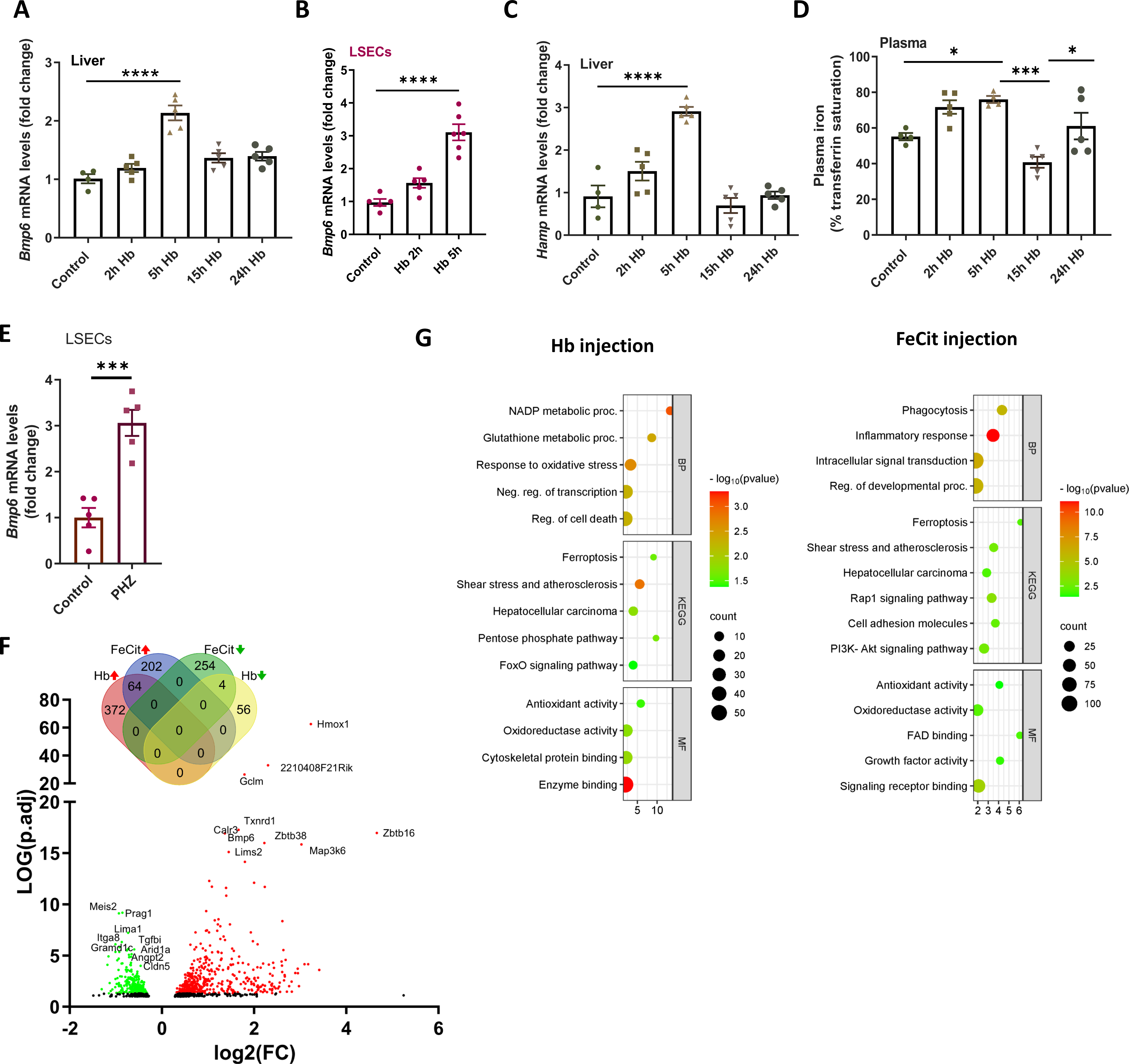
LSECs trigger the iron-sensing BMP6 angiokine upon hemolysis. (A-D) Mice were injected with native hemoglobin (Hb, 10 mg/mouse) and analyzed at indicated time points. (A, B) *Bmp6* and (C) *Hamp* gene expression measured by RT-qPCR in whole livers or FACS-sorted LSECs, as indicated. (D) Plasma iron levels were determined by transferrin saturation measurements. (E) *Bmp6* expression in FACS-sorted LSECs in mice injected with (PHZ, 0.125 mg/g, 6 h). (F-G) Mice were injected with Hb (10 mg) or Ferric citrate (FeCit, 150 µg) for 5 h. (F) A volcano plot of differentially regulated genes identified by the RNA-seq transcriptomic analysis in FACS-sorted LSECs. The green color indicates negatively, and the red color positively-regulated genes. Venn diagram shows genes regulated by Hb and FeCit. (G) functional enrichment among genes induced by Hb or FeCit. The x-axis represents the fold of enrichment. The y-axis represents GO terms of biological processes (BP), Kyoto Encyclopedia of Genes and Genomes terms (KEGG), and GO terms of molecular function (MF). The size of the dot represents the number of genes under a specific term. The color of the dots represents the adjusted P-value. Data are expressed as mean ± SEM and each data point represents one biological replicate. One-way ANOVA with Tukey’s Multiple Comparison tests was used in A-D, while Welch’s unpaired t-test was used to determine statistical significance in E; ns - not significant, *p<0.05, ***p<0.001 and ****p<0.0001.

Collectively, these findings support the role of LSECs in Hb clearance and demonstrate their high capacity to trigger the iron-sensing *Bmp6* angiokine in response to excessive Hb, along with specific pro-inflammatory markers and factors associated with actin remodeling.

## Discussion

Recent advances have revealed the critical role of specialized hepatic sinusoidal endothelium in maintaining systemic and liver homeostasis. The scavenging activity of LSECs is essential for the hepatic clearance of biological macromolecules, such as denatured collagen, glycans/glycation end products, modified low-density lipoproteins, small immune complexes, lipopolysaccharides and viruses (20, 21, 27, 48, 49). In addition, LSECs regulate the vascular tone in the liver, balance the tolerogenic immune milieu with the onset of immune responses, ensure physiological zonation of hepatic immune cells (49, 50), and control iron balance by producing the angiokines BMP2 and BMP6 (22, 23). Our study identified the novel homeostatic role of LSECs in the clearance of free Hb, an activity in which their scavenging and iron-regulatory functions converge.

The clearance functions of LSECs have been attributed to their extraordinary endocytic activity and surface expression of scavenger receptors (20). We discovered that LSECs have a remarkable and constitutive capacity for macropinocytosis, similar to that of macrophages, which use this pathway for antigen presentation, and in contrast to the inducible macropinocytosis of cancer cells, which is critical for nutrient acquisition (29, 31). We showed that this capability distinguishes LSECs from ECs of other organs, thus significantly expanding the scarce knowledge on the macropinocytic activity of ECs (51) and assigning physiological significance to early observations that LSECs can engage macropinocytosis for antigen capture (52). We propose that the macropinocytosis of LSECs likely contributes to their scavenging function. While this possibility merits future studies, macropinocytosis has recently been identified as a major driver of LDL uptake by macrophages, leading to foam cell formation in atherosclerosis (53). Hence, our findings aid in shifting the paradigm from the sole role of scavenger receptor-mediated endocytosis in the clearance of blood-borne macromolecules, emphasizing the importance of macropinocytic uptake. Further studies are required to elucidate the mechanisms by which β-catenin signaling supports macropinocytosis of LSECs and whether its pharmacological blockade impairs liver clearance functions. Finally, our data also demonstrated that the macropinocytic activity of KCs supports their uptake of both free Hb and Hb complexed with Hp, broadening the significance of this route in Hb clearance.

Our study employed *in vivo* approaches to verify the role of macrophages in Hb uptake and revealed that LSECs qualitatively and quantitatively outcompete CD163-expressing KCs and RPMs in this task. These provocative findings may be explained by the fact that the role of CD163 in Hb uptake has been mainly investigated by its ectopic expression in non-macrophage cells and by using polyclonal anti-CD163 blocking antibodies in cultured macrophages (9, 11, 54). Our study does not exclude the contribution of iron-recycling macrophages to Hb uptake under hemolytic conditions, nor does it imply that other tissues/endothelial cells are unaffected by prolonged and pathological exposure to free Hb and heme (55). The extent to which LSEC-mediated Hb uptake protects other organs in hemolytic disorders, such as sickle cell disease or thalassemia, and whether and how LSEC functions are impaired under such conditions, remains to be determined. It is noteworthy that CD163-expressing macrophages play important anti-inflammatory roles, such as in the tumor microenvironment or arthritis, but these tissue-specific roles do not appear to be related to Hb clearance (56–58).

Most importantly, our study defined and quantified the individual contributions of splenic and hepatic cell types to physiological iron recycling. Consistent with previous studies (3, 4), we provide evidence that RPMs are efficient at phagocytosis of intact stressed RBCs, whereas the liver is the site of scavenging steady-state hemolytic products. KCs were efficient in the uptake of RBC ghosts and outperformed RPMs in Hb uptake. LSECs, owing to their anatomical location, exceptional macropinocytic activity, and expression of iron-recycling proteins, have emerged as a novel cell type involved in the maintenance of iron homeostasis, specialized for the removal of spleen-derived Hb. Consistent with this newly proposed role of LSECs, endothelial-specific FPN knock-out mice have been reported to exhibit marked iron deficiency, anemia, and iron loading in liver NPCs (59). Further studies using cell-type specific genetic deletions of proteins involved in iron recycling are crucial to quantify the exact contribution of RPMs, KCs, and LSECs to the circulating plasma iron pool and thus assess their relative involvement in iron recycling from RBCs.

## Authorship Contributions

KMS, GZ, TPR, ZS, and MMia conceived and planned the experiments; ZS, GZ, AJ, PS, RM, MC, IR, KJ, and MMo performed research; GZ, ZS, AJ, PS, RM, MC, MK, RS analyzed data; GZ, ZS, AJ, PS, RM, MC, KJ, KMS, MMik, visualized data; AE provided CD163 KO mice, DS and GD isolated and provided primary human NPCs; MMik and RS curated data; KMS, TPR supervised the study; TPR, AJ, MMia and AE edited the manuscript; KMS, GZ, ZS wrote the manuscript.

## Supporting information

Gating Strategies

## Acknowledgments

This work was funded by the National Science Centre grant 2018/31/B/NZ4/03676 to KMS and the Foundation of Polish Science grant TEAM TECH/2016-1/8 to TPR. We express gratitude to Marta Niklewicz for her substantial technical assistance in this project. We would like to thank Daria Zdżalik-Bielecka for her valuable advice during the initial phases of the project, and to Rafał Mazgaj for providing technical assistance in conducting experiments with confocal microscopy. Additionally, we extend our thanks to Damian Strzemecki for his technical assistance in performing mice injections. We thank Tara Arvedson (Amgen Inc. USA) for the anti-ferroportin antibody, Elizabeta Nemeth for PR73, and Aneta Suwińska for sharing UBI-GFP/BL6 mice. We thank Julian Connor Eckel for his excellent technical assistance in the isolation and provision of primary human liver cells. Many thanks to Agnieszka Popielska and Anna Kosson, and the staff of the Experimental Medicine Centre (Bialystok, Poland) and Mossakowski Medical Research Institute (Warsaw, Poland) for their technical support. This research was performed thanks to the IIMCB IN-MOL-CELL Infrastructure (RRID: SCR_021630) funded by the European Union–NextGenerationEU under the National Recovery and Resilience Plan. IN-MOL-CELL Infrastructure was also funded by the European Union under Horizon Europe (Project 101059801 - RACE) and by the RACE-PRIME project carried out within the IRAP program of the Foundation for Polish Science co-financed by the European Union under the European Funds for Smart Economy 2021-2027 (FENG).

## References

1. Slusarczyk P, Mleczko-Sanecka K. The Multiple Facets of Iron Recycling. Genes (Basel) 2021;12.

2. Muckenthaler MU, Rivella S, Hentze MW, Galy B. A Red Carpet for Iron Metabolism. Cell 2017;168:344–361.

3. Youssef LA, Rebbaa A, Pampou S, Weisberg SP, Stockwell BR, Hod EA, Spitalnik SL. Increased erythrophagocytosis induces ferroptosis in red pulp macrophages in a mouse model of transfusion. Blood 2018;131:2581–2593.

4. Ma S, Dubin AE, Zhang Y, Mousavi SAR, Wang Y, Coombs AM, Loud M, et al. A role of PIEZO1 in iron metabolism in mice and humans. Cell 2021;184:969–982 e913.

5. Klei TRL, Dalimot J, Nota B, Veldthuis M, Mul FPJ, Rademakers T, Hoogenboezem M, et al. Hemolysis in the spleen drives erythrocyte turnover. Blood 2020;136:1579–1589.

6. Kato GJ, Steinberg MH, Gladwin MT. Intravascular hemolysis and the pathophysiology of sickle cell disease. J Clin Invest 2017;127:750–760.

7. Schaer DJ, Vinchi F, Ingoglia G, Tolosano E, Buehler PW. Haptoglobin, hemopexin, and related defense pathways-basic science, clinical perspectives, and drug development. Front Physiol 2014;5:415.

8. Wang Y, Kinzie E, Berger FG, Lim SK, Baumann H. Haptoglobin, an inflammation-inducible plasma protein. Redox Rep 2001;6:379–385.

9. Kristiansen M, Graversen JH, Jacobsen C, Sonne O, Hoffman HJ, Law SK, Moestrup SK. Identification of the haemoglobin scavenger receptor. Nature 2001;409:198–201.

10. Boretti FS, Baek JH, Palmer AF, Schaer DJ, Buehler PW. Modeling hemoglobin and hemoglobin:haptoglobin complex clearance in a non-rodent species-pharmacokinetic and therapeutic implications. Front Physiol 2014;5:385.

11. Schaer DJ, Schaer CA, Buehler PW, Boykins RA, Schoedon G, Alayash AI, Schaffner A. CD163 is the macrophage scavenger receptor for native and chemically modified hemoglobins in the absence of haptoglobin. Blood 2006;107:373–380.

12. Etzerodt A, Kjolby M, Nielsen MJ, Maniecki M, Svendsen P, Moestrup SK. Plasma clearance of hemoglobin and haptoglobin in mice and effect of CD163 gene targeting disruption. Antioxid Redox Signal 2013;18:2254–2263.

13. Theurl I, Hilgendorf I, Nairz M, Tymoszuk P, Haschka D, Asshoff M, He S, et al. On-demand erythrocyte disposal and iron recycling requires transient macrophages in the liver. Nat Med 2016;22:945–951.

14. Vinchi F, De Franceschi L, Ghigo A, Townes T, Cimino J, Silengo L, Hirsch E, et al. Hemopexin therapy improves cardiovascular function by preventing heme-induced endothelial toxicity in mouse models of hemolytic diseases. Circulation 2013;127:1317–1329.

15. Vinchi F, Sparla R, Passos ST, Sharma R, Vance SZ, Zreid HS, Juaidi H, et al. Vasculo-toxic and pro-inflammatory action of unbound haemoglobin, haem and iron in transfusion-dependent patients with haemolytic anaemias. Br J Haematol 2021;193:637–658.

16. Fagoonee S, Gburek J, Hirsch E, Marro S, Moestrup SK, Laurberg JM, Christensen EI, et al. Plasma protein haptoglobin modulates renal iron loading. Am J Pathol 2005;166:973–983.

17. Marro S, Barisani D, Chiabrando D, Fagoonee S, Muckenthaler MU, Stolte J, Meneveri R, et al. Lack of haptoglobin affects iron transport across duodenum by modulating ferroportin expression. Gastroenterology 2007;133:1261–1271.

18. Poisson J, Lemoinne S, Boulanger C, Durand F, Moreau R, Valla D, Rautou PE. Liver sinusoidal endothelial cells: Physiology and role in liver diseases. J Hepatol 2017;66:212–227.

19. Sorensen KK, McCourt P, Berg T, Crossley C, Le Couteur D, Wake K, Smedsrod B. The scavenger endothelial cell: a new player in homeostasis and immunity. Am J Physiol Regul Integr Comp Physiol 2012;303:R1217–1230.

20. Koch PS, Lee KH, Goerdt S, Augustin HG. Angiodiversity and organotypic functions of sinusoidal endothelial cells. Angiogenesis 2021;24:289–310.

21. Schledzewski K, Geraud C, Arnold B, Wang S, Grone HJ, Kempf T, Wollert KC, et al. Deficiency of liver sinusoidal scavenger receptors stabilin-1 and -2 in mice causes glomerulofibrotic nephropathy via impaired hepatic clearance of noxious blood factors. J Clin Invest 2011;121:703–714.

22. Koch PS, Olsavszky V, Ulbrich F, Sticht C, Demory A, Leibing T, Henzler T, et al. Angiocrine Bmp2 signaling in murine liver controls normal iron homeostasis. Blood 2017;129:415–419.

23. Canali S, Zumbrennen-Bullough KB, Core AB, Wang CY, Nairz M, Bouley R, Swirski FK, et al. Endothelial cells produce bone morphogenetic protein 6 required for iron homeostasis in mice. Blood 2017;129:405–414.

24. Mleczko-Sanecka K, Silvestri L. Cell-type-specific insights into iron regulatory processes. Am J Hematol 2021;96:110–127.

25. Slusarczyk P, Mandal PK, Zurawska G, Niklewicz M, Chouhan K, Mahadeva R, Jonczy A, et al. Impaired iron recycling from erythrocytes is an early hallmark of aging. Elife 2023;12.

26. Strauss O, Phillips A, Ruggiero K, Bartlett A, Dunbar PR. Immunofluorescence identifies distinct subsets of endothelial cells in the human liver. Sci Rep 2017;7:44356.

27. Ganesan LP, Kim J, Wu Y, Mohanty S, Phillips GS, Birmingham DJ, Robinson JM, et al. FcgammaRIIb on liver sinusoidal endothelium clears small immune complexes. J Immunol 2012;189:4981–4988.

28. Guilliams M, Bonnardel J, Haest B, Vanderborght B, Wagner C, Remmerie A, Bujko A, et al. Spatial proteogenomics reveals distinct and evolutionarily conserved hepatic macrophage niches. Cell 2022;185:379–396 e338.

29. Canton J. Macropinocytosis: New Insights Into Its Underappreciated Role in Innate Immune Cell Surveillance. Front Immunol 2018;9:2286.

30. Zdzalik-Bielecka D, Poswiata A, Kozik K, Jastrzebski K, Schink KO, Brewinska-Olchowik M, Piwocka K, et al. The GAS6-AXL signaling pathway triggers actin remodeling that drives membrane ruffling, macropinocytosis, and cancer-cell invasion. Proc Natl Acad Sci U S A 2021;118.

31. Commisso C, Davidson SM, Soydaner-Azeloglu RG, Parker SJ, Kamphorst JJ, Hackett S, Grabocka E, et al. Macropinocytosis of protein is an amino acid supply route in Ras-transformed cells. Nature 2013;497:633–637.

32. Klein D, Demory A, Peyre F, Kroll J, Augustin HG, Helfrich W, Kzhyshkowska J, et al. Wnt2 acts as a cell type-specific, autocrine growth factor in rat hepatic sinusoidal endothelial cells cross-stimulating the VEGF pathway. Hepatology 2008;47:1018–1031.

33. Tejeda-Munoz N, Albrecht LV, Bui MH, De Robertis EM. Wnt canonical pathway activates macropinocytosis and lysosomal degradation of extracellular proteins. Proc Natl Acad Sci U S A 2019;116:10402–10411.

34. Redelman-Sidi G, Binyamin A, Gaeta I, Palm W, Thompson CB, Romesser PB, Lowe SW, et al. The Canonical Wnt Pathway Drives Macropinocytosis in Cancer. Cancer Res 2018;78:4658–4670.

35. Kalucka J, de Rooij L, Goveia J, Rohlenova K, Dumas SJ, Meta E, Conchinha NV, et al. Single-Cell Transcriptome Atlas of Murine Endothelial Cells. Cell 2020;180:764–779 e720.

36. Wang F, Ding P, Liang X, Ding X, Brandt CB, Sjostedt E, Zhu J, et al. Endothelial cell heterogeneity and microglia regulons revealed by a pig cell landscape at single-cell level. Nat Commun 2022;13:3620.

37. Dolat L, Spiliotis ET. Septins promote macropinosome maturation and traffic to the lysosome by facilitating membrane fusion. J Cell Biol 2016;214:517–527.

38. Ramirez C, Hauser AD, Vucic EA, Bar-Sagi D. Plasma membrane V-ATPase controls oncogenic RAS-induced macropinocytosis. Nature 2019;576:477–481.

39. Maxson ME, Sarantis H, Volchuk A, Brumell JH, Grinstein S. Rab5 regulates macropinocytosis by recruiting the inositol 5-phosphatases OCRL and Inpp5b that hydrolyse PtdIns(4,5)P2. J Cell Sci 2021;134.

40. Schnatwinkel C, Christoforidis S, Lindsay MR, Uttenweiler-Joseph S, Wilm M, Parton RG, Zerial M. The Rab5 effector Rabankyrin-5 regulates and coordinates different endocytic mechanisms. PLoS Biol 2004;2:E261.

41. Stefanova D, Raychev A, Deville J, Humphries R, Campeau S, Ruchala P, Nemeth E, et al. Hepcidin Protects against Lethal Escherichia coli Sepsis in Mice Inoculated with Isolates from Septic Patients. Infect Immun 2018;86.

42. Makia NL, Bojang P, Falkner KC, Conklin DJ, Prough RA. Murine hepatic aldehyde dehydrogenase 1a1 is a major contributor to oxidation of aldehydes formed by lipid peroxidation. Chem Biol Interact 2011;191:278–287.

43. Zurawska G, Jonczy A, Niklewicz M, Sas Z, Rumienczyk I, Kulecka M, Piwocka K, et al. Iron-triggered signaling via ETS1 and the p38/JNK MAPK pathway regulates Bmp6 expression. Am J Hematol 2024;99:543–554.

44. Gupta A, Fei YD, Kim TY, Xie A, Batai K, Greener I, Tang H, et al. IL-18 mediates sickle cell cardiomyopathy and ventricular arrhythmias. Blood 2021;137:1208–1218.

45. BoseDasgupta S, Moes S, Jenoe P, Pieters J. Cytokine-induced macropinocytosis in macrophages is regulated by 14-3-3zeta through its interaction with serine-phosphorylated coronin 1. FEBS J 2015;282:1167–1181.

46. Visweshwaran SP, Nayab H, Hoffmann L, Gil M, Liu F, Kuhne R, Maritzen T. Ena/VASP proteins at the crossroads of actin nucleation pathways in dendritic cell migration. Front Cell Dev Biol 2022;10:1008898.

47. Recouvreux MV, Commisso C. Macropinocytosis: A Metabolic Adaptation to Nutrient Stress in Cancer. Front Endocrinol (Lausanne) 2017;8:261.

48. Malovic I, Sorensen KK, Elvevold KH, Nedredal GI, Paulsen S, Erofeev AV, Smedsrod BH, et al. The mannose receptor on murine liver sinusoidal endothelial cells is the main denatured collagen clearance receptor. Hepatology 2007;45:1454–1461.

49. Shetty S, Lalor PF, Adams DH. Liver sinusoidal endothelial cells - gatekeepers of hepatic immunity. Nat Rev Gastroenterol Hepatol 2018;15:555–567.

50. Gola A, Dorrington MG, Speranza E, Sala C, Shih RM, Radtke AJ, Wong HS, et al. Commensal-driven immune zonation of the liver promotes host defence. Nature 2021;589:131–136.

51. Lin XP, Mintern JD, Gleeson PA. Macropinocytosis in Different Cell Types: Similarities and Differences. Membranes (Basel) 2020;10.

52. Connolly MK, Bedrosian AS, Malhotra A, Henning JR, Ibrahim J, Vera V, Cieza-Rubio NE, et al. In hepatic fibrosis, liver sinusoidal endothelial cells acquire enhanced immunogenicity. J Immunol 2010;185:2200–2208.

53. Lin HP, Singla B, Ahn W, Ghoshal P, Blahove M, Cherian-Shaw M, Chen A, et al. Receptor-independent fluid-phase macropinocytosis promotes arterial foam cell formation and atherosclerosis. Sci Transl Med 2022;14:eadd2376.

54. Schaer CA, Schoedon G, Imhof A, Kurrer MO, Schaer DJ. Constitutive endocytosis of CD163 mediates hemoglobin-heme uptake and determines the noninflammatory and protective transcriptional response of macrophages to hemoglobin. Circ Res 2006;99:943–950.

55. Buehler PW, Humar R, Schaer DJ. Haptoglobin Therapeutics and Compartmentalization of Cell-Free Hemoglobin Toxicity. Trends Mol Med 2020;26:683–697.

56. Etzerodt A, Tsalkitzi K, Maniecki M, Damsky W, Delfini M, Baudoin E, Moulin M, et al. Specific targeting of CD163(+) TAMs mobilizes inflammatory monocytes and promotes T cell-mediated tumor regression. J Exp Med 2019;216:2394–2411.

57. Etzerodt A, Moulin M, Doktor TK, Delfini M, Mossadegh-Keller N, Bajenoff M, Sieweke MH, et al. Tissue-resident macrophages in omentum promote metastatic spread of ovarian cancer. J Exp Med 2020;217.

58. Svendsen P, Etzerodt A, Deleuran BW, Moestrup SK. Mouse CD163 deficiency strongly enhances experimental collagen-induced arthritis. Sci Rep 2020;10:12447.

59. Zhang Z, Guo X, Herrera C, Tao Y, Wu Q, Wu A, Wang H, et al. Bmp6 expression can be regulated independently of liver iron in mice. PLoS One 2014;9:e84906.

