## Supplementary Figures for "Liver sinusoidal endothelial cells constitute a major route for hemoglobin clearance"

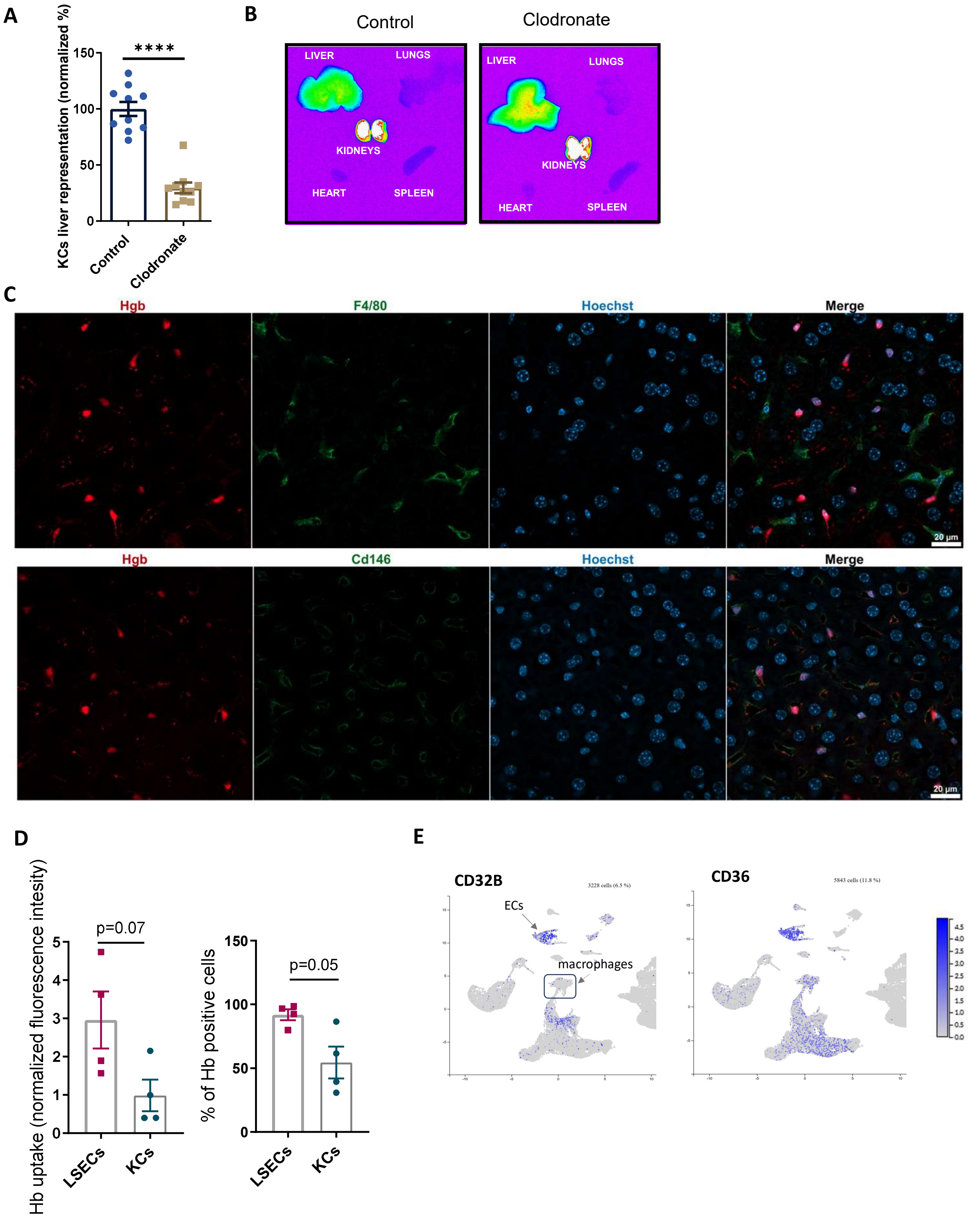


**Figure S1. LSECs represent the major cell type that sequesters Hb.**
(A-B) Hemoglobin (Hb) distribution in control and macrophage-depleted mice (clodronate) injected with Alexa Fluor 750 labeled Hb (Hb-AF750, 10 µg/mouse), imaged with Bruker *in vivo* Imaging System. (A) Efficiency of macrophage depletion in the liver examined by the percentage of liver KCs in control and clodronate injected mice. (B) Representative images of organs isolated from Hb-AF750 (10 µg/mouse, 1 h) *i.v.*-injected mice. (C) Frozen liver slices from mice injected with Hb-AF647 (red) were processed and stained for Kupffer cells (KCs) (F4/80, green) or LSECs (CD146, green) and nuclei (blue). (D) Murine NPCs *in vitro* cultures were treated with Hb-AF750 (0.5 µg/ml) for 1 h. Normalized Hb-AF750 fluorescence intensity and percentage of Hb-AF750$^{+}$ LSECs and KCs, measured with flow cytometry. (E) Plots illustrating high mRNA expression of human LCES markers CD32B and CD36 in liver ECs, visualized using the Human Liver Cell Atlas, Guilliams et al. 2022). Data are expressed as mean ± SEM and each data point represents one biological replicate. Welch's unpaired t-test was used to determine statistical significance in panel A and D, ****p<0.0001.

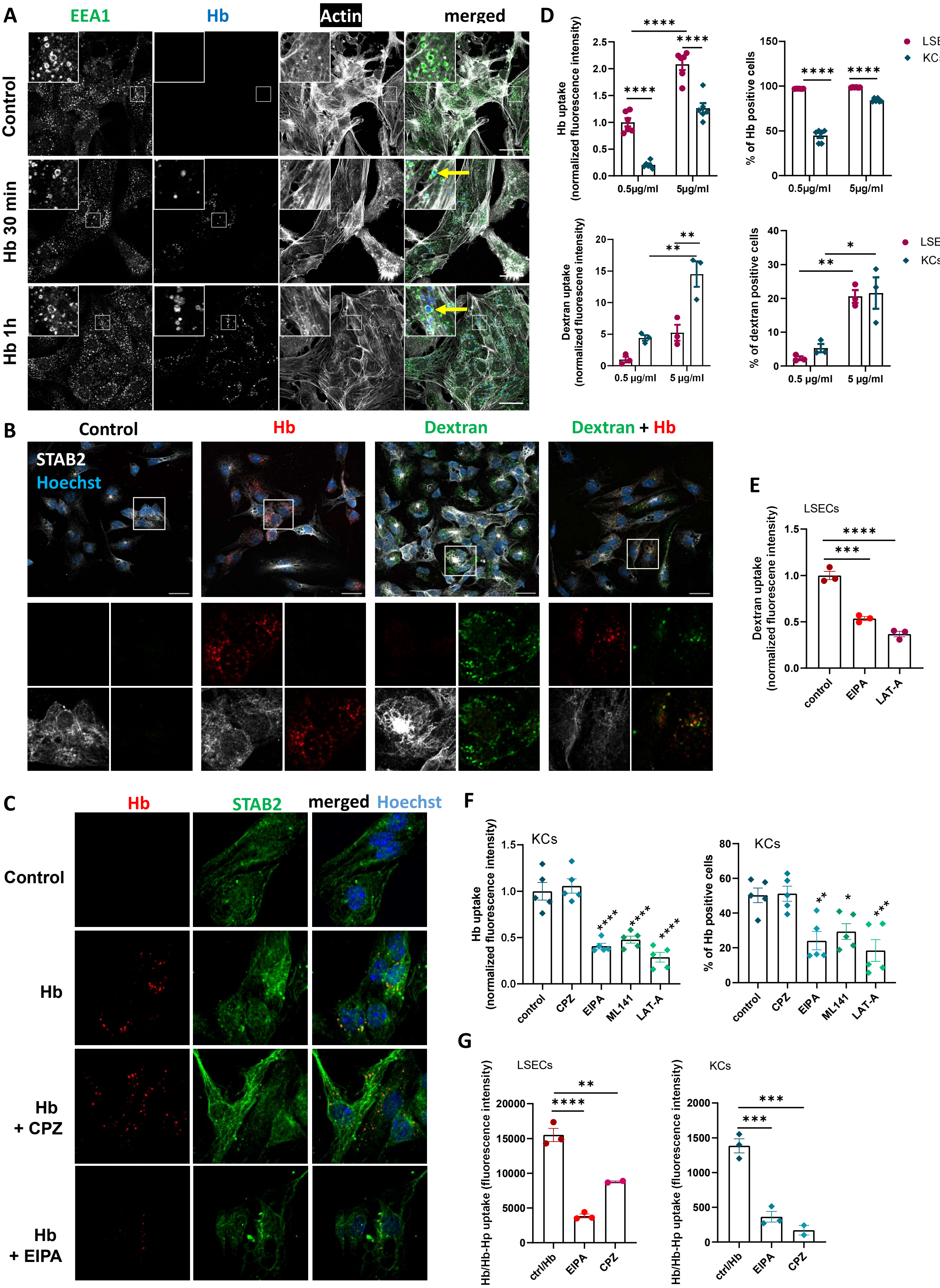


**Figure S2. Primary LSECs and KCs take up Hb via macropinocytosis.**
Murine NPCs *in vitro* cultures were treated with Hb-AF750 or fluorescently-labeled dextran (0.5 or 5 µg/ml) for 10 min, 30 min or 1 h. (A) Hb-AF647 vesicle localization was imaged in NPCs *in vitro* cultures depleted from macrophages. Arrows indicate Hb-AF647 (blue) presence in the EEA-1$^{+}$ (green) and Actin$^{+}$ (white) vesicles. Areas in the highlighted rectangles are shown at higher magnification (left upper corner). Scale bars, 20 µm. (B) Co-localization of Hb-AF647 (red) with rhodamine-dextran (green) was imaged in STAB2$^{+}$ LSECs (white). Arrow indicates co-localization of Hb and rhodamine-dextran at the 10 min time-point. Area in the highlighted rectangle is shown at higher magnification in separate channels below. Scale bar, 20 µm. (C) NPCs were pretreated with the inhibitor of clathrin-mediated endocytosis chlorpromazine (CPZ, 2 µM) or the macropinocytosis blocker EIPA (25 µM), before Hb-AF750 treatment for 10 min. Cells were fixed and stained for STAB2 (green) and nuclei (blue). Scale bars, 20 µm. (D-F) Hb-AF750/FITC-dextran fluorescence intensity and percentage of Hb-AF750$^{+}$/dextran$^{+}$ LSECs or KCs, measured with flow cytometry 1 h after (D) administration of the increasing doses of the indicated cargo or (E and F) upon pretreatment with the indicated inhibitors. (G) Hb:Hp-AF750 fluorescence intensity in LSECs or KCs upon pretreatment with the indicated inhibitors, measured with flow cytometry 1 h after administration of Hb-Hp-AF750. Numerical data are expressed as mean ± SEM and each data point represents one biological replicate. Two-way ANOVA with Tukey's Multiple Comparison test was used to determine statistical significance in D; one-way ANOVA with Tukey's Multiple Comparison test was used to determine statistical significance in E-G; ns - not significant, *$p<0.05$, **$p<0.01$, ***$p<0.001$ and ****$p<0.0001$.

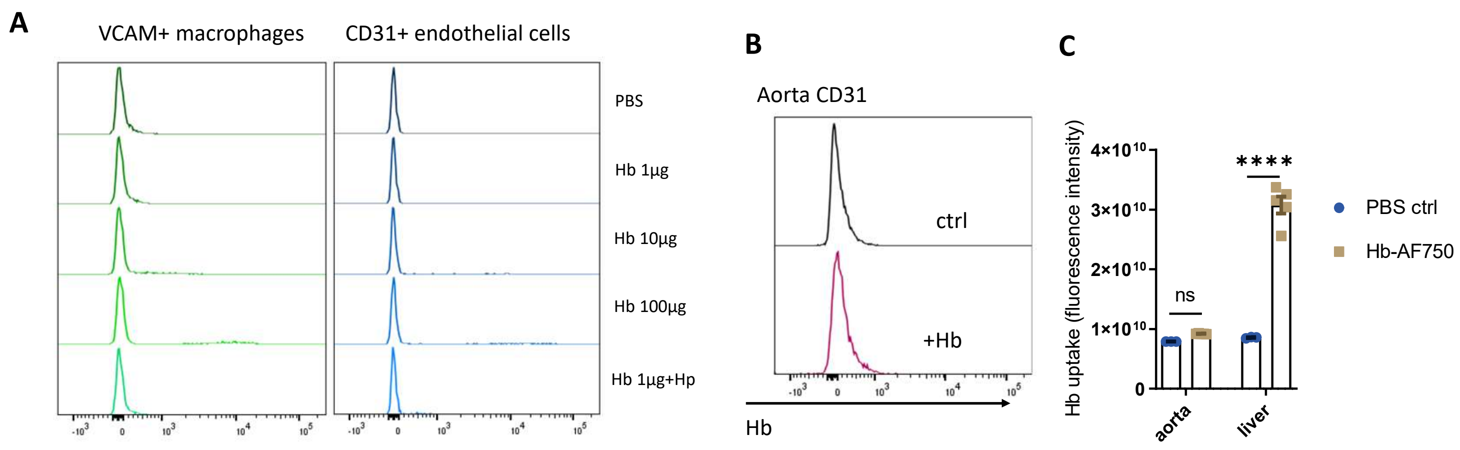


**Figure S3. Bone marrow cells and aortic endothelium fail to sequester Hb.**

Mice were injected i.v. with 1, 10, 100 µg of Hb-AF750 or Hb-AF750:Hp complex (1 µg: 32 µg) for 1 h. (A) Histograms of Hb-AF750 fluorescence in VCAM$^{+}$ macrophages or CD31$^{+}$ endothelial cells in the bone marrow. (B) Histograms of Hb-AF750 uptake by CD31$^{+}$ endothelial cells from aorta, 1 h after Hb-AF750 injection (10 µg/mouse). (C) Mice were injected with Hb-AF750 (10 µg/mouse) for 1 h, and livers and aortas were excised, and total fluorescence was measured by Bruker *in vivo* Imaging System. Quantification of the signal from Hb-AF750 in the organs ARE presented as total fluorescent counts. Data are expressed as mean ± SEM and each data point represents one biological replicate. Two-way ANOVA with Tukey's Multiple Comparison tests was used to determine statistical significance in C; ns - not significant and ****p<0.0001.

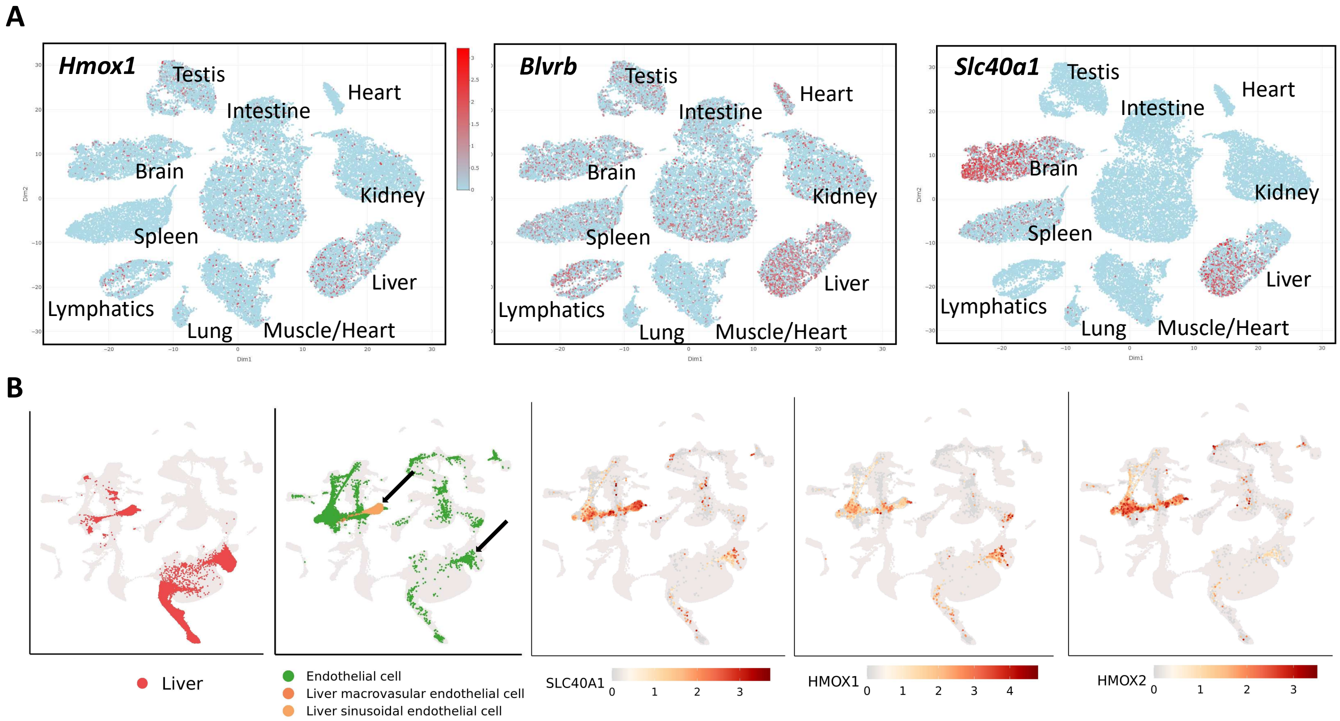


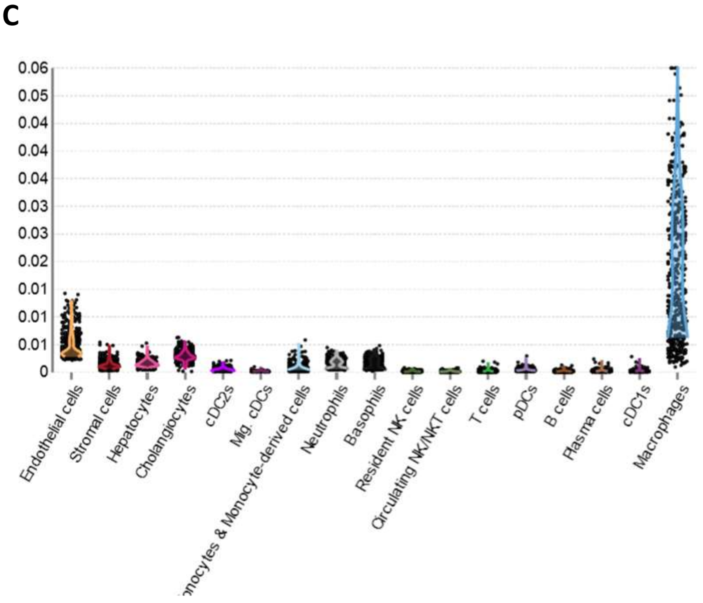


**Figure S4. Liver endothelial cells are distinguished by high mRNA expression levels of genes involved in iron recycling.**

(A) Visualization of the expression levels of *Hmox1*, *Blvrb* and *Slc40a1* obtained with EC Atlas which collects data from single-cell RNA sequencing of endothelial cells (ECs) from different mouse organs (Kalucka et al. 2020). (B) Visualization of the expression levels of *Slc40a1*, *Hmox1* and *Hmox2* in pig liver endothelial cells (determined by single-cell RNA-seq; Wang et al. 2022).(C) Endothelial cells show second highest expression of *Slc40a1* among all liver cells, as determined by single-cell RNA seq (Liver Cell Atlas; Guilliams et al. 2022).

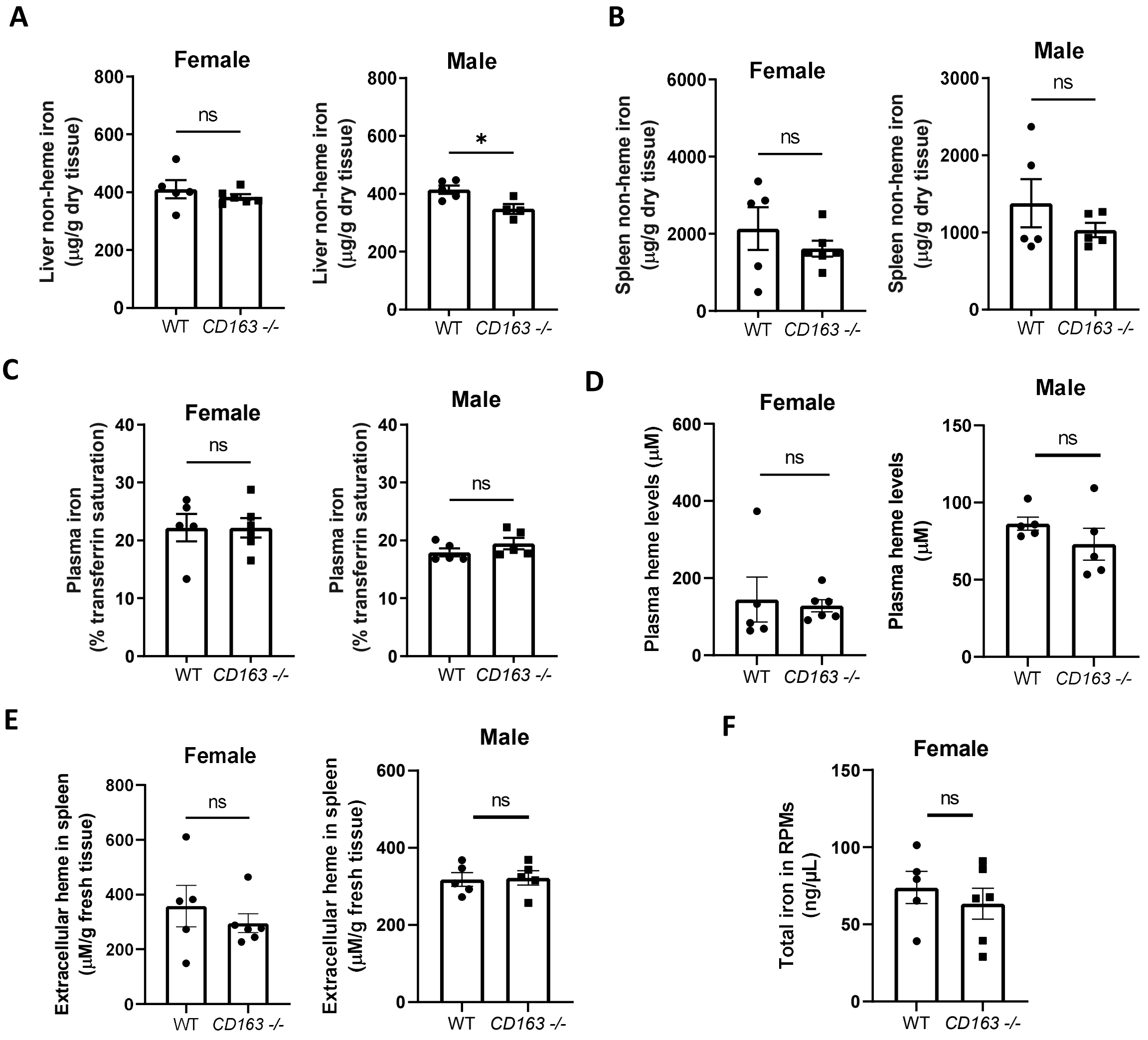


**Figure S5. CD163 KO mice show no major differences in systemic and splenic iron parameters**

(A-E) The phenotype of *Cd163*$^{-/-}$ mice was compared with *wild-type* (WT) littermates. (A-B) Non-heme iron content in (A) the liver and (B) spleen of female and male mice. (C) Plasma iron levels were determined by transferrin saturation measurements. (D,E) Heme levels were measured in the (D) plasma and (E) extracellular fluid from the spleen using Heme Assay Kit. (F) Total iron levels in magnetically-sorted RPMs were measured with Iron assay kit. Data are expressed as mean ± SEM and each data point represents one biological replicate. Welch's unpaired t-test was used to determine statistical significance; ns - not significant, *p<0.05.

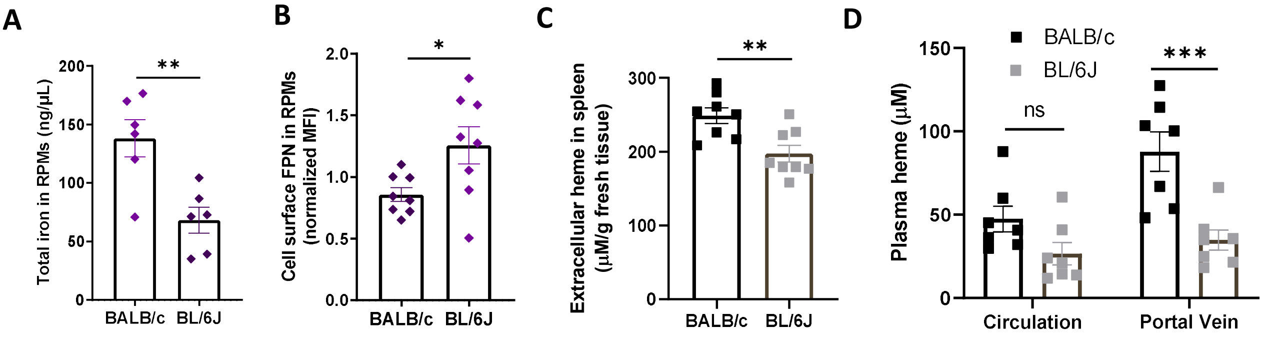


**Figure S6. Comparison of parameters of splenic iron-recycling between BALB/c and C57BL/6J mice.**

(A) The total intracellular iron content in magnetically-sorted RPMs was assessed using the Iron Assay Kit. (B) FPN surface levels were measured in RPMs by flow cytometry. (C) Extracellular heme content in the spleen and (D) heme levels in the portal vein and circulation plasma were measured with Heme Assay Kit. Data are expressed as mean ± SEM and each data point represents one biological replicate. Welch's unpaired t-test was used to determine statistical significance in A-C, while two-way ANOVA with Tukey's Multiple Comparison tests was used in D; ns - not significant, *p<0.05, **p<0.01 and ***p<0.001.

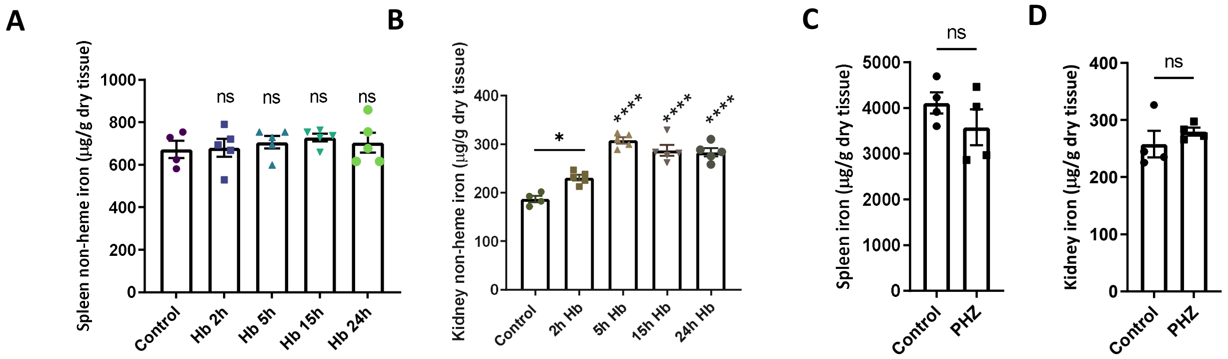


**Figure S7. Alterations of splenic and renal iron levels upon Hb injection and PHZ-induced hemolysis.**

(A-B) Mice were injected with Hb (10 mg/mouse) for indicated time points. Non-heme iron content in the (A) spleens and (B) kidneys. (C-D) Hemolysis was induced by i.p. injection of phenylhydrazine (PHZ, 0.125 mg/g) for 6 h. Non-heme iron content in the (C) spleens and (D) kidneys. Data are expressed as mean ± SEM and each data point represents one biological replicate. Welch's unpaired t-test was used to determine statistical significance in C and D, one-way ANOVA with Tukey's Multiple Comparison test was used in A and B; ns - not significant, *p<0.05 and ****p<0.0001.

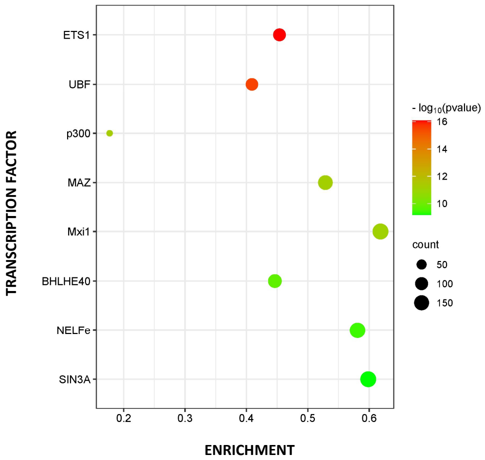


**Figure S8. ETS1 emerged as a key transcription factor responsible for LSEC transcriptome response upon Hb injection.**

The set of genes induced in FACS-sorter LSECs upon injection of mice with 10 mg of Hb was compared with a genome-wide ChIP-seq dataset using the *Cscan* program. Potential common transcriptional regulators of input genes are shown.
