## Supplementary Methods for "Liver sinusoidal endothelial cells constitute a major route for hemoglobin clearance"

### Primary cell culture

Murine liver cells were isolated from female BALB/c mice according to the standard two-step perfusion method with minor modifications. Briefly, mice were euthanized, and livers were perfused in situ (5 ml/min) with Liver Perfusion Medium (Gibco, #17701038) for 15 min through the inferior vena cava after transection of the portal vein. Next, the medium was exchanged for prewarmed (37°C) Liver Digest Medium (Gibco, #17703034), and perfusion continued for the next 15 min. Digested livers were gently disintegrated in Hepatocyte Wash Medium (Gibco, #17704024), and filtered through a 100-µm cell strainer. The cell suspension was then centrifuged at 50 g for 3 min to separate hepatocytes from the supernatant containing non-parenchymal cells (NPCs). Hepatocytes were further purified by layering them on 1.06 g/mL Percoll solution (Sigma-Aldrich, #GE17-0891-01) and centrifugation at 750 g for 20 min without break. NPCs were centrifuged 2 more times at 50 g, for 3 min, and spun at 650 g for 15 min, 4°C. The pellet was resuspended in RBCs Lysis Buffer (BioLegend, #420301), incubated for 3 min, washed, and centrifuged again for 15 min, 650 g, 4°C. The resulting pellet was resuspended in RBC Lysis Buffer (BioLegend, #420301) and incubated for 3 min. The NPCs were then washed and centrifuged again at 650 g for 15 min at 4°C. Subsequently, the NPCs were resuspended in Williams E Medium (Sigma-Aldrich, #W1878-500ML), containing 10% FBS, 1% Penicillin/Streptomycin, 1% Glutamax (Gibco, #35050061), and insulin at a concentration of 10 µg/ml (Sigma-Aldrich, #I9278). Cells were seeded on collagen I-coated (Corning, # 354236) plates or glass microscope slides at a density of 30,000 cells/cm$^2$. After 3 hours, the cells were washed with PBS, and the medium was replaced with Williams E Medium supplemented with 4% FBS, 1% Penicillin/Streptomycin, and 1% Glutamax.

Primary human NPCs (hNPCs) and hepatocytes were isolated from the same liver tissue samples using a two-step collagenase perfusion technique (1, 2). Briefly, liver tissue was perfused first with a buffer containing ethylenediaminetetraacetic acid (EGTA) (Sigma-Aldrich, USA, #03777-10 g,) followed by a digestion buffer with collagenase P (Roche, Switzerland, #11213873001). Hepatocytes were separated by washing and centrifuging the cell suspension twice at 50 g in phosphate-buffered saline (PBS) with $Mg^{2+}/Ca^{2+}$ (Gibco, USA, #14040174). The supernatants containing hNPCs were then united and centrifuged at 300 g and subsequently at 650 g (3). Cell pellets containing hNPCs were resuspended in PBS with $Mg^{2+}/Ca^{2+}$, and the hNPC fraction was counted and checked for viability using the Trypan blue (Sigma Aldrich, #T8154) exclusion method. Finally, the hNPC fraction was prepared for shipment by centrifugation, resuspension in Chill Protec Plus (Merck, Germany, #F2295), and transfer into cryovials (Sarstedt, Nümbrecht, Germany, #72.379).

After overnight transport at 4ºC, the hNPCs pellet was washed with Williams E Medium (Sigma-Aldrich, #W1878-500ML), containing 10% FBS, and centrifuged for 5 min, 650g, RT (room temperature). The pellet was resuspended in RBC Lysis Buffer and incubated for 5 min. The hNPCs were then washed and centrifuged again at 650 g for 10 min at RT. hNPCs pellet was resuspended in Williams E Medium (Sigma-Aldrich, #W1878-500ML), containing 10% FBS, 1% Penicillin/Streptomycin, 1% Glutamax (Gibco, #35050061), and human insulin at a concentration of 5 µg/ml (Capricorn, # INS-K) and seeded on collagen-I- coated plates.

**Cell culture treatments**

Murine NPCs primary cells were cultured in Williams E medium supplemented with 4% FBS, 1% Penicillin/Streptomycin, and 1% Glutamax. The next day after cells seeding cells were stimulated with freshly prepared fluorescently stained hemoglobin (Hb-AF647 or Hb-AF750, 0.5 µg/ml), hemoglobin-haptoglobin complex (Hb-AF750:Hp, 0.5 µg:15 µg/ml) or Dextran Tetramethylrhodamine, 70,000 MW, Lysine Fixable (1 mg/ml, ThermoScientific, #D1818) for 1 h. When indicated, the following compounds were used for pre-treatments: Wnt/beta-catenin inhibitor PRI-724 (5 µM, for 24 h, Selleckchem, #S8968); inhibitor of clathrin-mediated endocytosis, chlorpromazine (2 µM, for 1 h, Sigma-Aldrich, #C8138); inhibitor of macropinocytosis, EIPA (25 µM, for 1 h, MedChemExpress, #HY-101840); inhibitor of Cdc42 GTPase, ML141 (10 µM, for 1 h, Selleckchem, #S7686), an actin-depolymerizing agent, latrunculin A (1 µM, for 30 min, Tocris, #3973) and inhibitor of caveolae-mediated endocytosis, nystatin (240U/ml for 30min, Sigma-Aldrich,# N1638).

The next day after seeding, hNPCs were stimulated with freshly prepared fluorescently stained human hemoglobin (Sigma, #H7379; Hb-AF647, 0.5 µg/ml for immunofluorescence imaging, 5 µg/ml for flow cytometry analysis) or Dextran, Texas Red™, 70,000 MW, Lysine Fixable (1 mg/ml, ThermoScientific, # D1864) for 1 h. When indicated, the following compounds were used for pre-treatments: chlorpromazine (2 µM, for 1 h, Sigma-Aldrich, #C8138); EIPA (25 µM, for 1 h, MedChemExpress, #HY-101840) and latrunculin A (1 µM, for 30 min, Tocris, #3973).

**Isolation of liver non-parenchymal cells for FACS sorting**

For FACS-sorting of murine liver sinusoidal endothelial cells (LSECs) and Kupffer cells (KCs), livers were perfused as described above with an additional step of purification. Briefly, NPCs depleted of RBCs were mixed with 1.06 g/mL Percoll (Sigma-Aldrich, #GE17-0891-01) solution (1:1) and centrifuged at 800 g, 30 min at RT. Cell pellets were resuspended in PBS and stained with LIVE/DEAD™ Fixable Violet Cell Stain Kit (ThermoScientific, #L34955) according to the manufacturer's instructions, and centrifuged at 650 g, 5 min, 4ºC. Next, cells were resuspended with FACS buffer (1% BSA in PBS) and incubated with Truestain Fcblock (BioLegend, #101320) 1:100 for 5 min. Next cells were stained with the following antibodies (BioLegend): rat anti-mouse CD45 Pe-Cy7 (#103114), F4/80 APC-Cy7 (#123118), CD11b PerCP (#1012300), CD146 PE (#134704) and unconjugated rabbit-anti mouse STAB2 (St John's Laboratory, #STJ192359) for 30 min at 4°C, in 1:100 dilution. Cells were washed and stained with the secondary antibodies donkey-anti-rabbit Alexa Fluor 647 (ThermoScientific, #A-31571) or donkey-anti-rabbit Alexa Fluor 488 (ThermoScientific, #A-21206) in dilution 1:200. Next, cells were washed and sorted using BD FACS Aria II sorter (BD Biosciences). Cells were sorted directly into TRIzol™ LS Reagent (ThermoScientific, #10296028) at a quantity of 20,000-50,000 cells, with a maximum flow rate of 2,000 events per second using an 85 or 100 µm nozzle. The sorting purity was approximately 90% from $CD45^+$ cells. Gating strategy details are reported in the Supplementary Data.

**Preparation of single-cell suspension for flow cytometry**

For flow cytometry analysis of the liver cells, mice were euthanized, and liver lobes were dissected. Next, liver lobes were perfused with PBS until blood removal, minced with scissors and enzymatically digested in RPMI 1640 medium (CAPRICORN, #RPMI-STA) containing type IV collagenase (1 mg/ml, Sigma Aldrich, #C5318-5G) and DNAse (20 mg/ml, Sigma Aldrich, #DN25-1G) for 40 min at 37°C with shaking. Next, the cell suspension was passed through a 100 µm cell strainer, washed with PBS, and centrifuged 2 x at 50 g, 3 min, RT to remove hepatocyte pellets. Next, the supernatant was centrifuged at 650 g, 15 min, and 4°C. Cells were suspended in RBCs Lysis Buffer, incubated for 5 min, RT, then washed with PBS and centrifuged at 650 g, 15 min, 4°C. The cell pellet was resuspended in PBS and stained with antibodies as described below. For the splenocytes, the spleen was minced with scissors and subjected to digestion under similar conditions as the liver but with a shorter incubation time of 20 min at 37°C. Next, the organ pieces were passed through a 100 µm cell separation strainer, washed with PBS, and centrifuged at 500 g, 5 min, and 4°C. Cells were suspended in RBCs Lysis Buffer, incubated for 7 min, RT, next washed with PBS and centrifuged at 500 g, 5 min, 4°C. For the analysis of femurs and tibias, the epiphysis of the dissected bones was cut off and bones were placed vertically in a specially cut tip for an automatic pipette placed in an Eppendorf tube, so that the bone did not touch the bottom. The tubes were centrifuged at 1000 g, 1 min, RT. The cell pellet was suspended in RBCs Lysis Buffer and incubated for 5 min at RT, washed with PBS, and centrifuged at 500 g, 5 min, 4°C. The cell pellet was resuspended in the same enzymatic mixture used for liver and spleen digestion and incubated for 15 min at 37°C with shaking. The cells were then washed with PBS and centrifuged at 500 g, 5 min, and 4°C. Aorta was dissected and minced with scissors. Next tissue was enzymatically digested with a mix of enzymes: type I collagenase, type XI collagenase, type I-s hyaluronidase, and DNase I according to the protocol published previously.(4) The organ pieces were then filtered through a 100 µm cell separation strainer, rinsed with PBS, and centrifuged at 500 g, 5 min, and 4°C.

**Flow cytometry**

Murine cell pellets were resuspended in PBS and stained with LIVE/DEAD™ Fixable Violet/Aqua Cell Stain Kit (ThermoScientific, #L34955, # L34957) or Zombie Aqua™ Fixable Viability Kit (BioLegend, #423101) according to the manufacturer's instructions, and centrifuged at 650 g, 5 min, 4°C. Next, cells were resuspended with FACS buffer (1% BSA in PBS) and incubated with rat serum (5%) and Truestain Fcblock (BioLegend, #101320) 1:100 for 5 min. In the next step, liver cells were stained with the following rat anti-mouse antibodies (BioLegend): anti-CD45 Pe-Cy7 (#103114), anti-F4/80 APC-Cy7 (#123118), anti-CD11b PerCP (#1012300), anti-CD146 PE (#134704) and unconjugated rabbit-anti mouse STAB2 (St John's Laboratory, #STJ192359) for 30 min at 4°C, in 1:100 dilution. Secondary antibody staining was performed with donkey-anti-rabbit IgG Alexa Fluor 647 (ThermoScientific, #A-31571) or donkey-anti-rabbit IgG Alexa Fluor 488 (ThermoScientific, #A-21206) in dilution: 1:200. After discrimination of dead cells and doublets, LSEC cells were gated as a population moderately expressing CD45, lacking macrophage-specific proteins F4/80 and CD11b, and expressing STAB2 and CD146 receptors. KCs were gated as a population with high expression of CD45 and F4/80 and moderate expression of CD11b.

For analysis of the splenic cell populations, the following surface rat anti-mouse BioLegend antibodies were used: anti-CD45 PerCP (#103129), anti-F4/80 PE (#123109), anti-CD11b BrilliantViolet 605 (#101237), anti-CD45R/B220 Pacific Blue (#103227), anti-CD3 Pacific Blue (#100213), anti-Gr1 Pacific Blue (#108430), anti-Ter119 Pacific Blue (#116231) and anti-CD31 PE-Cy7 (#102524). Splenic RPMs were gated as a population negative for CD3, Gr1, B220 and Ter119, with high expression of CD45, F4/80, and moderate expression of CD11b. Splenic endothelial cells were gated as a population negative for CD45, expressing CD31. For analysis bone marrow macrophages and endothelial cells (ECs) the following surface rat anti-mouse BioLegend antibodies were used: anti-CD45 PerCP(#103129), anti-CD11b BrilliantViolet 605 (#101237), anti-VCAM PE (#105713), anti-CD45R/B220 Pacific Blue (#103227), anti-CD3 Pacific Blue (#100213), anti-Gr1 Pacific Blue (#108430), anti-Ter119 Pacific Blue (#116231) and anti-CD31 PE-Cy7 (#102524). Macrophages were gated as a population negative for CD3, Gr1, B220, and Ter119, with high expression of CD45, VCAM, and moderate expression of CD11b. ECs were gated as a population negative for CD45, expressing CD31. Immediately before the analysis, the cell suspension was transferred to tubes with a sieve with a pore diameter of 35 μm.

The content of intracellular ferrous iron (LIP, $Fe^{2+}$) was measured using FerroOrange (DojinD, #F374). Briefly, surface-stained cells were incubated with 1 μM FerroOrange in HBSS for 30 min at 37 °C and analyzed directly by flow cytometry without further washing. Detection of FPN was performed with a non-commercial antibody that recognizes the extracellular loop of mouse FPN [rat monoclonal antibody, Amgen, clone 1C7; directly conjugated using Alexa Fluor 488 Labeling Kit (Abcam, ab236553)](5). For intracellular HO1 staining, surface-stained cells were fixed with 4% PFA and permeabilized with 0.5% Triton-X in PBS. Next, cells were stained for 30 min at 4 °C with primary anti-HO-1 (ENZO, #SPA-896) conjugated with Alexa Fluor 488. Conjugation was performed with Conjugation Kit (Abcam, #ab236553) according to the manufacturer's protocol. Erythrophagocytosis capacity in RPMs was determined by intracellular staining of the erythrocytic marker Ter-119 (BioLegend, #116201 and #116215) as described previously.(5). The panel of antibodies used for FPN and HO-1 staining consisted of anti-CD45 PE-Cy7, anti-F4/80 anti-APC-Cy7, anti-CD11b PerCP, anti-CD146 PE, and anti-STAB2-Alexa Fluor 647 (Gbiosciences, #ITN 2255-647). Due to the wide emission spectrum, the panel employed for experiments involving FerroOrange was modified by reducing the inclusion of anti-CD11b and anti-CD146 markers and included anti-CD45 Pacific Blue (#103126), anti-F4/80 APC-Cy7 (#123118) and anti-STAB2-Alexa Fluor 647 (Gbiosciences, #ITN 2255-647) for liver cells staining and anti-CD45 APC-Cy7 (#103116), anti-CD31 APC (#102509), anti-CD45R/B220 Pacific Blue (#103227), anti-CD3 Pacific Blue (#100213), anti-Gr1 Pacific Blue (#108430), anti-Ter119 Pacific Blue (#116231) for EC isolated from spleen and bone marrow.

After hemoglobin treatment, human hNPCs were washed with FACS buffer and centrifuged at 650g, 5min, 4°C. hNPCs pellets were stained with LIVE/DEAD™ Fixable Aqua Cell Stain Kit (ThermoScientific, # L34957) according to the manufacturer's instructions, and centrifuged at 650 g, 5 min, 4ºC. Next, cells were resuspended with FACS buffer and incubated with human

Human TruStain FcX™ (BioLegend, # 422302) 1:100 for 5 min. In the next step, human liver cells were stained with mouse anti-human CD36 BUV421 (1:100) and CD32B PE (1:100) antibodies for 1h at 4°C. After staining, cells were washed with FACS buffer and centrifuged at 650 g, 5 min, 4°C. Pellets were fixed with 4% PFA for 15min at RT, washed with FACS buffer, and centrifuged at 750 g, 5 min, 4°C.

Analyses were performed using BD LSRFortessa X-20 (BD Biosciences), BD Canto II (BD Biosciences), BD Aria II sorter (BD Biosciences) or CytoFLEX (Beckman Coulter) flow cytometers and were analyzed with FlowJo or CytExpert, respectively. The geometric mean fluorescence intensities (MFI) corresponding to the probes/target protein levels were determined. For quantifications, the MFI of the adequate fluorescence minus one (FMO) control was subtracted from samples MFI, and data were further normalized. All gating strategy details are reported in the Supplementary Data.

**Isolation of murine hemoglobin (Hb) and conjugation of mouse Hb, human Hb and bovine serum albumin (BSA)**

To isolate hemoglobin (Hb) from mouse erythrocytes, peripheral blood was collected after euthanasia by cardiac puncture to heparinized tubes. Blood was centrifuged at 500 g for 5 min, RT, and the plasma fraction, and the buffy coat were discarded. Erythrocytes were suspended in PBS and centrifuged at 500 g, 5 min, and 4°C, and this washing was repeated 4 times. Erythrocytes were then centrifuged at 3000 g, 5 min, 4°C, the pellet was resuspended in 20 ml of cold sterile deionized water and incubated overnight at 4°C. Finally, the solution was centrifuged at 3000 g, 5 min, 4°C, and concentrated using an Amicon® Ultra-15 filter unit with a 10 kDa cutoff. Hb concentration was calculated from the Beer-Lambert law and heme extinction coefficient (167,000 M−1·cm−1) at 415 nm that was measured using NanoDrop 2000 spectrophotometer (Thermo Fisher Scientific).

Alexa Fluor NHS esters Alexa Fluor 647 (#A20006) or Alexa Fluor 750 (#A20111) (Thermo Fisher Scientific) were dissolved in DMSO and diluted in 0.1 M NaHCO3 pH 8.3. Isolated murine hemoglobin, human hemoglobin (Sigma-Aldrich, # H7379) or BSA were dissolved in 0.1 M NaHCO3 pH 8.3. Protein and ester solutions were mixed in a volume ratio of 1:1 at a molar ratio of 1: 2.5, and incubated at 25°C in the dark for 1 h with shaking. Then the conjugate was washed with 0.1M NaHCO3 pH 8.3 and concentrated using an Amicon® Ultra-15 filter unit with 10 kDa cut-off at 4000 g, 15 min, 4°C. Hb concentration was calculated from the Beer-Lambert law and heme extinction coefficient (167,000 $M^{-1}$·$cm^{-1}$) at 415 nm that was measured using NanoDrop 2000 spectrophotometer NanoDropTM (Thermo Fisher Scientific). BSA concentration was measured using standard methods. The effectiveness of conjugation was confirmed by mass spectrometry analysis. To prepare of Hb-haptoglobin (Hp) complex Hb-AF750 and human Hp (Sigma-Aldrich, #SRP6506) solutions were mixed in a molar ratio (Hb-AF750:Hp) of 1:6 and incubated for 30 min at 4°C in the dark. Then the complex was purified from free Hb or Hp by centrifugation using an Amicon® Ultra-0.5 filter unit with 100 kDa cutoff at 4000 g, 3 min, 4°C.

**RBC preparation and staining**

RBC isolation, labeling, and RBC ghost preparations were performed as described previously (5) with some modifications. Briefly, whole blood obtained from the wild-type C57BL/6J and UBI-GFP/BL6 mice were collected to CPDA-1 solution (Sigma-Aldrich, #C4431). The suspensions were mixed with HBSS, loaded on Lymphosep (Biowest, #L0560-500), and centrifuged at 400 g for 15 min at RT. The buffy coat was removed, and the suspension of RBCs was washed with HBSS twice. When indicated RBCs were stressed (sRBCs) by heating for 30 min at 48°C with shaking. $1\times10^{10}$ RBC were resuspended in 1 ml diluent C, mixed with 1 ml diluent C containing 7.5 µM PKH26 (Sigma-Aldrich, #MINI26-1KT), and incubated in the dark for 5 min in 37°C, the reaction was stopped by adding 2 ml HBSS/1% BSA and suspension was washed with HBSS. For *in vivo* experiments, RBCs were resuspended to 50% hematocrit in HBSS. RBC ghosts were obtained by lysis of PKH26-stained RBCs derived from C57BL/6J mice (volume of the suspension was 2-times more than PKH26-stained UBI-GFP/BL6 RBCs) with PBS to water (1:15) mixture followed by at least 3 centrifugations at 13,000 g, 5 min each until the pellet became brighter. The final pellet was resuspended in HBSS.

**Immunofluorescence**

The mouse liver was fixed in 4% PFA at 4 °C for 24 h. Tissues were washed with PBS (3x30min) and soaked in 12.5% and 25% sucrose for 1.5 h and 48 h, respectively. Liver samples were embedded in Cryomatrix medium (Epredia™, #6769006), frozen on liquid nitrogen, sectioned in 10-µm slices using a cryo-microtome (Leica), and stored at -20°C. Before staining, the sections were left in RT for 30 min, surrounded with Pap Pen Liquid Blocker (Ted Pella, 22311) washed in PBS for 10 min, and permeabilized with PBS/ 0.1% Triton X-100 (Thermo Fisher, #85111) for 20 min. Non-specific antibody binding was blocked by incubating sections in PBS/ 3% BSA (Bioshop, #ALB001.250) for 2 h in RT. For LSEC detection, tissues were incubated with anti-CD146 PE (BioLegend, #134704) 1:100 in PBS/3% BSA for 2 h in RT. For KCs detection, tissues were incubated with anti-F4/80 PE (BioLegend, #123110) 1:100 in

PBS/3% BSA for 2 h in RT. After antibody incubation, tissues were washed 5x5 min with PBS/0.1% Triton X-100 to remove unbound antibodies. For nuclear staining, sections were incubated with Hoechst 33342 (ThermoFisher, #H3570) diluted in PBS/0.1% Triton X-100 (final concentration – 1 ug/ml) for 10 min in RT. Next, sections were washed 3x5 min with PBS/0.1% Triton X-100, incubated in PBS for 10 min, and mounted with ProLong™ Glass Antifade Mountant (Thermo Fisher, #P36982). For immunofluorescence staining of *in vitro* cultured NPCs, cells were seeded on round coverslips coated with collagen I (Corning, #354236). For macropinosomes visualization, seeding of NPCs was performed with an additional step of macrophage depletion with anti-F4/80 magnetic beads (Miltenyibiotec, #130-110-443) according to the manufacturer's protocol. For Hb and dextran intracellular colocalization cells were fixed with 4% PFA for 10 min at RT, washed with PBS, and incubated in PBS/5% BSA for 1 h in RT for blocking. After blocking, cells were incubated overnight at 4°C with anti-F4/80 PE and/or anti-STAB2 (St John's Laboratory, #STJ192359, 1:100). After incubation with primary antibody cells were washed with PBS, and incubated with secondary antibody: donkey-anti rabbit IgG AlexaFluor 488 (Thermo Fisher Scientific, #A-31571, 1:300) for 1 h at RT. For FPN staining NPCs were washed with PBS and blocked with 1%BSA/PBS for 1h at RT. After blocking, cells were stained with anti-CD31 FITC (Biolegend, #102506, 1:50), anti- F4/80 PE (Biolegend, #123110, 1:100) and anti-FPN (Kind gift from Tara Arvedson, Amgen, 1:100 -conjugated with AF647 fluorescent dye – Abcam, #ab269823) for 45min at RT. NPCs were washed with PBS and fixed with 4% PFA for 10 min at RT. After fixation cells were washed with PBS, stained with Hoechst 33342 for 10 min at RT, and mounted on the slides with ProLong™ Glass Antifade Mountant.

For intracellular HO1 and biliverdin reductase b (BLVRB) staining, NPCs were washed with PBS and blocked with 1%BSA/PBS for 1h at RT. After blocking, cells were stained with anti-CD31 FITC (Biolegend, #102506, 1:50) and anti- F4/80 PE (Biolegend, #123110, 1:100), washed with PBS and fixed with 4% PFA for 10 min at RT. After fixation, NPCs were washed with PBS, permeabilized with PBS/ 0.1% Triton X-100 for 5 min at RT and blocked again with 1%BSA/PBS for 1h at RT. After blocking, cells were incubated overnight at 4°C with anti-HO1 (Enzo, #ADI-OSA-150-F, 1:100) or anti-BLVRB (Proteintech, #17727-1-AP, 1:100) primary antibodies. After incubation with primary antibody cells were washed with PBS, and incubated with donkey-anti rabbit IgG AlexaFluor 647 (Thermo Fisher Scientific, #A-31571, 1:300) for 1 h at RT. After incubation with secondary antibodies, NPCs were washed with PBS, stained with Hoechst 33342 for 10 min at RT, and mounted on the slides with ProLong™ Glass

Antifade Mountant. Samples were imaged using a confocal microscope LSM 800 (Zeiss) equipped with an EC Plan-Neofluar 40x/1.30 Oil DIC M27 oil objective and T-PMT detectors. Images were acquired with a resolution of 2101 x 2101 pixels. The images were processed using ImageJ software with linear gamma adjustments of contrast and brightness. For macropinosomes visualization, fixed cells were permeabilized and blocked with solution I - PBS/ 0,1% w/v Saponin, 0,2% w/v/gelatin, 5 mg/ml BSA for 10 min RT. After blocking cells were incubated in solution II – PBS 0,01% Saponin $^{+}$0,2% gelatin with primary antibody anti-EEA1 (Enzo Life Sciences, #ALX-210-239, 1:1000) for 1 h at RT. After washing in solution II, cells were incubated with secondary antibody donkey anti-rabbit IgG 488 (Thermo Fisher Scientific, #A-21206, 1:500) for 1 h at RT. Phalloidin Atto 390 (Sigma-Aldrich, #50556, 1:500) to stain actin was added during incubation with fluorescent secondary antibodies. After incubation with antibodies, cells were washed with PBS and mounted on the slides with Moviol. Sections and slides were visualized with a Zeiss LSM 710 equipped with an Objective Plan-Apochromat 63x/1.4 Oil DIC M27 oil objective and T-PMT detectors. Macropinosome size was determined with ZEN 2011 software. The remaining images were processed using Adobe Photoshop software with linear adjustments of contrast and brightness.

Human non-parenchymal cells seeded on collagen I – coated plates (Ibidi, #80841) were washed with PBS and fix with 4% PFA for 15min in RT. After fixation, cells were washed with PBS and block in 2% BSA for 1h. After blocking, cells were incubated with anti-human CD36 PE (Biolegend, #336206, 1:50), anti-human CD32B PE (Biolegend, #398404, 1:50) or anti-human CD36 BUV421 (Biolegend, # 336230, 1:50) antibodies for 1h. For intracellular EEA1 staining, cells were additionally permeabilized and blocked with PBS/ 0,1% w/v Saponin, 0,2% w/v/gelatin, 5 mg/ml BSA solution for 10 min RT. After blocking cells were incubated in PBS 0,01% Saponin $^{+}$0,2% gelatin solution with primary antibody anti-EEA1 (Enzo Life Sciences, #ALX-210-239, 1:1000) for 1 h at RT. After wash, cells were incubated with secondary antibody donkey anti-rabbit IgG 488 (Thermo Fisher Scientific, #A-21206, 1:500) for 1 h at RT.

**Heme analysis and serum iron quantification**

Tissue non-heme iron content was measured using the bathophenanthroline colorimetric method and calculated against dry tissue weight. Serum iron content (SFBC) and unsaturated iron-binding capacity (UIBC) were measured with the SFBC and the UIBC kit (Biolabo, #97408). Transferrin saturation was calculated using the formula SFBC / (SFBC + UIBC) x 100. Heme content was measured as described previously.(5) Briefly, to determine the extracellular heme content in the spleen, the entire spleen was dissected, weighed, and gently mashed through a 100 µm strainer rinsed with 3 ml of HBSS (ThermoScientific, #14025092). The resulting suspension was centrifuged at 400 g for 10 min at 4 °C and supernatant was collected. To eliminate the remaining cells and membranes, the supernatant was centrifuged again at 1000 g for 10 minutes at 4 °C, and the supernatant was transferred to the fresh tube. To determine heme content in the blood, blood from the circulation and portal vein was collected to the heparin-coated tube, centrifuged at 400 g for 10 min at 4 ° C, and plasma was transferred to the fresh tube. Heme concentrations were determined using the Heme Assay Kit (Sigma Aldrich, #MAK316), following the instructions provided by the manufacturer. The absorbance was measured at a wavelength of 400 nm. The amount of heme was calculated against Heme Calibrator and additionally normalized to the initial weight of fresh spleens (for splenic extracellular heme content).

**RNA isolation and qPCR**

RNA from tissues was extracted using TRIzol™ Reagent (ThermoScientific, #15596026) and from sorted cells using TRIzol™ LS Reagent (ThermoScientific, #10296028) according to the manufacturer's instructions with an additional step of RNA precipitation with glycogen (ThermoScientific, #R0551). RNA (100-400 ng) was reverse transcribed into cDNA using random primers (ThermoScientific, #48190011) and RevertAid H Minus Reverse Transcriptase (ThermoScientific, #EP0452). The cDNA products were amplified by quantitative polymerase chain reaction (qPCR) using the SG qPCR Master Mix (2x) (Eurex, #E0401-01). Real-time qPCR (RT-qPCR) was run in LightCycler 96 Real-Time PCR System (Roche). Primers are listed in Table1 below:

| **Gene name** | **Primer Fw (5'->3')** | **Primer Rev (5'->3')** |
|---|---|---|
| *Rpl19* | AGGCATATGGGCATAGGGAAGAG | TTGACCTTCAGGTACAGGCTGTG |
| *Bmp6* | ATGGCAGGACTGGATCATTGC | CCATCACAGTAGTTGGCAGCG |
| *Hmox-1* | AGGCTAAGACCGCCTTCCT | TGTGTTCCTCTGTCAGCATCA |
| *Hamp* | ATACCAATGCAGAAGAGAAGG | AACAGATACCACACTGGGAA |

Table 1. Primers used for RT-qPCR analysis.

**RNA sequencing**

Transcriptome analysis of LSECs was conducted using the AmpliSeq method. RNA integrity was assessed using Agilent RNA 6000 Nano Kit on Agilent™ 2100 Bioanalyzer™, and cDNA library preparation was performed using a commercially available kit (Ion AmpliSeq™ Transcriptome Mouse Gene Expression Kit, Thermo Scientific) following the manufacturer's recommendations. Targeted cDNA fragments were amplified using the Ion AmpliSeq™ Library Kit 2.0. cDNA libraries were measured using Agilent™ 2100 Bioanalyzer™. Raw reads were processed using the Torrent Suite tool and mapped to the mouse genome mm10 AmpliSeqTranscriptome version using Torrent Mapping Alignment Program (TMAP). Data for iron citrate injection were deposited previously in the GEO repository under accession no GSE235976 (secured with a token: ubcbegoyrxglfeb to allow review), whereas for Hb injection under no GSE240270 (with a token: uhkrgywyhledjmj).

**Statistical analysis**

Data are represented as mean ± SEM. The number of mice/samples per group or the number of independent cell-based experiments are shown in the figures. Statistical analysis was conducted using GraphPad Prism software (Version 9.4.1). When two groups were compared two-tailed unpaired Welch's t-test was applied, whereas for more groups, the One-Way or Two-Way Analysis of Variance (ANOVA) with post-hoc Tukey's test for multiple comparisons was performed, as indicated in figure legends. For all experiments, $\alpha=0.05$. Statistically significant changes between means were indicated as follows: ns, not significant for $p>0.05$, * for $p<0.05$, ** for $p<0.01$, *** for $p<0.001$, and **** for $p<0.0001$.

**References**

1. Kegel V, Deharde D, Pfeiffer E, Zeilinger K, Seehofer D, Damm G. Protocol for Isolation of Primary Human Hepatocytes and Corresponding Major Populations of Non-parenchymal Liver Cells. J Vis Exp 2016:e53069.

2. Damm G, Schicht G, Zimmermann A, Rennert C, Fischer N, Kiessig M, Wagner T, et al. Effect of glucose and insulin supplementation on the isolation of primary human hepatocytes. EXCLI J 2019;18:1071-1091.

3. Zimmermann A, Hansel R, Gemunden K, Kegel-Hubner V, Babel J, Blaker H, Matz-Soja M, et al. In Vivo and In Vitro Characterization of Primary Human Liver Macrophages and Their Inflammatory State. Biomedicines 2021;9.

4. Gjurich BN, Taghavie-Moghadam PL, Galkina EV. Flow Cytometric Analysis of Immune Cells Within Murine Aorta. Methods Mol Biol 2015;1339:161-175.

5. Slusarczyk P, Mandal PK, Zurawska G, Niklewicz M, Chouhan K, Mahadeva R, Jonczy A, et al. Impaired iron recycling from erythrocytes is an early hallmark of aging. Elife 2023;12.
