## Supplementary material for "Liver sinusoidal endothelial cells constitute a major route for hemoglobin clearance": Gating Strategies

Gating strategy for Figure S1A

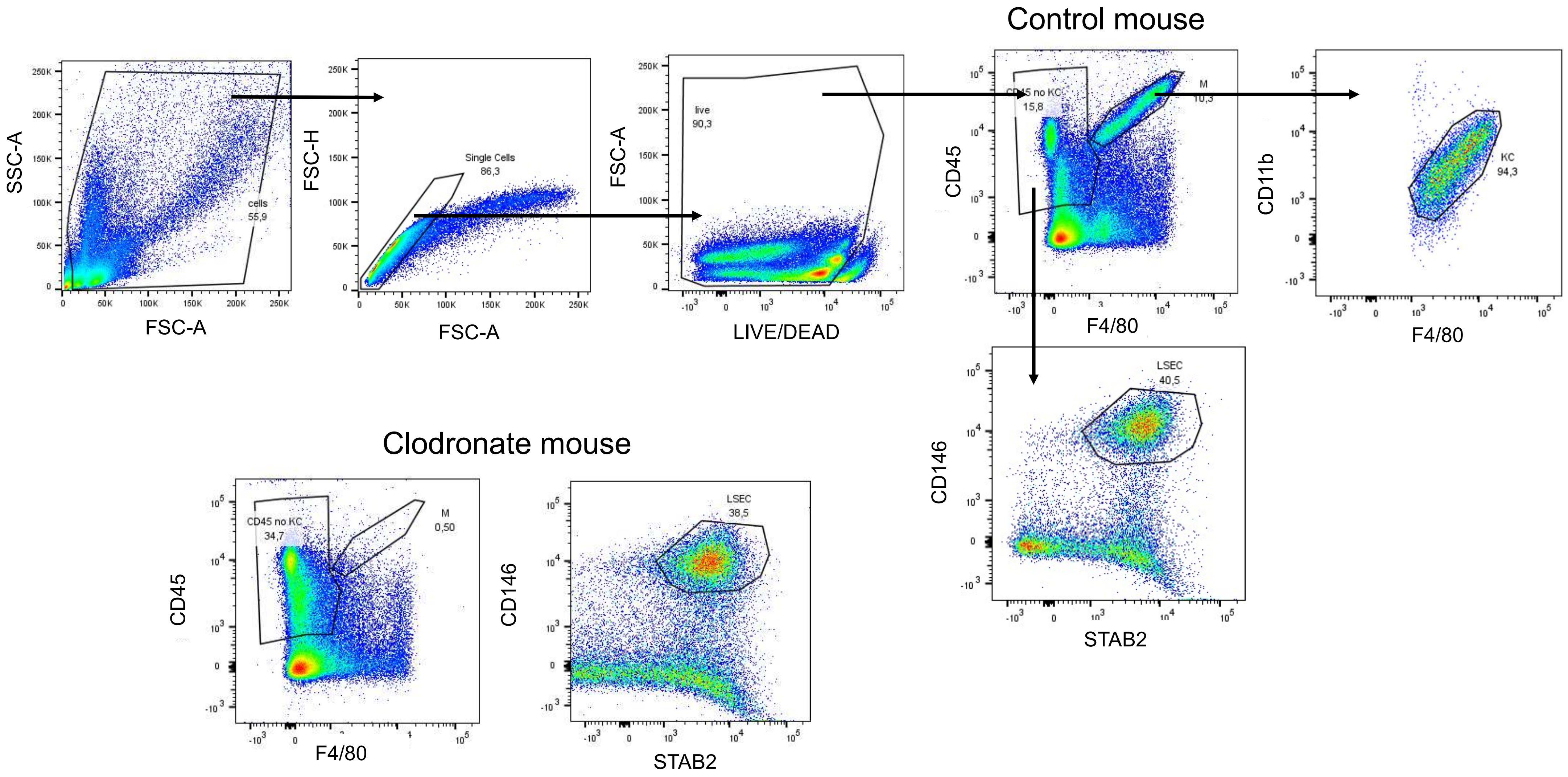

### Gating strategy for Figure 1B

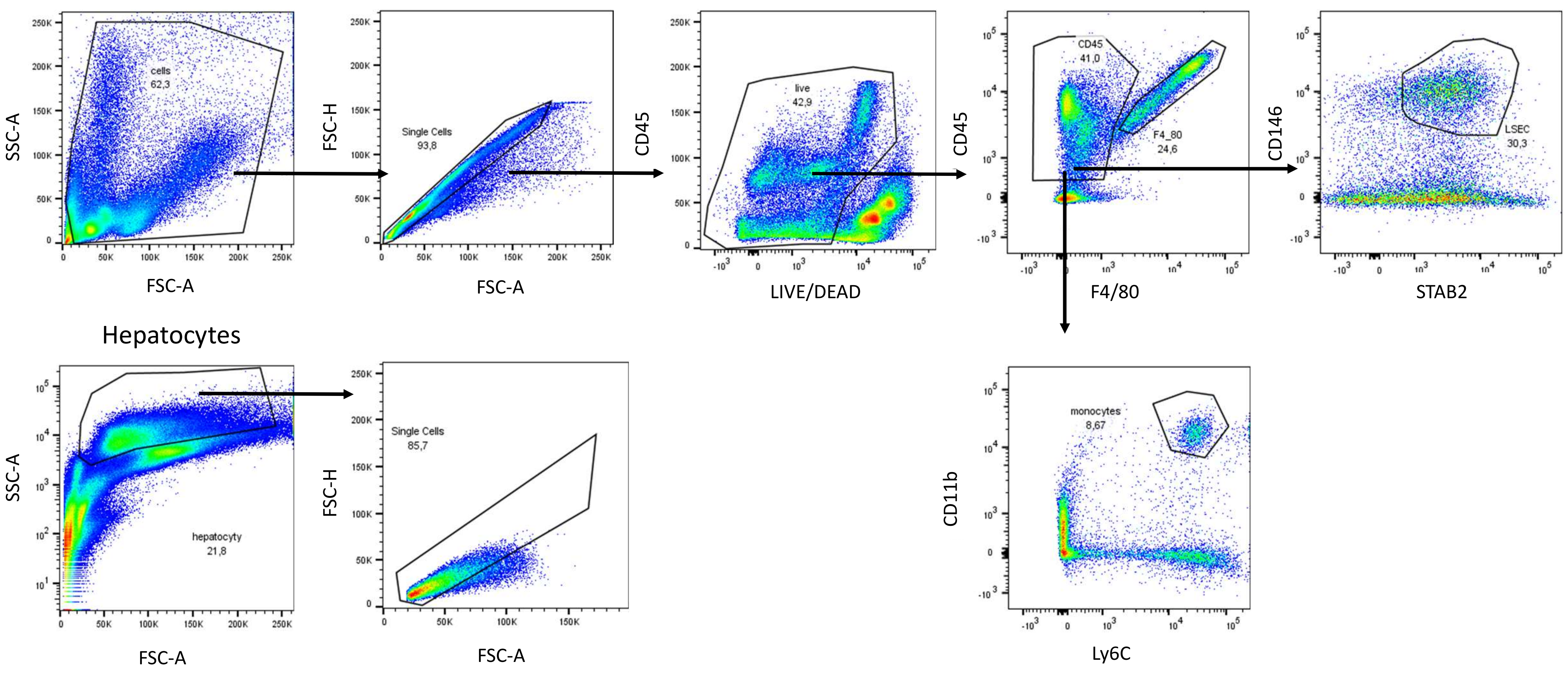

Gating strategy for Figures 2A, B, G, S1D and S2D-G (primary murine liver NPCs)

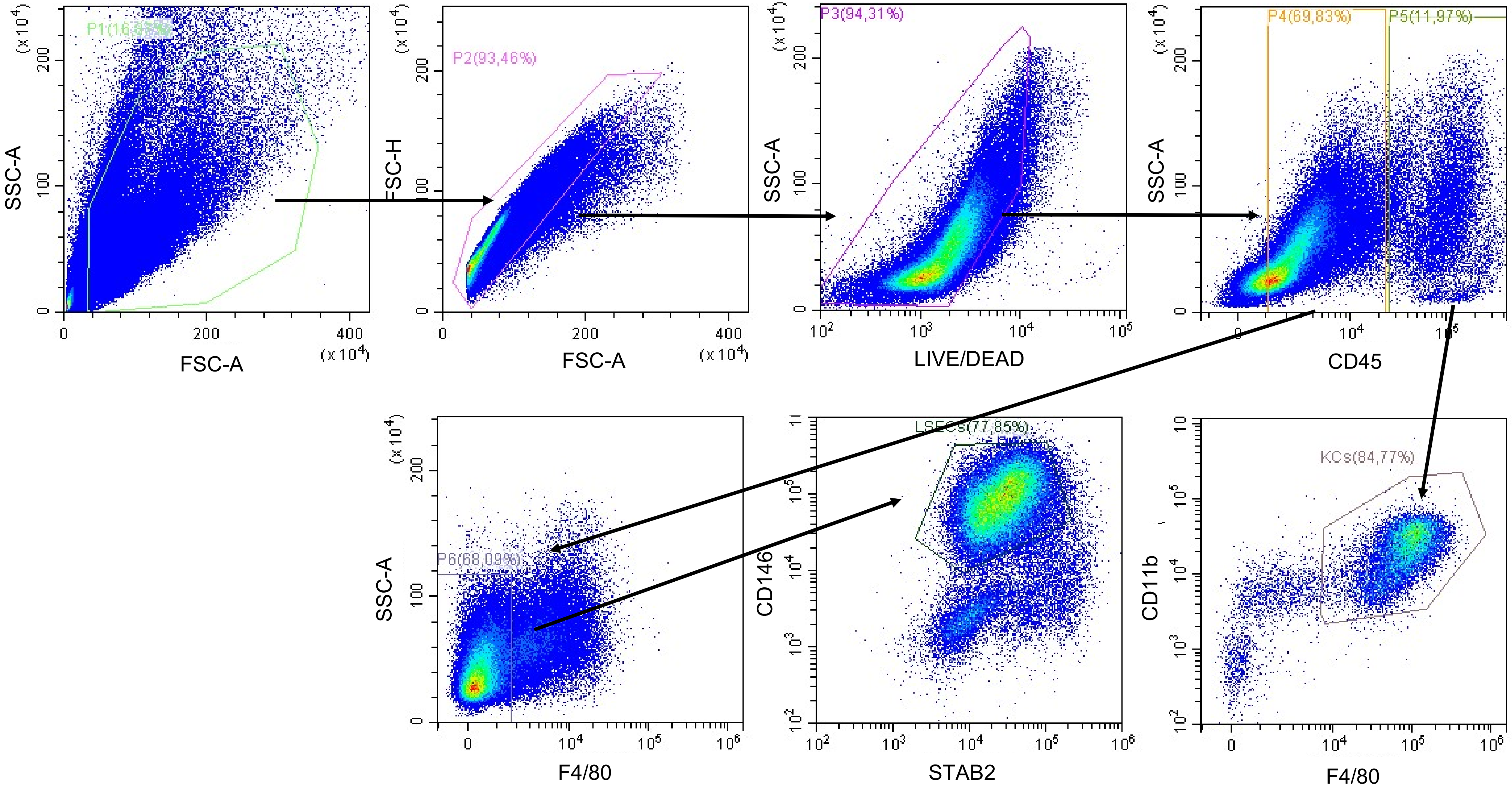

Gating strategy for Figures 1E, 2H (primary human liver NPCs cells)

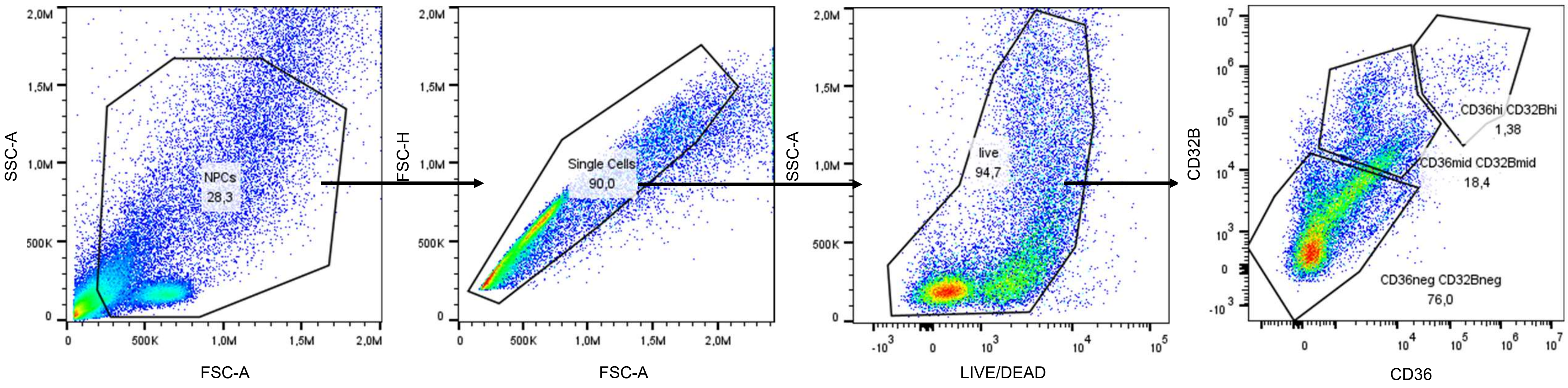

Gating strategy for Figure 2I

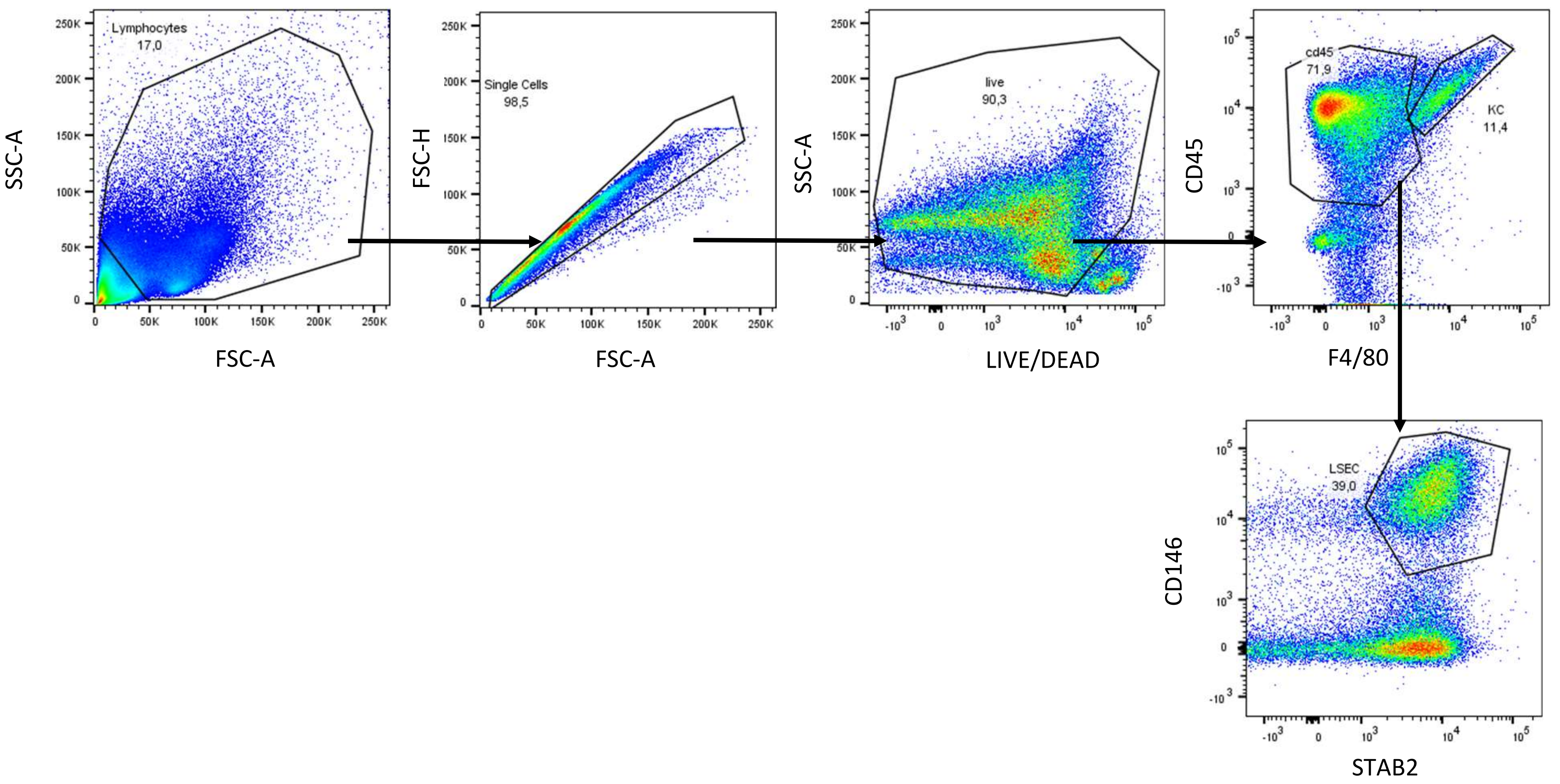

Gating strategy for Figures 3A-D

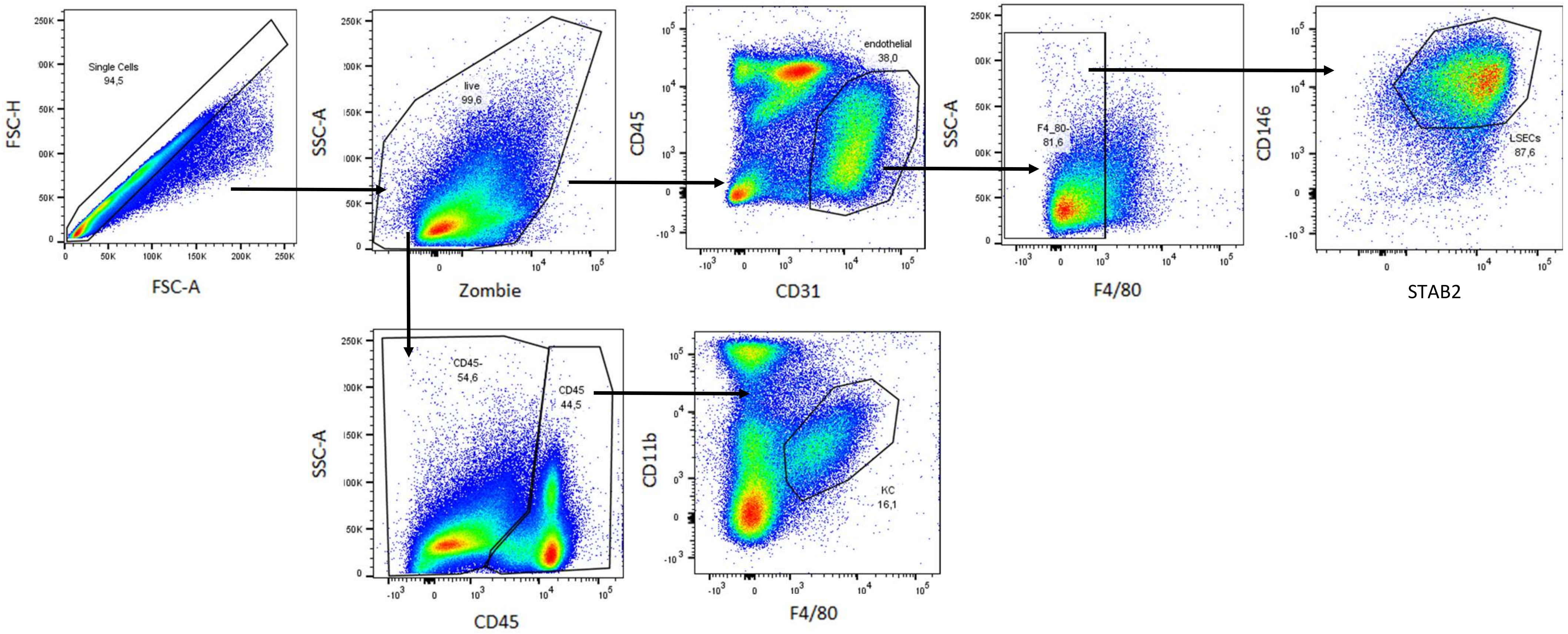

Figures 3E-H

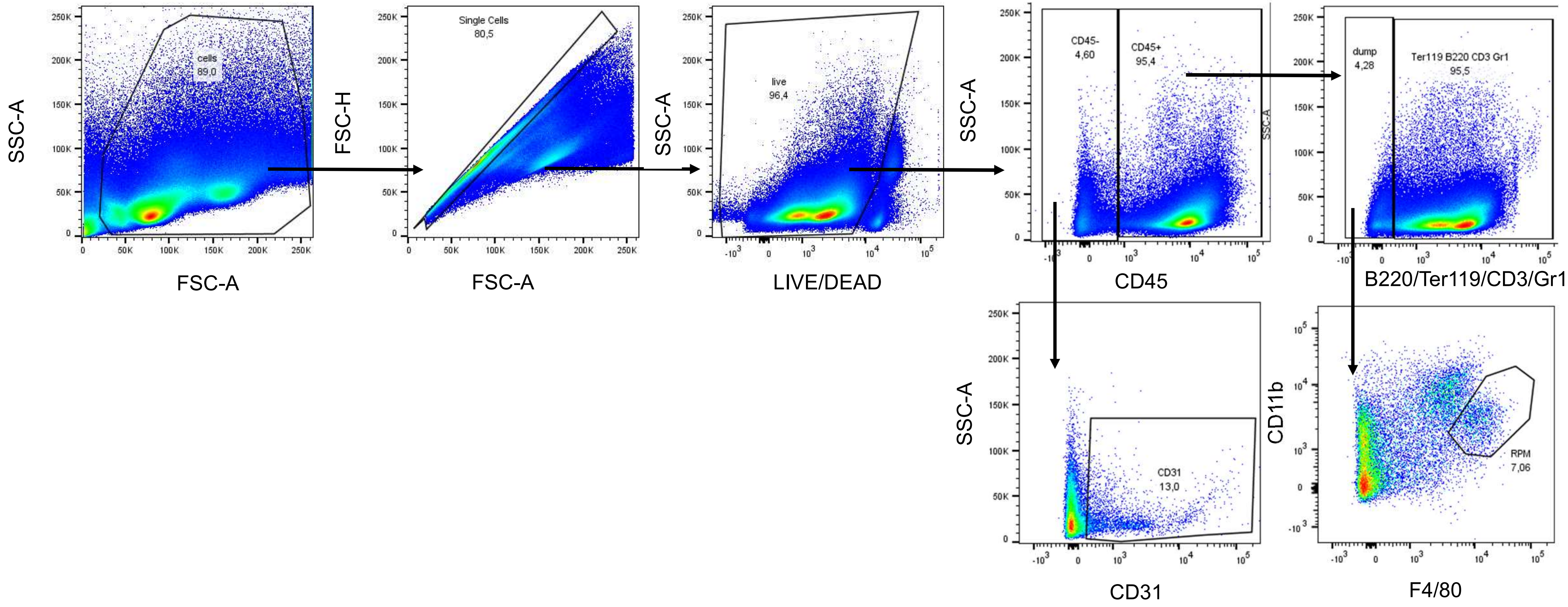

SSC-A
FSC-A
cells
89,0
FSC-H
FSC-A
Single Cells
80,5
SSC-A
LIVE/DEAD
live
96,4
SSC-A
CD45
CD45-
4,60
CD45+
95,4
B220/Ter119/CD3/Gr1
dump
4,28
Ter119 B220 CD3 Gr1
95,5
SSC-A
CD31
CD31
13,0
CD11b
F4/80
RPM
7,06

### Gating strategy for Figures S3A (bone marrow)

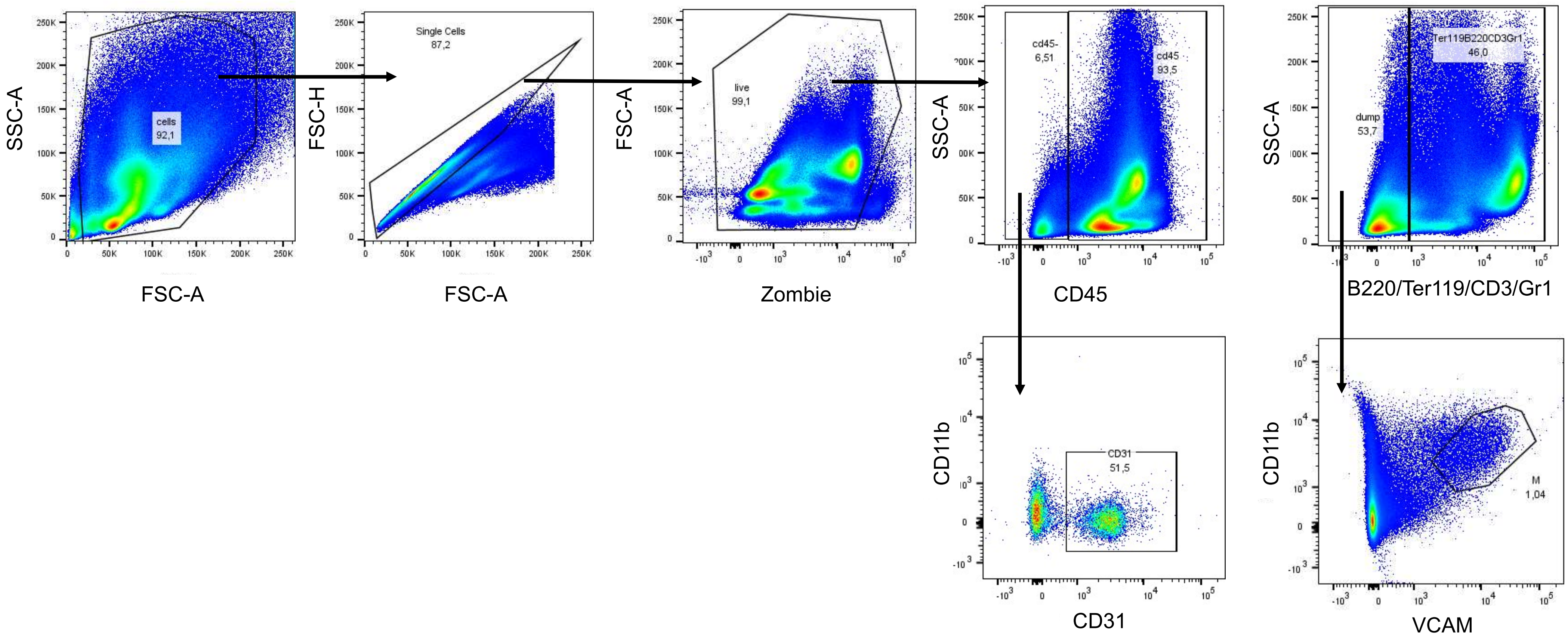

Gating strategy for Figure S3B (aorta)

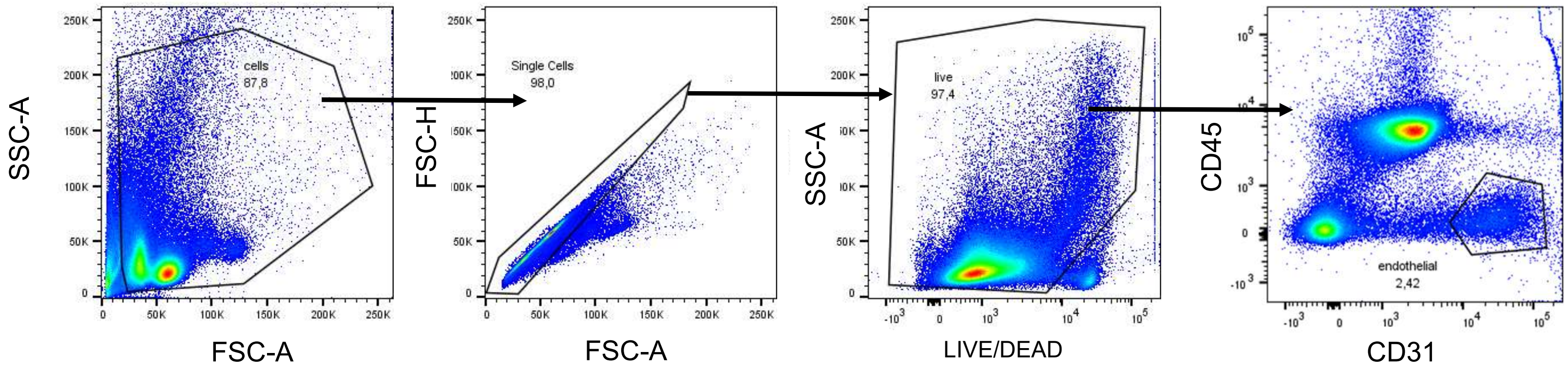

Gating strategy for Figure 4E and 5F – sorting LSECs and KCs

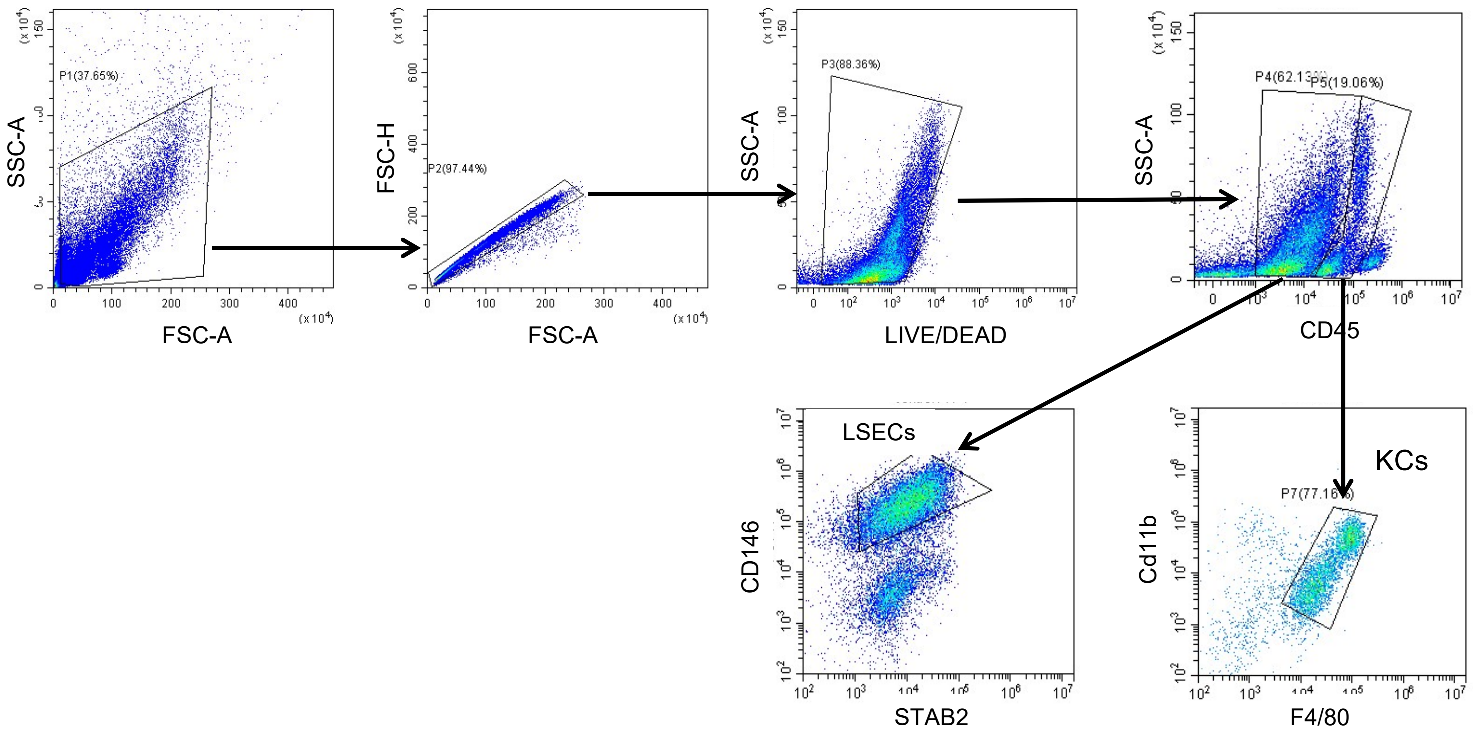

Gating strategy for Figures 4E (sorting spleen endothelial cells)

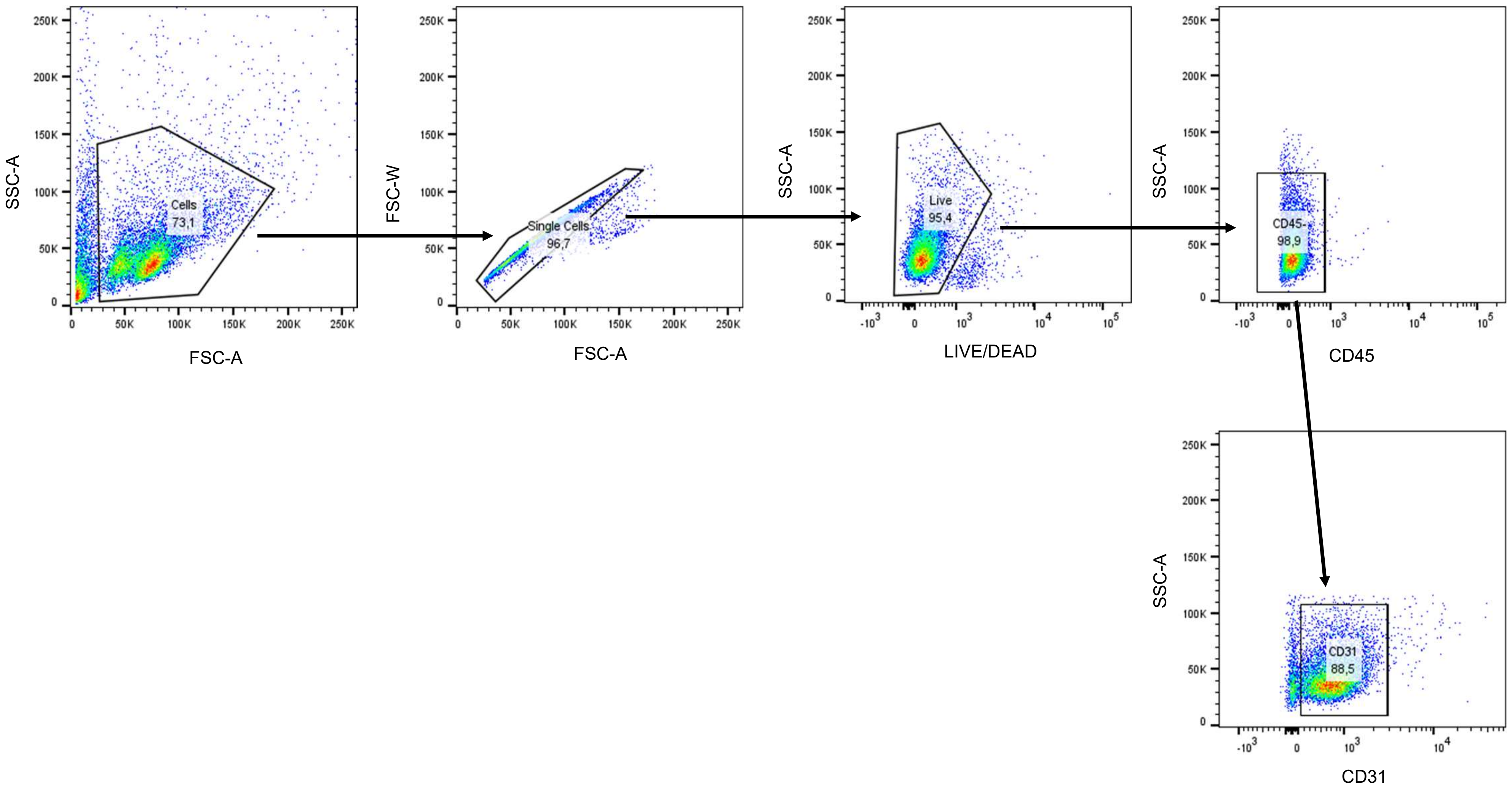

Gating strategy for Figures 4E (sorting heart endothelial cells)

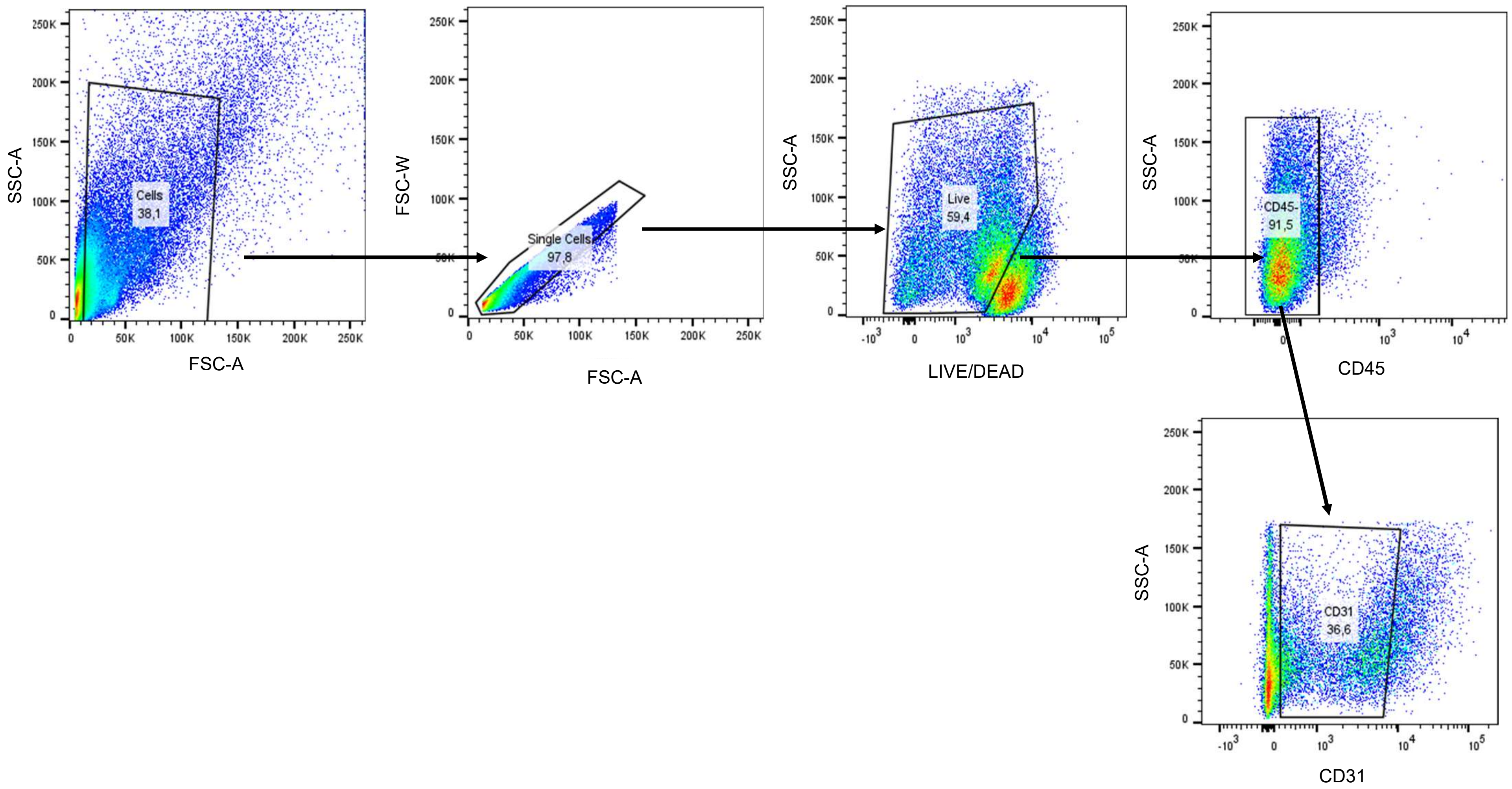

Gating strategy for Figure 4F (liver), 4H and 5J (Ferroorange)

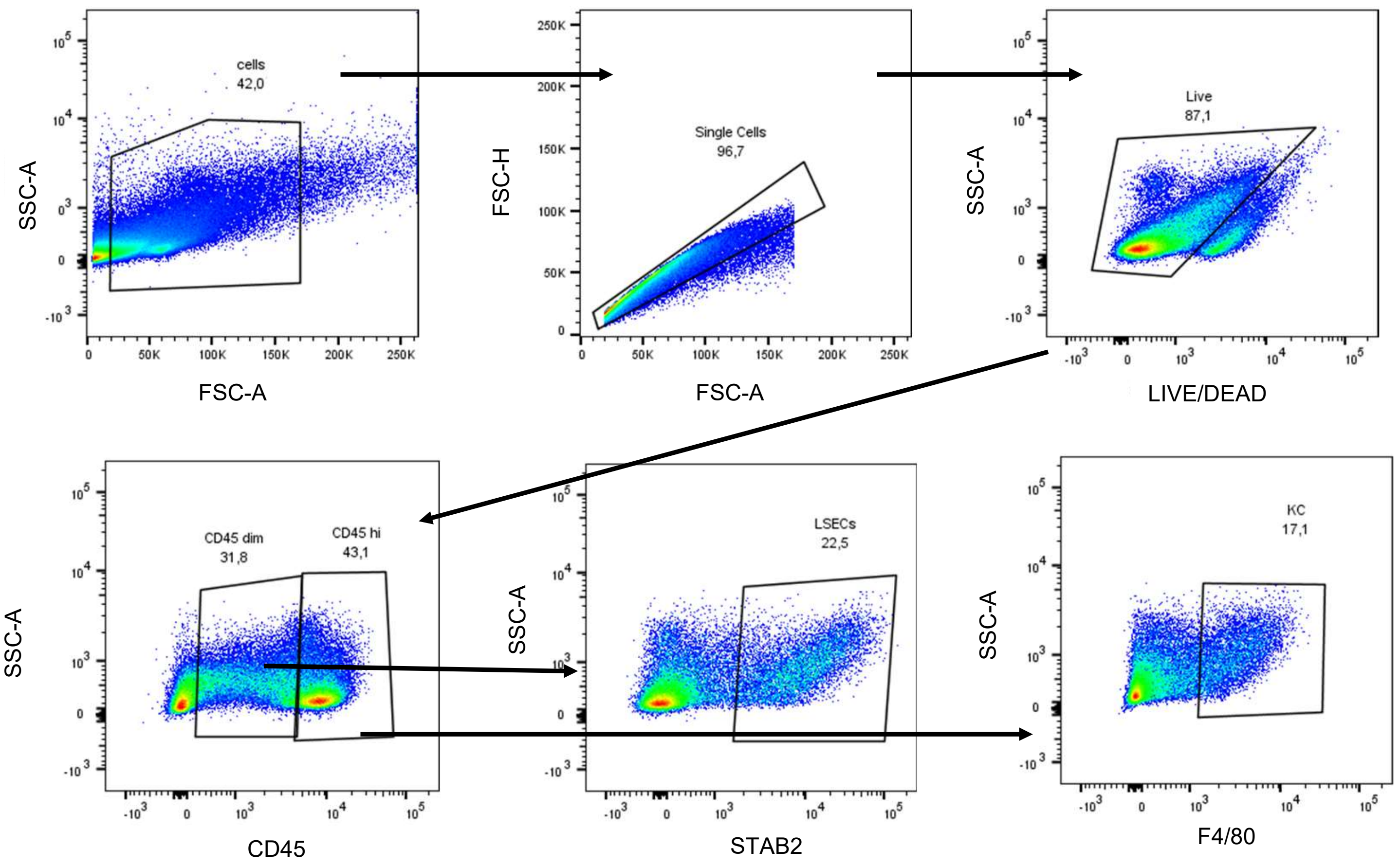

Gating strategy for Figure 4F (spleen)

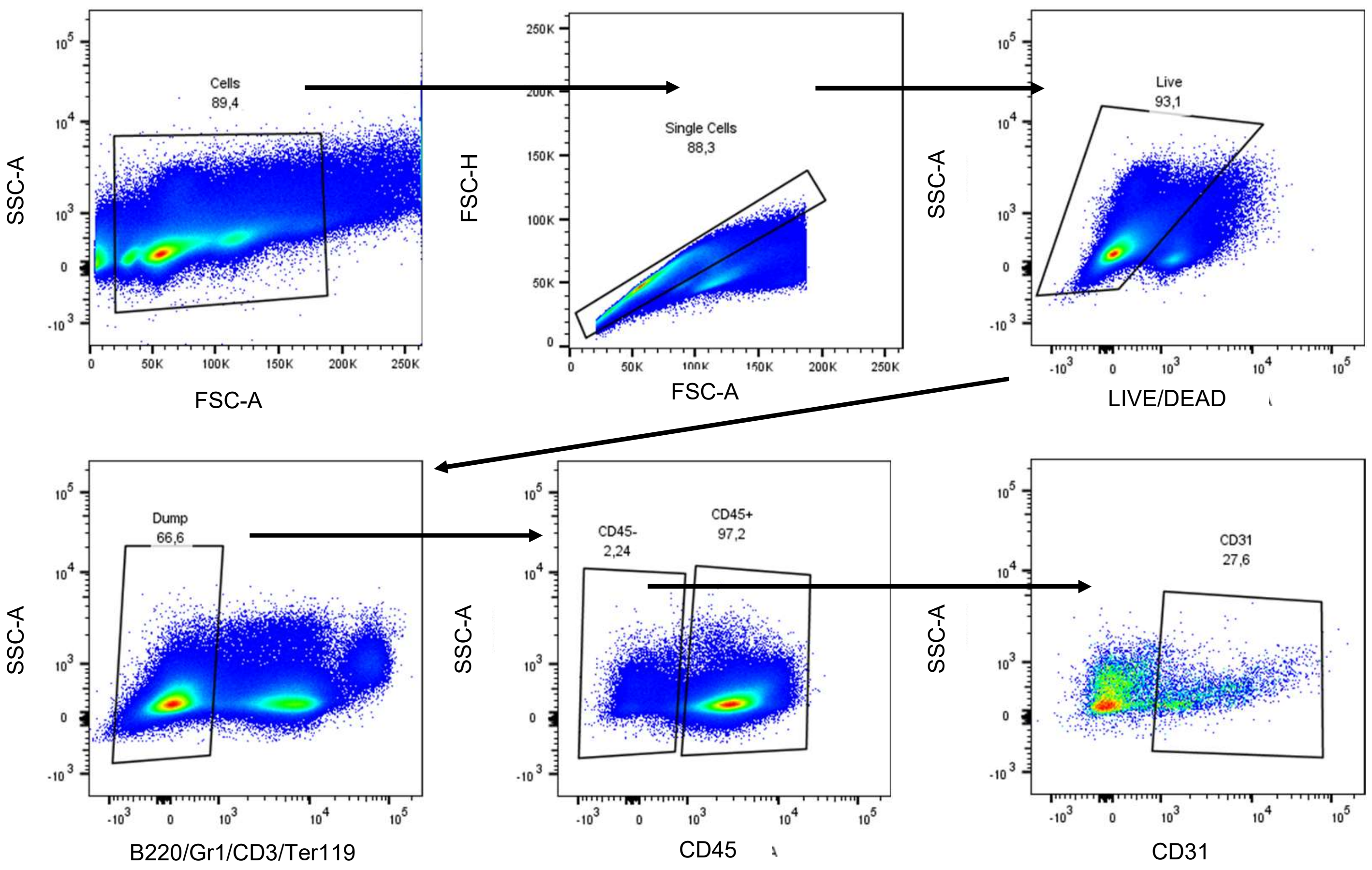

Gating strategy for Figure 4F (bone marrow)

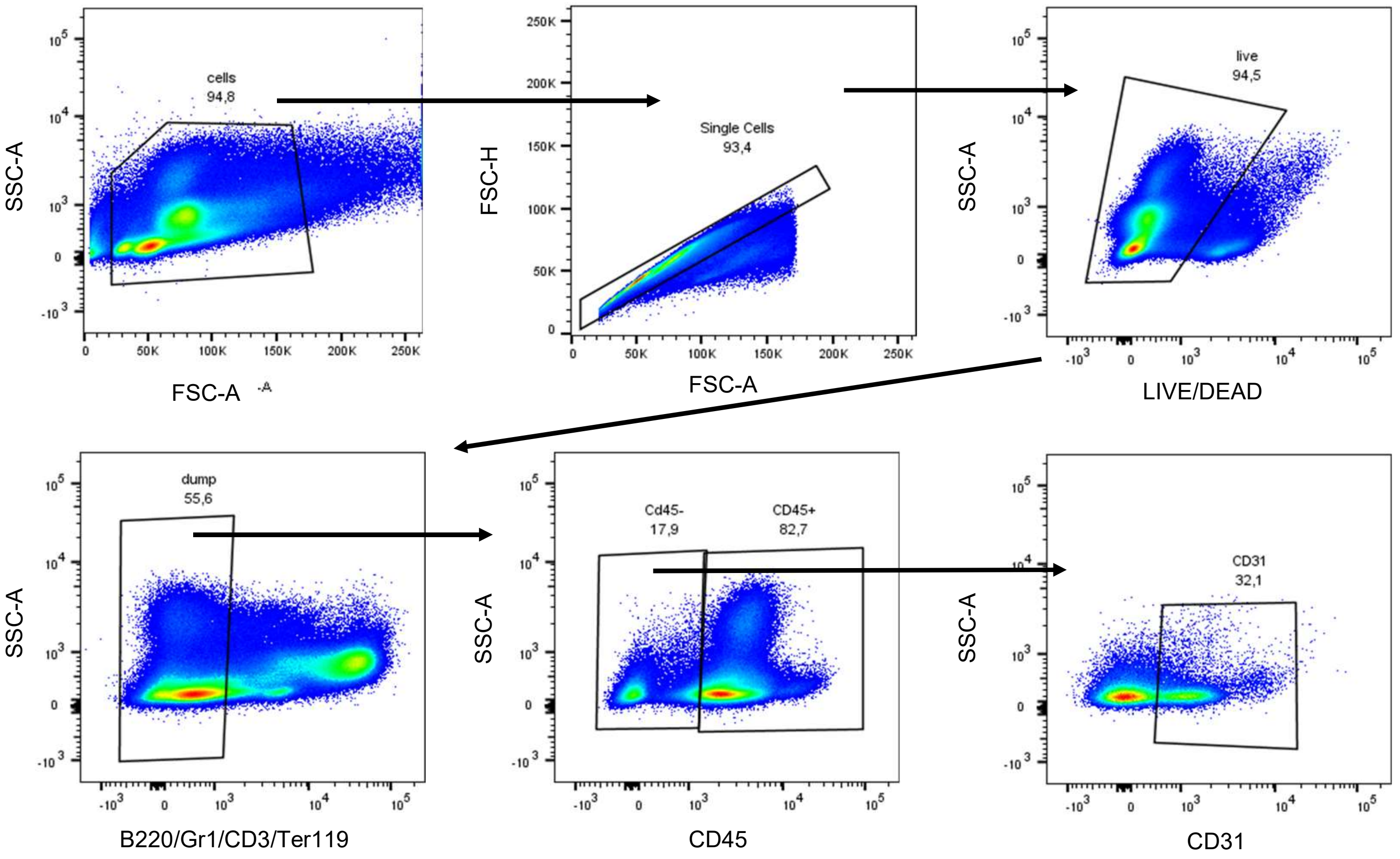

Gating strategy for Figures 4C and D

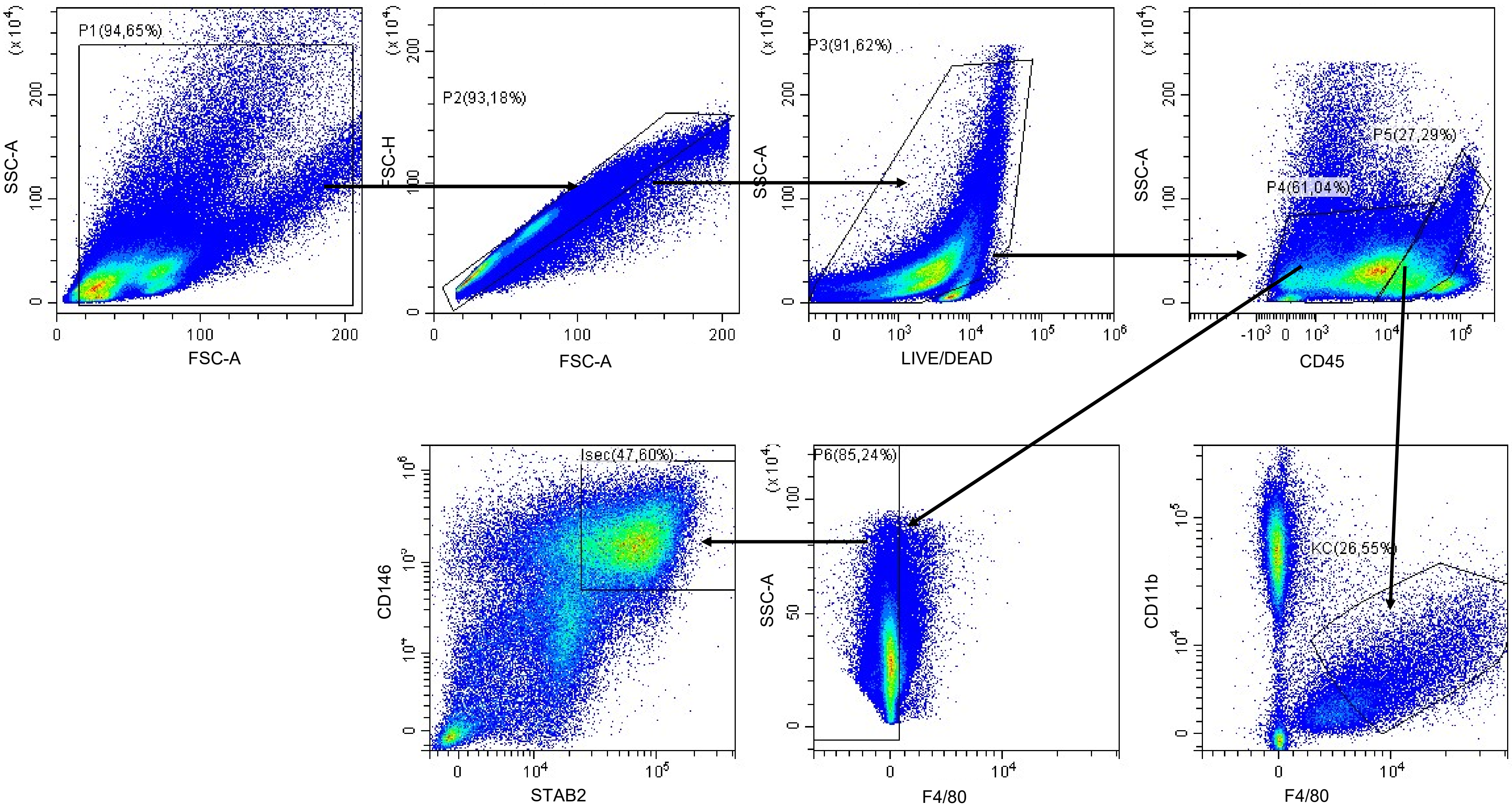

Gating strategy for Figures 4G and Figure 5K

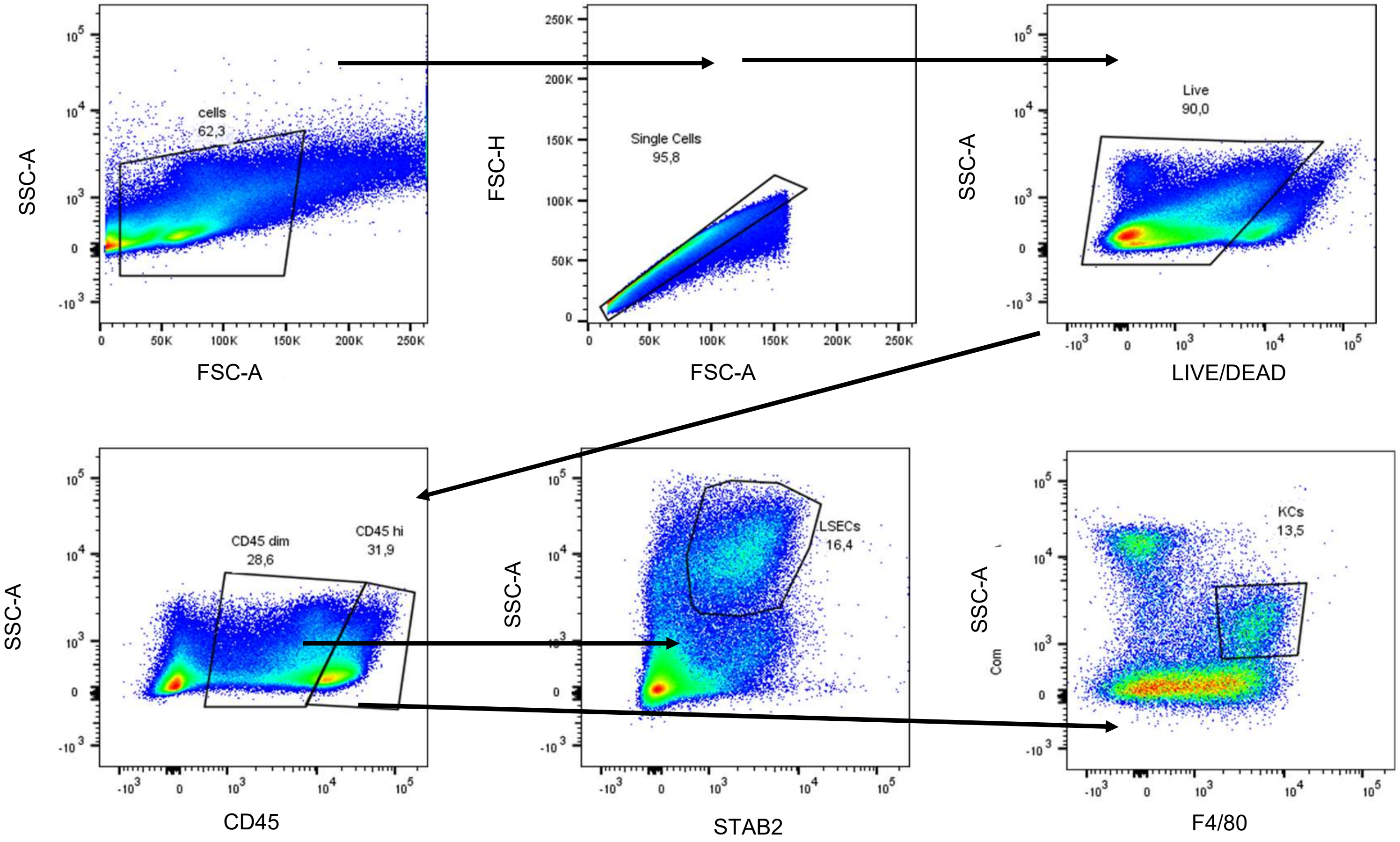

Gating strategy for Figure 5A (spleen control mouse)

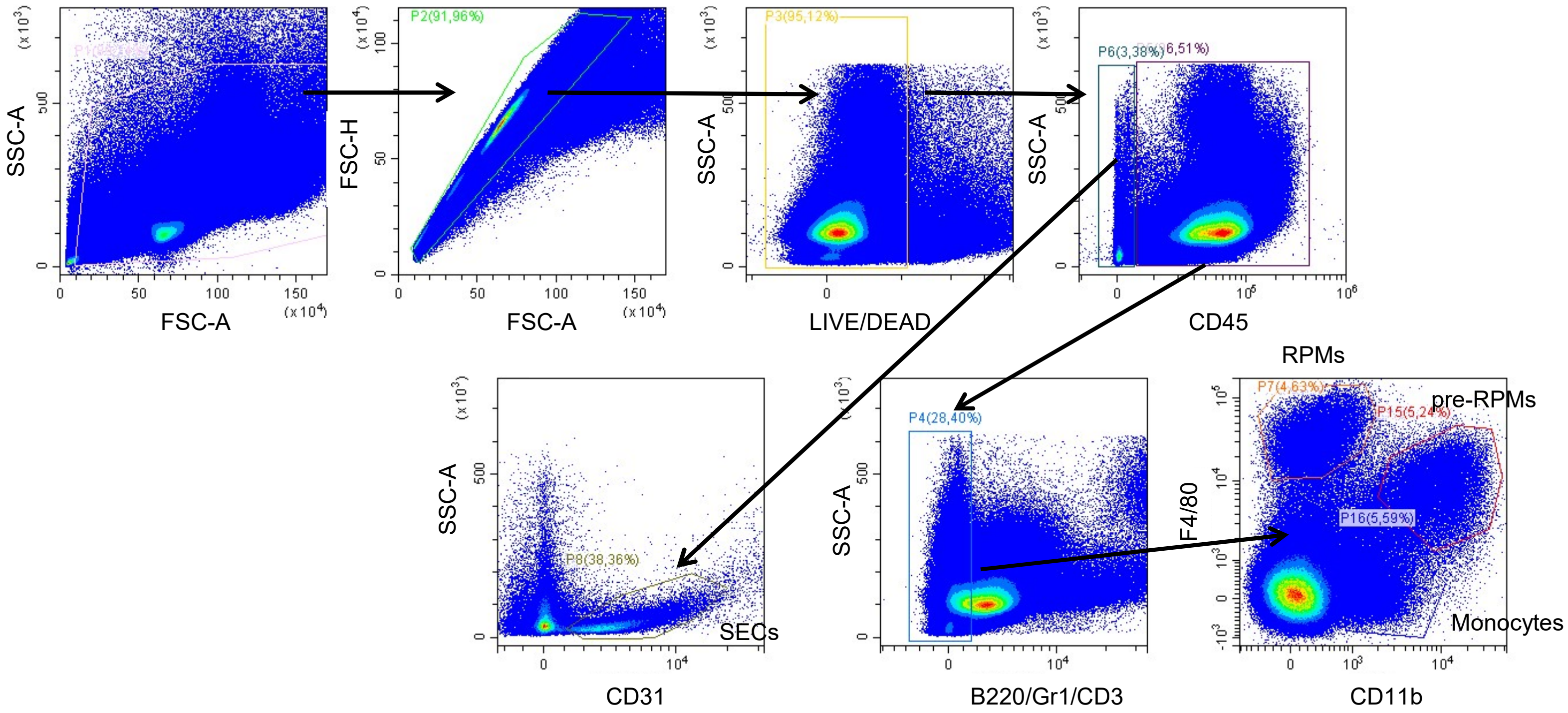

Gating strategy for Figure 5A (spleen + GFP RBCs)

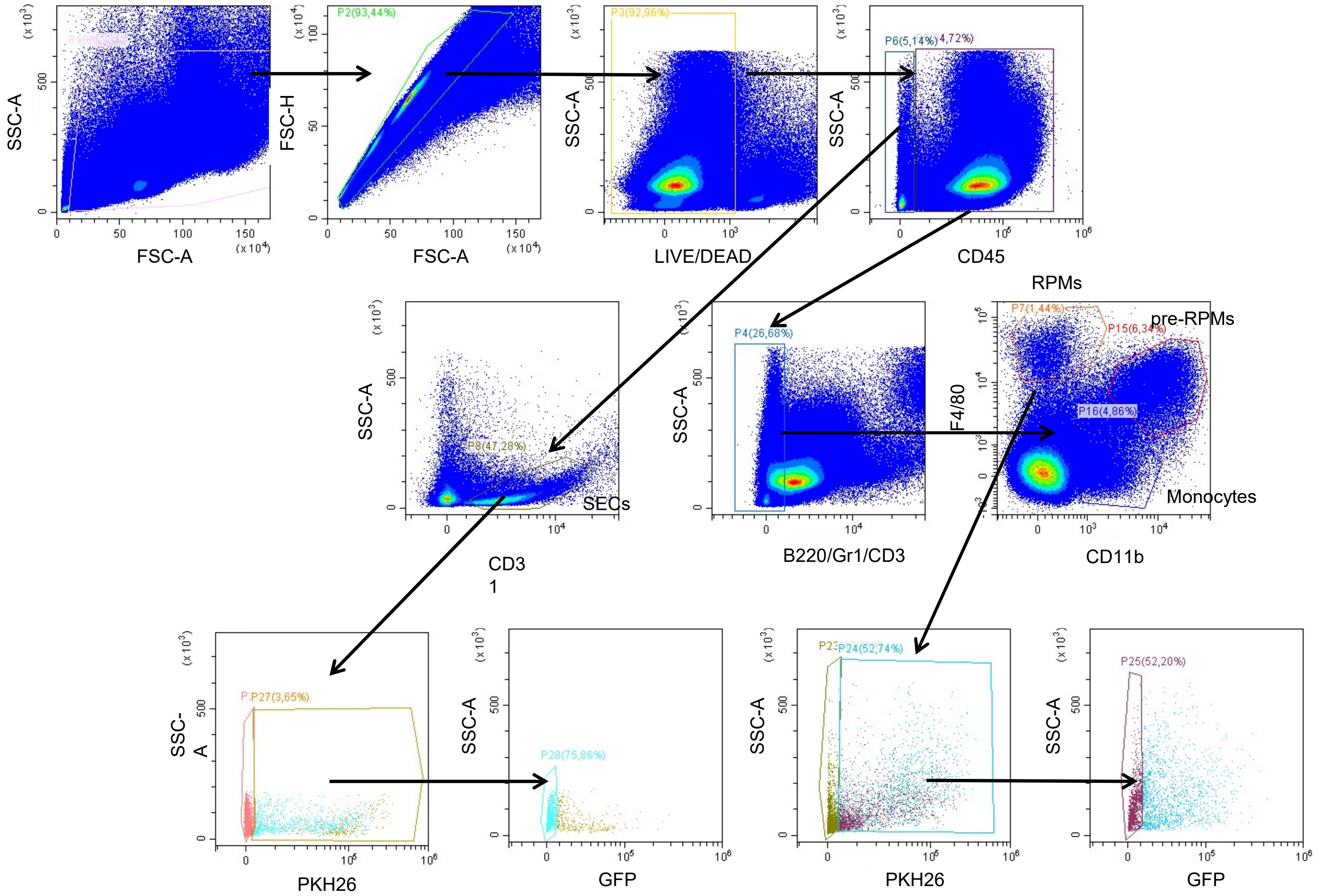

Gating strategy for Figure 5A (spleen dendritic cells)

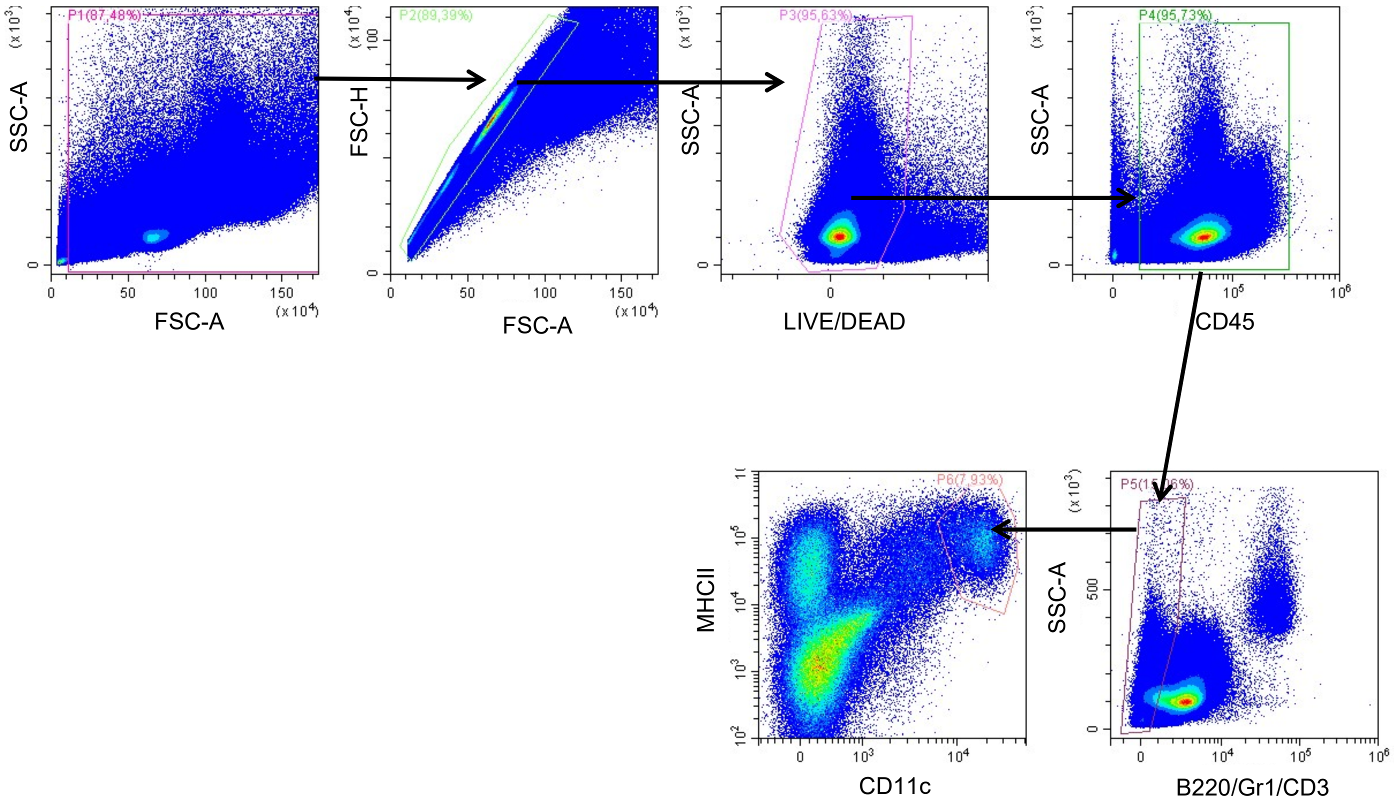

Gating strategy for Figure 5B (liver control mouse)

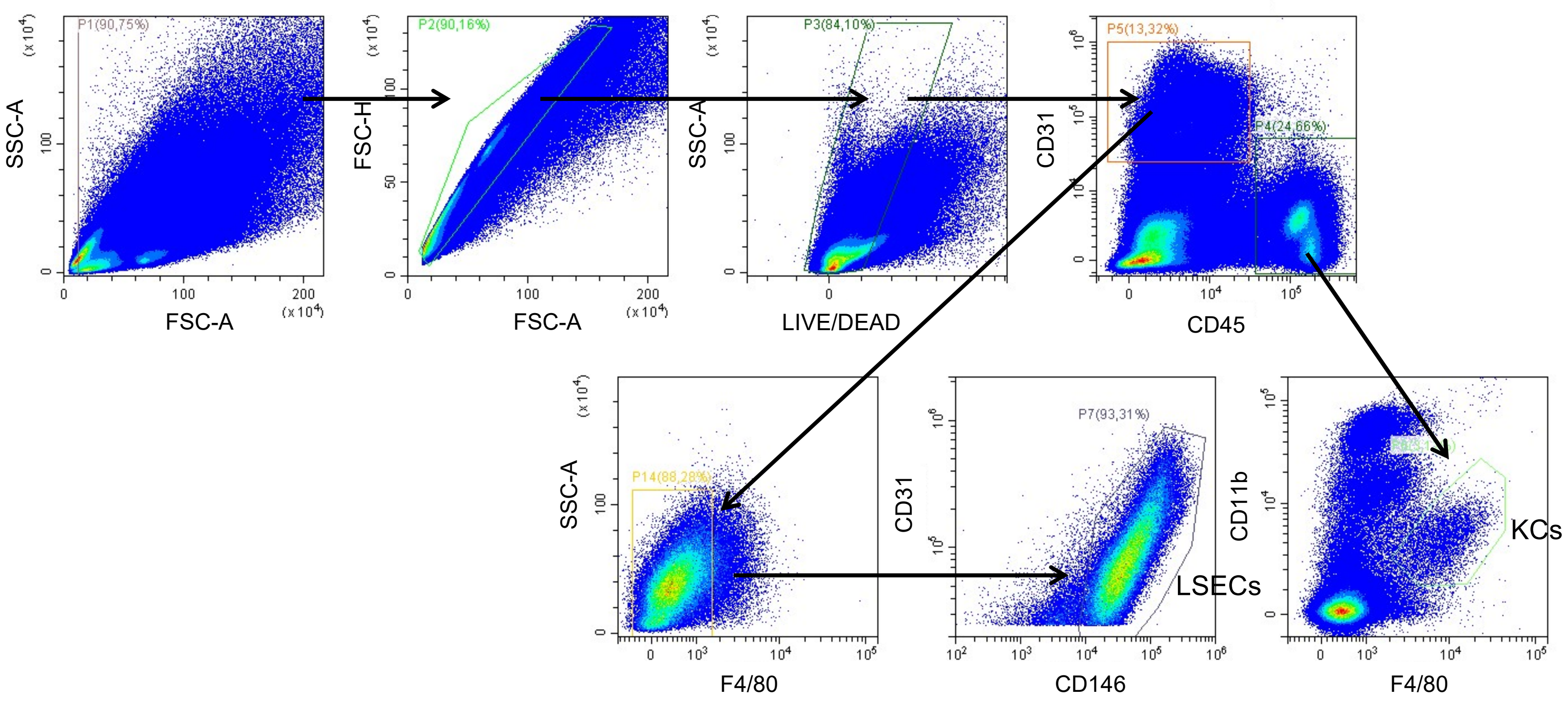

Gating strategy for Figure 5B (liver + GFP RBCs)

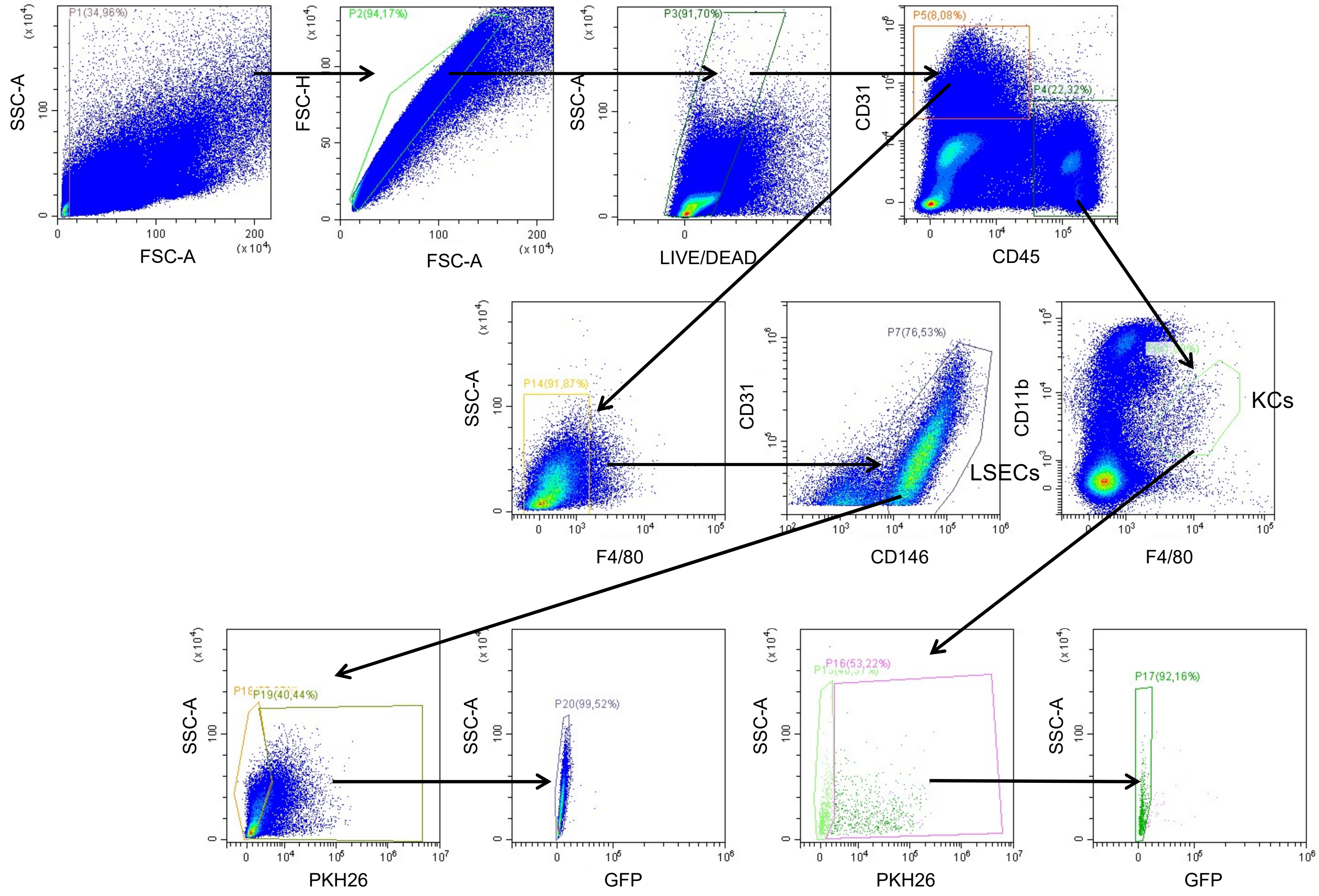

Gating strategy for Figure 5B (liver dendritic cells)

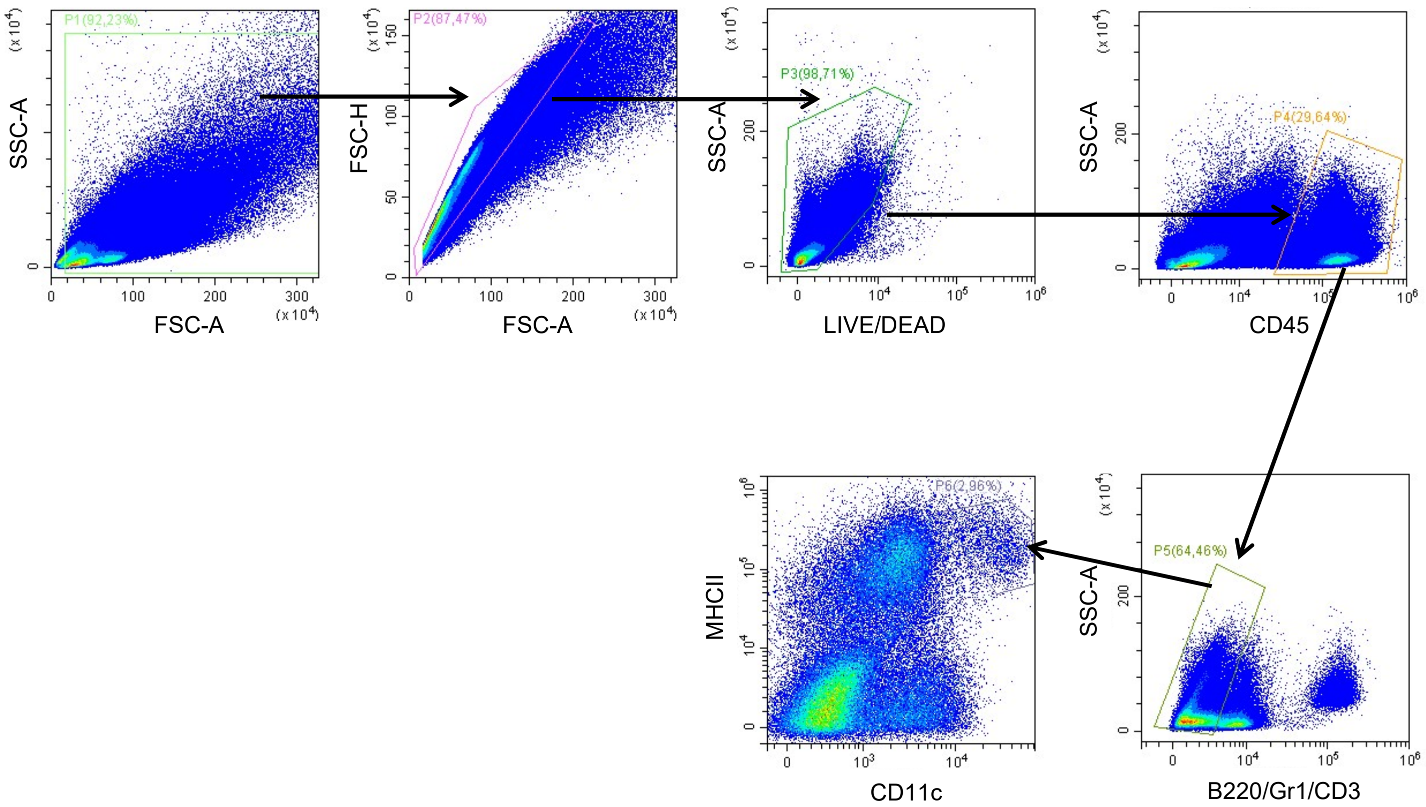

Gating strategy for Figure 5G

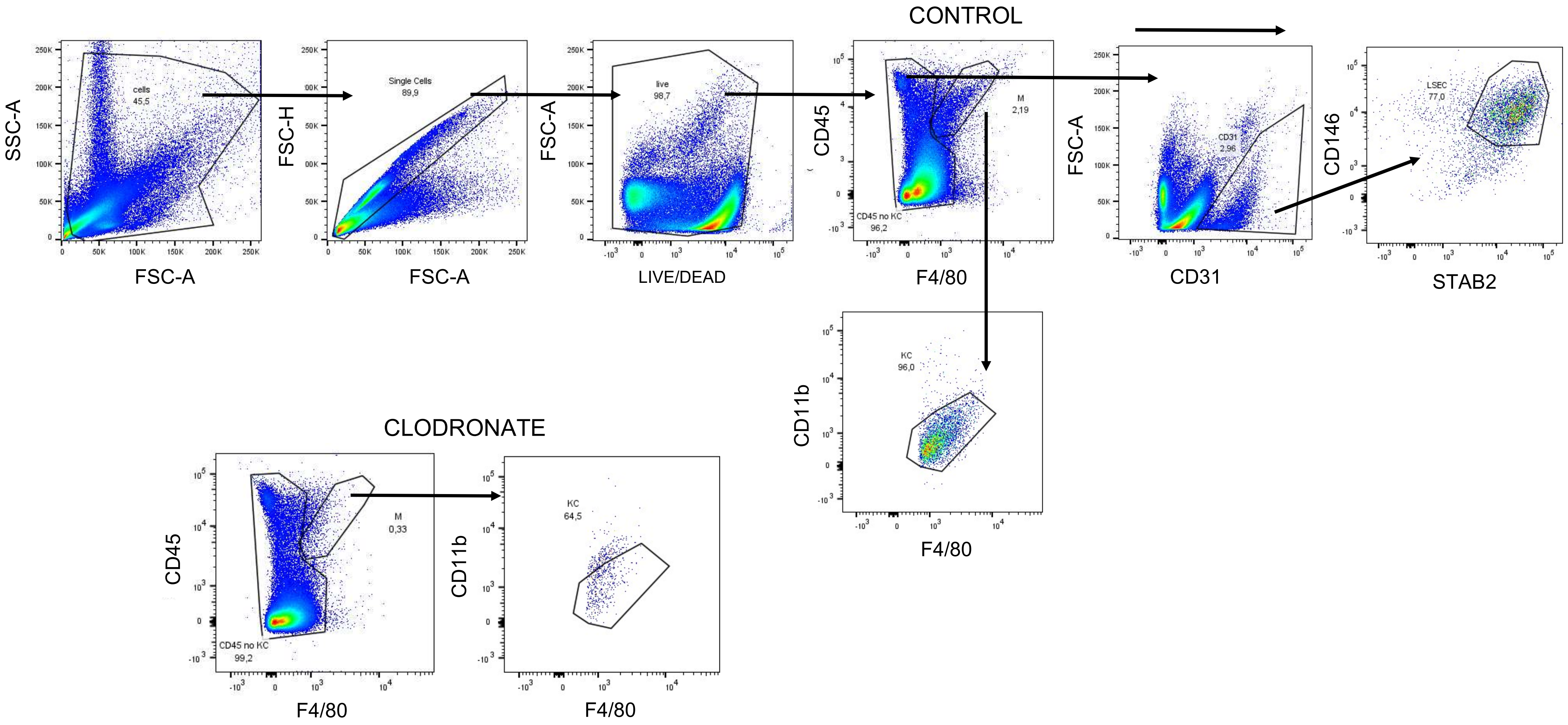

### Gating strategy for Figures 5H and S6B

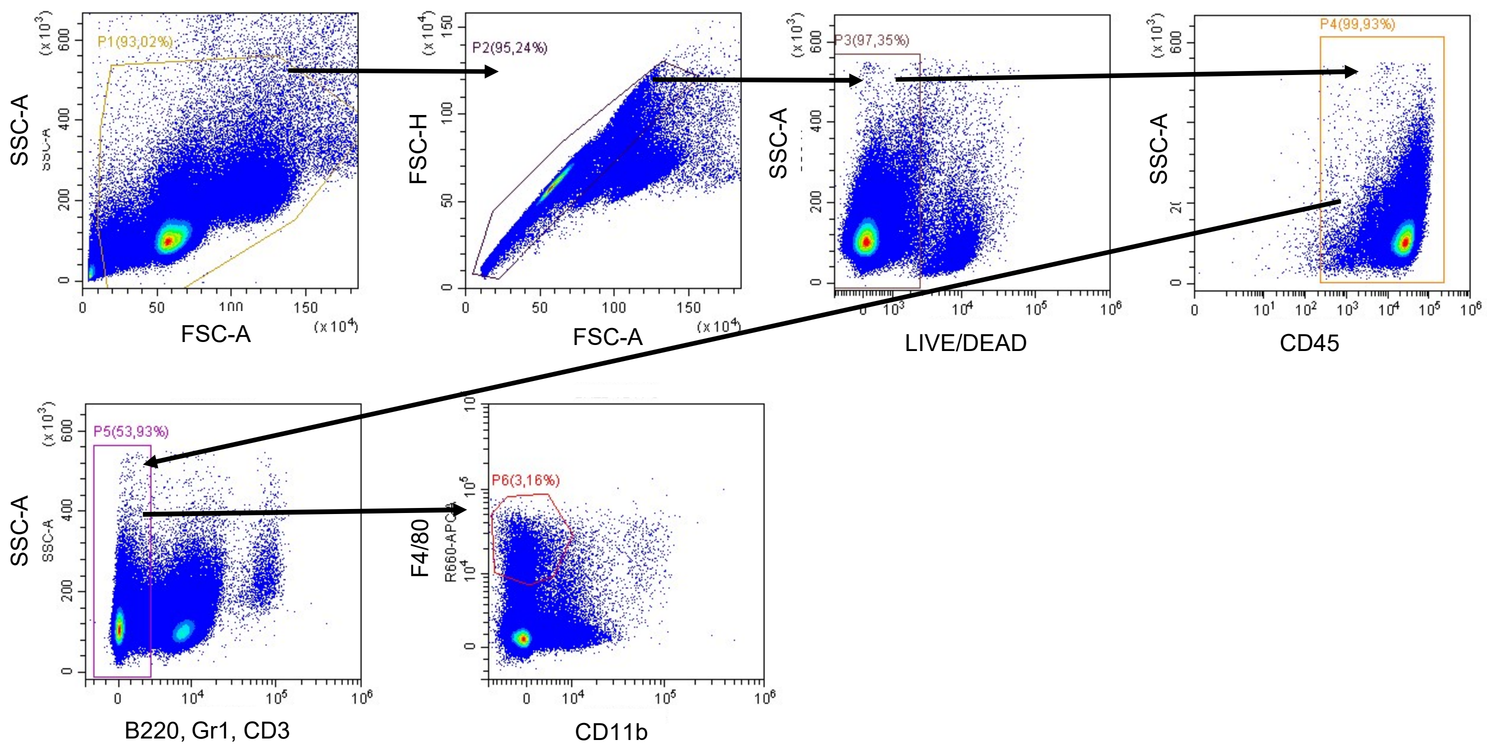

Gating strategy for Figures 6D, 6E, 7B, 7E and 7F, G (liver cell FACS sorting for gene expression)

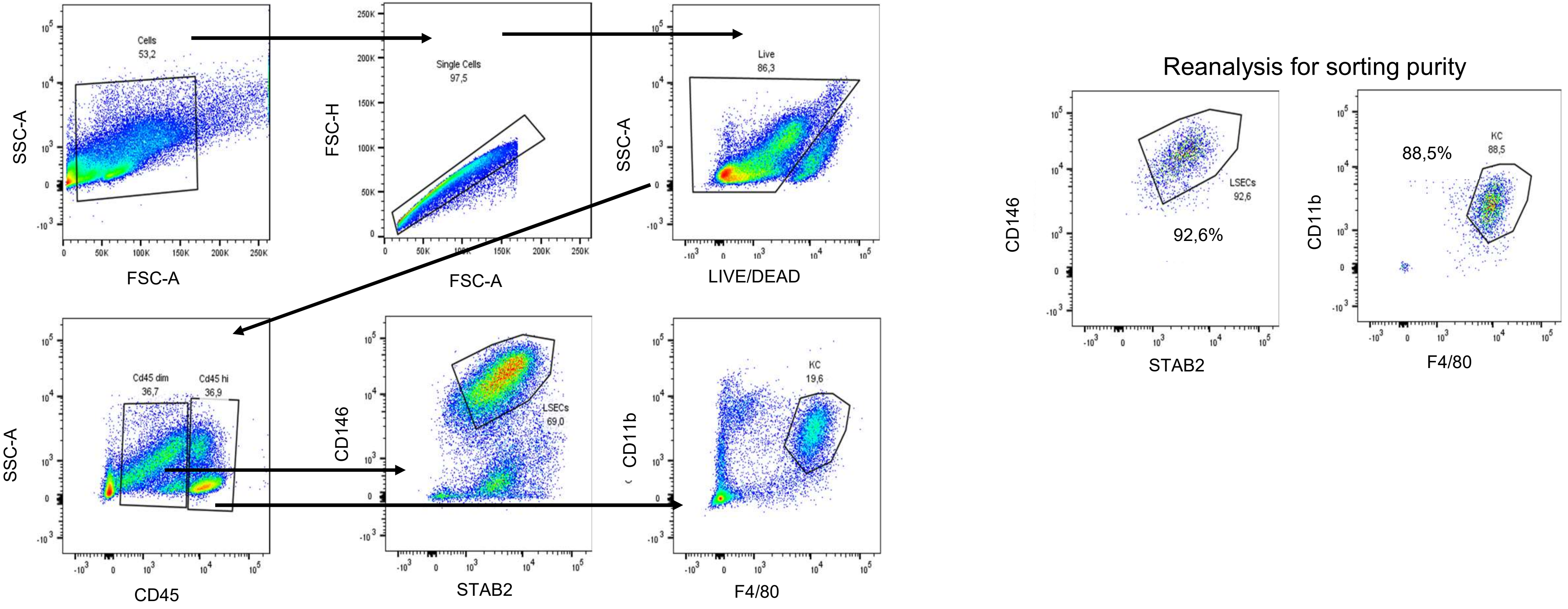
